# Compositional phylogenomic modelling resolves the ‘Zoraptera problem’: Zoraptera are sister to all other polyneopteran insects

**DOI:** 10.1101/2021.09.23.461539

**Authors:** Erik Tihelka, Michael S. Engel, Jesus Lozano-Fernandez, Mattia Giacomelli, Ziwei Yin, Omar Rota-Stabelli, Diying Huang, Davide Pisani, Philip C.J. Donoghue, Chenyang Cai

**Affiliations:** State Key Laboratory of Palaeobiology and Stratigraphy, Nanjing Institute of Geology and Palaeontology, and Center for Excellence in Life and Paleoenvironment, Chinese Academy of Sciences, Nanjing 210008, People’s Republic of China; School of Earth Sciences, University of Bristol, Life Sciences Building, Tyndall Avenue, Bristol BS8 1TQ, United Kingdom; Division of Entomology, Natural History Museum, University of Kansas, Lawrence, Kansas, USA; Department of Ecology & Evolutionary Biology, University of Kansas, Lawrence, Kansas, USA; School of Biological Sciences, University of Bristol, Life Sciences Building, Tyndall Avenue, Bristol BS8 1TQ, United Kingdom; Institute of Evolutionary Biology (CSIC-UPF), Barcelona, Spain; Laboratory of Systematic Entomology, College of Life Sciences, Shanghai Normal University, 100 Guilin Road, Shanghai n/a, China; Research and Innovation Centre, Fondazione Edmund Mach, 38010 San Michele all Adige (TN), Italy; Center Agriculture Food Environment, University of Trento, 38010 San Michele all Adige, Italy

**Keywords:** Zoraptera problem, Pterygota, stem-group fossil, compositional heterogeneity, topological conflict, molecular clock

## Abstract

The evolution of wings propelled insects to their present mega-diversity. However, interordinal relationships of early-diverging winged insects and the timescale of their evolution are difficult to resolve, in part due to uncertainties in the placement of the enigmatic and species-poor order Zoraptera. The ‘Zoraptera problem’ has remained a contentious issue in insect evolution since its discovery more than a century ago. This is a key issue because different placements of Zoraptera imply dramatically different scenarios of diversification and character evolution among polyneopteran. Here, we investigate the systematic placement of Zoraptera using the largest protein-coding gene dataset available to date, deploying methods to mitigate common sources of error in phylogenomic inference, and testing historically proposed hypotheses of zorapteran evolution. We recover Zoraptera as the earliest-diverging polyneopteran order, while earwigs (Dermaptera) and stoneflies (Plecoptera) form a monophyletic clade (Dermoplectopterida) sister to the remainder of Polyneoptera. The morphology and palaeobiology of stem-zorapterans are informed by Mesozoic fossils. The gut content and mouthparts of a male specimen of *Zorotypus nascimbenei* from Kachin amber (Cretaceous) reveal a fungivorous diet of Mesozoic zorapterans, akin to extant species. Based on a set of 42 justified fossil and stratigraphic calibrations, we recover a Devonian origin of winged insects and Polyneoptera, suggesting that these groups coincided with the rise of arborescence during the diversification of early terrestrial plants, fungi, and animals. Our results provide a robust framework for understanding the pattern and timescale of early winged insect diversification.

## Introduction

The ability to fly paved the way for the winged insects (Pterygota) to become the most diverse and abundant animal group, and today pterygote insects account for over 98% of hexapod species richness^1^. However, the interordinal relationships within Polyneoptera (an ancient and diverse clade of winged insects with a fossil record extending to the Carboniferous) remain unclear^2–7^. Polyneoptera includes the first great diversification of neopteran insects, those winged insects with the novel ability to fold their wings over the body at rest, thereby protecting wings when not in use and also allowing wings to take on many novel secondary specializations^1^. Accordingly, Polyneoptera constitutes the first major proliferation of insect lineages that remain significant components of the insect fauna to this day. The evolutionary roots of the eleven extant polyneopteran orders (earwigs, grasshoppers, mantises, roaches, stick insects, and allies) have proven difficult to resolve ^6,8–10^. Uncertainties in polyneopteran relationships have implications for our understanding of the early diversification of neopteran winged insects since reconstructing the morphology and biology of the ancestral polyneopteran is contingent on resolving the early divergences in the clade. The timescale of polyneopteran evolution has likewise remained difficult to constrain, with molecular clock analyses of different topologies yielding a wide range of dates spanning the Early Devonian to the Middle Triassic^4,7,11–15^.

Crucial to disentangling the early evolution of Polyneoptera is the phylogenetic position of the enigmatic order Zoraptera, sometimes referred to as ‘groundlice’ or ‘angel insects’. With less than 50 described extant species, zorapterans are the third smallest and arguably the least understood insect order^16^. Since their discovery in 1913^17^, the phylogenetic position of zorapterans has represented one of the most persistent problems in reconstructing the insect tree of life. They have been placed in a bewildering number of positions, including as sister to Acercaria (Hemiptera and allies), all Holometabola (insects that undergo full metamorphosis), Dictyoptera (Mantodea + Blattodea), Eukinolabia (Phasmatodea + Embioptera), earwigs (Dermaptera), and webspinners (Embiodea) ^18–25^. Over the last decade, phylogenetic analyses of morphological and molecular data have unequivocally supported the placement of zorapterans within Polyneoptera, but systematic affinities with almost all polyneopteran clades have been suggested ^2,4,16,21,26–28^, leading Beutel and Weide ^21^ to dub this the ‘Zoraptera problem’, reflecting its resistance to resolution. The Zoraptera problem results, at least in part, from the insects’ unusual biology and associated morphological specializations; the minute adults occur as either fully-sighted (i.e., well-developed compound eyes) alate individuals or blind and wingless morphs that live in colonies inside decaying logs or termite nests. Zorapterans are primarily distributed in the pantropics, although some species extend northwards in North America and Asia. Aside from their unusual morphology and high rates of molecular evolution^18^, incongruences in resolving the systematic placement of zorapterans are exacerbated by their scant fossil record, which discloses little about their affinities^29^.

To investigate the phylogenetic position of Zoraptera, we reanalysed the most comprehensive genome-scale dataset available^2^ including representatives of all major clades of winged insects and all representatives of extant polyneopterans, and we used it to test alternative hypotheses of zorapteran evolution. To understand why past studies reached contrasting results for the relationships of the major orders, we investigated the effects of common sources of systematic error in phylogenomic inference through dataset curation and use of models that can account for site-specific compositional heterogeneity^30^. Complementing our transcriptome analyses, we discuss the morphology of fossil zorapterans from the mid-Cretaceous amber from northern Myanmar. An exquisitely preserved specimen of *Zorotypus nascimbenei* reveals that a fungivorous diet was already present among stem-zorapterans. Finally, we integrate transcriptomic and palaeontological data to elucidate the timescale of polyneopteran evolution and propose a revised classification of Zoraptera.

## Results

### Phylogenomics confidently assesses the placement of Zoraptera

Resolution of conflicting hypotheses on the early evolution of Polyneoptera has to account for common sources of error in phylogenetic analyses and demonstrate why past analyses arrived at divergent topologies^31^. Tree reconstruction artifacts are frequently recovered when models are used that make incorrect assumptions about the data^30,32^. To explore the effect of different substitution models on the recovered phylogeny of Polyneoptera, we tested models of increasing complexity^33^: (i) standard site-homogeneous amino acid substitution models LG+G and GTR+G, variations of which have been used in previous studies^2,4^, (ii) the multi-matrix mixture model LG4X, and (iii) the compositionally site-heterogeneous infinite mixture model CAT-GTR+G and the compositionally site-heterogeneous LG+C60+F+G model. Among-site heterogeneous mixture models such as CAT-GTR+G have been shown to offer improved resilience to long-branch attraction artefacts (LBA)^34^ and invariably provides better fit to the data than standard site-homogeneous models^31,35–39^. In our cross-validation analysis, CAT-GTR+G fitted the dataset better than the site-homogeneous LG+G and GTR+G models (CAT-GTR+G > LG+G: CV score = 34,548.8 ± 344.907; CAT-GTR+G > GTR+G: CV score = 29,639.7 ± 293.483). We thus treat the CAT-GTR+G tree as the most trustworthy topology (Fig. 1).

**Figure 1.**
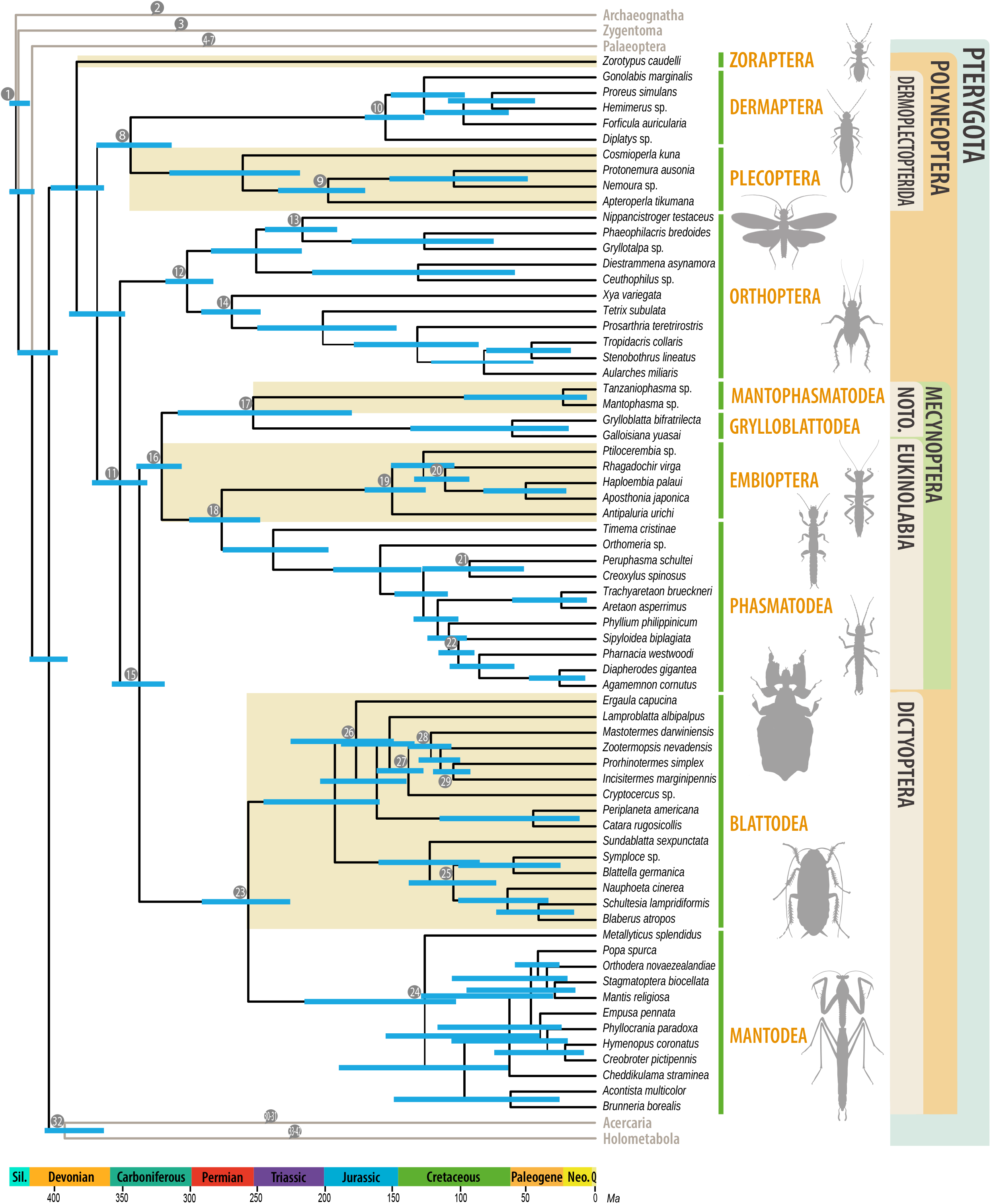
Timescale of Polyneoptera evolution from analyses with the site-heterogeneous model CAT-GTR+G of the 100-taxon dataset. All nodes are fully supported (BPP = 1). Ages were estimated based on 42 calibrated nodes, integrating the results of analyses using independent rates (IR) and autocorrelated rates (AC) molecular clock models in MCMCTree. Abbreviations: Noto., Notoptera; Sil., Silurian; Q., Quaternary. Numbered nodes indicate the calibrations, see Supplementary Information for full list of calibrations and Fig. S1 for a full calibrated tree.

Analyses conducted using the site-homogeneous models LG+G, GTR+G, and the LG4X model, recovered a monophyletic Polyneoptera with a Zoraptera + Dermaptera clade (bootstrap value = 63–98) as sister to the remaining orders (Figs S16–23). As expected, these results mirror those obtained by recent transcriptomic-based phylogenies using site-homogeneous models^2,4^, and highlight the importance of testing the effect of compositionally-heterogeneous models to ensure a complete exploration of the data.

Our CAT-GTR+G and LG+C60+F+G analyses recovered a monophyletic Polyneoptera with maximum support (Bayesian posterior probabilities [BPP] = 1; bootstrap values = 100), in congruence with recent phylotranscriptomic studies^2,4,6^. Our results however differ from recent phylogenomic analyses in the placement of the early-diverging orders. Zoraptera was recovered as sister to the remaining orders with full support (BPP = 1; bootstrap = 100). Dermaptera and Plecoptera formed a well-supported clade (BPP = 1; bootstrap 90–97), sister to the remainder of Polyneoptera, excluding Zoraptera. The relationships among the remaining polyneopteran orders were in line with previous transcriptomic phylogenies. Orthoptera was sister to the remainder of Polyneoptera (excluding Zoraptera, Dermaptera, and Plecoptera), Mantophasmatodea + Grylloblattodea (Notoptera) formed a clade sister to Phasmatodea + Embioptera (Eukinolabia) dubbed Mecynoptera, and Mantodea were sister to Blattodea (Dictyoptera). The position of Zoraptera remained stable across the four analysed datasets with different taxon sampling adjusted to exclude rogue taxa and to the effect of just using a single species per order. However, analyses of a 33-taxon dataset with CAT-GTR+G, sampling only one representative of each polyneopteran order, yielded an identical position of Zoraptera, but recovered Orthoptera as sister to Plecoptera, instead of as sister to the remaining polyneopterans as in analyses with denser taxon sampling. Analyses of the 33-taxon dataset with the LG+C60+F+G model recovered Zoraptera as sister to Dermaptera with low support (bootstrap = 85). Outside Polyneoptera, the monophyly of Palaeoptera and Acercaria, supported by traditional morphology-based classification^20^ but questioned by some recent analyses^4^, was recovered with full support (although support for Acercaria was low in analyses of the 102-taxon matrix under both models and the 106-taxon dataset under the LG+C60+F+G model; Figs S5, 8, 9).

### Evaluating alternative hypotheses of polyneopteran evolution

To compare support for alternative hypotheses of early polyneopteran evolution, we ran topology tests in IQ-TREE using the site-heterogeneous LG+C60+F+G model. Model selection suggested CAT-GTR+G to be the best fitting model. However, CAT-GTR+G is not implanted in IQ-TREE. Accordingly, to run our topology tests we used LG+C60+F+G, which despite not the best fitting model, can still account for compositional heterogeneity in a maximum likelihood framework. Congruently with the majority of recent morphological and morphological studies^2,4,22,40^, the Approximately Unbiased (AU) test rejected placements of Zoraptera outside of Polyneoptera. In addition, AU tests rejected the placement of Zoraptera outside of ‘basal Polyneoptera’, as well as sister to Plecoptera, sister to Polyneoptera excluding Dermaptera, or sister to a clade including Dermaptera and Plecoptera (*P*_AU_ = 0; Fig. 2, Tab. S1). The Haplocercata first topology favoured by analyses with site-homogeneous models ^2,4^, with Zoraptera as sister to Dermaptera, was also rejected (*P*_AU_ > 0.006; Fig. 2B). Out of 20 tested topologies, only the ‘Zoraptera-first’ topology recovered in our CAT-GTR+G analyses was not rejected (*P*_AU_ > 0.995; Fig. 2A).

**Figure 2.**
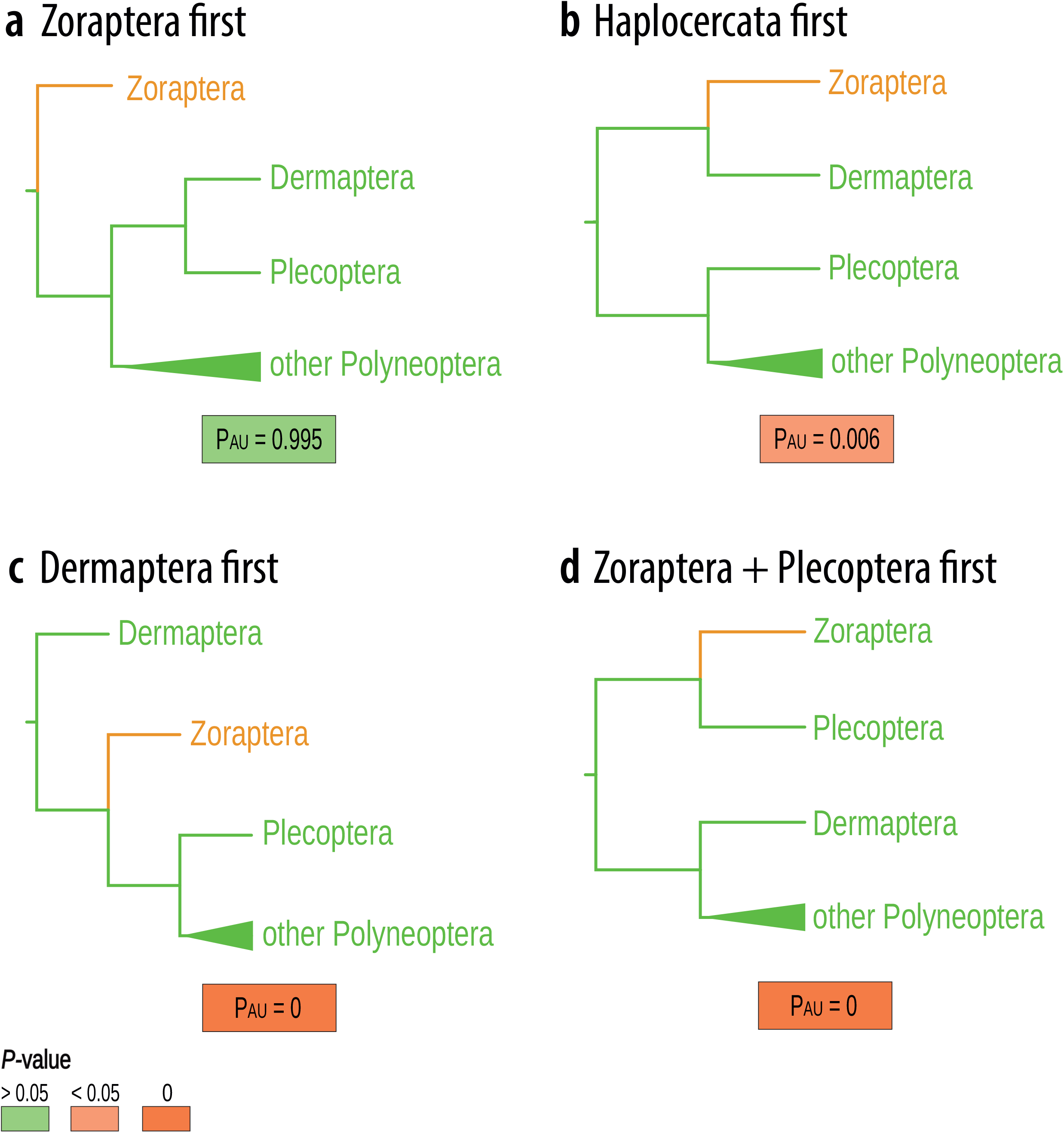
Competing hypotheses of Zorapteran placement, compared with Approximately Unbiased (AU) tests conducted with the LG+C60+F+G model in IQ-TREE. **a** Zoraptera first hypothesis supported by our analyses with the CAT-GTR+G and LG+C60+F+G models and transcriptomic analyses in Simon et al.^40^ and Letsch and Simon ^6^. **b** Haplocercata first hypothesis supported by transcriptomic analyses of Misof et al.^4^, Wipfler et al.^2^. **c** Dermaptera first hypothesis, sister to remaining Polyneoptera recovered by Wipfler et al.^2^ with the multispecies coalescent method. **d** Zoraptera + Plecoptera first hypothesis, supported by Letsch and Simon^6^ analysis using unreduced matrix. *P*-value > 0.05: topology not rejected; *P*-value < 0.05: topology rejected significantly; *P*-value = 0: topology rejected with high significance. The 20 tested topologies are listed in full in Tab. S1.

### Cretaceous fossils shed light on the biology of stem-group zorapterans

Fossils, in particular stem-zorapterans, may provide an independent line of evidence for evaluating competing hypotheses of polyneopteran relationships. However, all known zorapteran fossils from the Mesozoic only narrowly diverge from the body plan of extant zorapterans. To date, all described zorapterans were placed into the genus *Zorotypus*, with the sole exception of *Xenozorotypus burmiticus* from Cretaceous Burmese amber. *Xenozorotypus* possesses plesiomorphic characters such as the presence of the vein (M3+4) in the hind wing, which is otherwise absent from all extant zorapterans^29^. Other Cretaceous stem-zorapterans are included in the subgenus *Octozoros*.

Here we describe a single apterous male zorapteran *Zorotypus nascimbenei*^29^ (full morphological description provided in the Supplementary Information). Along with other members of the subgenus *Octozoros, Z. nascimbenei* is set apart from all crown-group zorapterans by its reduced antennae with eight segments, and plesiomorphic presence of a strong and expanded empodium on the metatarsus and jugate setae^41^. The exquisite preservation of *Z. nascimbenei* reveals novel details of stem-zorapteran palaeobiology. Over a hundred dark ~6 μm biconcave discal fungal spores are preserved around the appendages and attached to the abdomen of the fossil (Fig. 3c, f: fs). Their occurrence in small clusters (Fig. 3c, f, e: fsc) suggests that they were vectored by the zorapteran prior to entombment and not dispersed independently, as similar preservation is also found in pollen grains in amber transported by pollinators^42^. The translucent abdomen reveals similar darkened specks inside (Fig. 2c: fs-abd). The spore’s complete entombment in the amber matrix and resolution limitations of light microscopy preclude us from assigning them to a formal palynotaxon. Fungivory in *Z. nascimbenei* is also supported by the densely setose galeal brush of the fossil (Fig. 3b: gb). Apically setose galeae are also found in crown-group zorapterans and are thought to function in in moving small food particles to the oral cavity^21^. The grinding molae of the mandibles (Fig. 3b: md) may have been used to crush fungal matter prior to ingestion^21^. The association of *Z. nascimbenei* with fungal spores provides the earliest direct evidence of fungivory in Zoraptera and indicates that this feeding strategy had been exploited at least as early as the mid-Cretaceous. Modern zorapterans are opportunistic omnivores, feeding on fungal spores, hyphae, predaceous on springtails, mites and nematodes, occasionally cannibalistic, and scavenging dead arthropods^43^.

**Figure 3.**
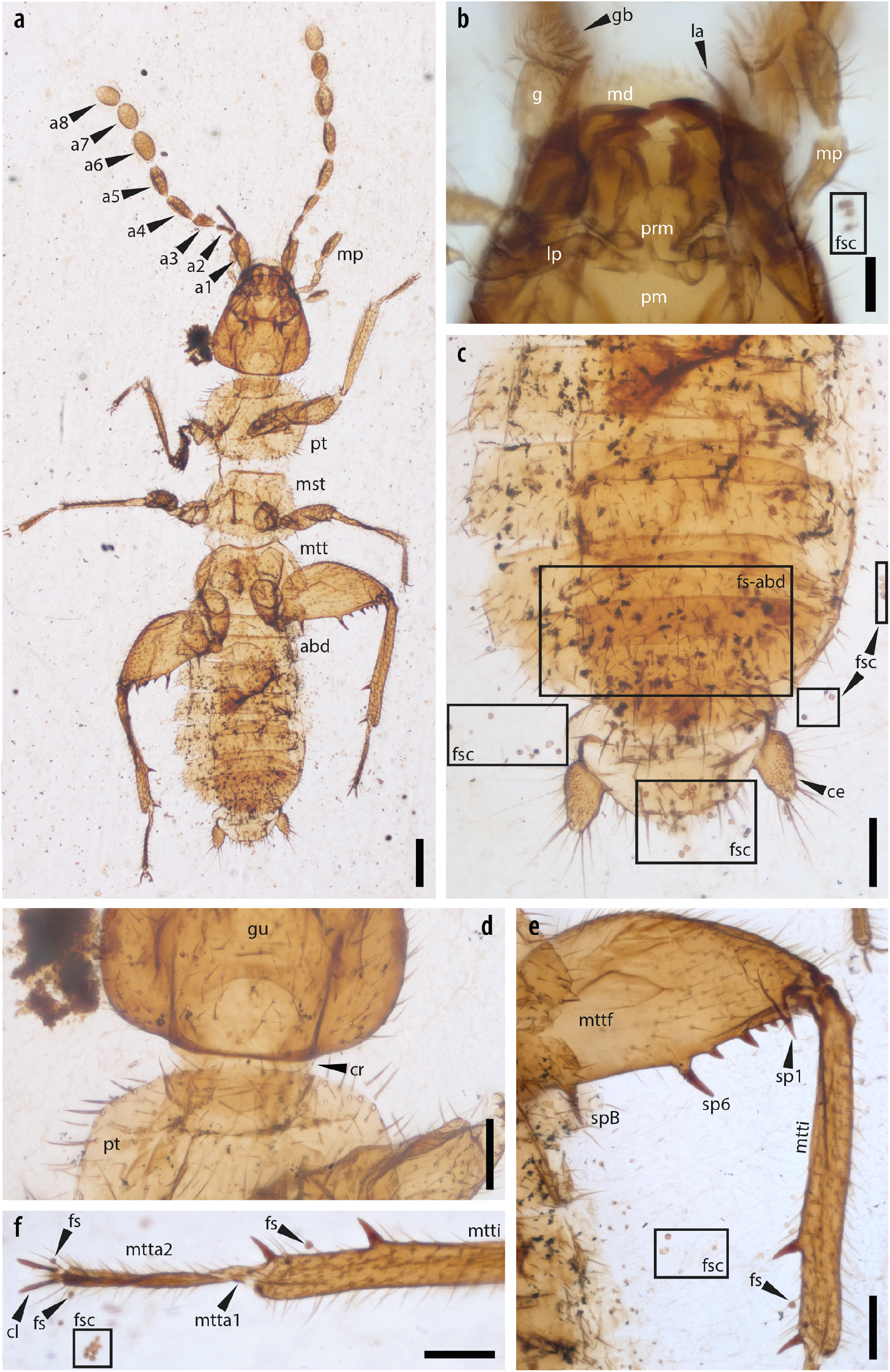
Stem-group zorapteran *Zorotypus nascimbenei* in mid-Cretaceous amber from northern Myanmar (~99 Ma; NIGP175112). **a** Habitus in ventral view. **b** mouthparts in dorsal view. **c** Abdominal apex in dorsal view. **d** Head-prothorax junction in dorsal view. **e** Metafemur and metatibia. **f** Metatibial apex and metatarsi. Abbreviations: a1–8, antennomeres 1–8; abd, abdomen; ce, cercus; cr, cervix; cl, pretarsal claw; fs, fungal spore; fsc, cluster of fungal spores; fsc-abd; putative fungal spores preserved inside the abdomen; g, galea, gb, apical galeal setose brush; gu, gula; la, lacinia; lb, labial palpus; md, mandible; mp, maxillary palpus; mst, mesothorax; mtt; metathorax; mtta1–2, metatarsomeres 1–2; mttf, metafemur; mtti, metatibia; pm, postmentum; prm, prementum; pt, prothorax, sp1–6, metafemoral spines 1–6; spB, basal metafemoral spine. Scale bars: 200 μm (**a**), 100 μm (**c–f**), 50 μm (**b**).

### Timescale of polyneopteran evolution

Previous divergence time analyses estimated an Early Devonian to the Middle Triassic origin of Polyneoptera, depending on the topology and analytical methods used^4,7,11–15^. Some of these estimates are untenable based on their incongruence with the palaeontological record, since polyneopterans have a fossil record extending from the Carboniferous^44^. To reconcile the timescale of polyneopteran evolution with the fossil record and our resolution of the Zoraptera problem, we established 42 new calibration points, more than have been used in any previous molecular clock analysis of winged insects. Furthermore, all calibrations are fully justified with respect to their stratigraphic age and phylogenetic position, in line with best practice recommendations^45^. Contrary to some previous analyses, we specified objective maximum age constraints based on the absence of given lineages in well-explored fossil deposits preserving insects. The full justified list of calibrations is provided in the Supplementary Information. These serve as a basis for future analyses of insect evolution, as well as our own.

We recovered a middle Silurian – Early Devonian (427–399 Ma) origin of the winged insects (Pterygota; Fig. 1). Our molecular clock analyses recovered a late Early to Late Devonian (402–365 Ma) origin of crown-Polyneoptera. These dates correspond broadly with previous molecular clock estimates of insect diversification^4,46,47^. They fall within the notorious ‘hexapod gap’, a period between the Early Devonian and Late Carboniferous which preserves little terrestrial sedimentary rock exposures and where insect fossils are absent^48^.

Under the ‘Zoraptera-first’ scenario, the divergence of the lineage that gave rise to Zoraptera is backdated to the Devonian (Tab. 1). This significantly predates previous estimates that dated the origin of the lineage in the early Permian–Carboniferous ^4,16^. The ancestral lineage of Dermaptera and Plecoptera diverged in a Late Devonian Pennsylvanian interval (368–314 Ma). This is in congruence with the fossil record; stem-Plecoptera and †Protelytroptera (a group of Palaeozoic dermapteran-like insects) co-occur in some Permian localities^13,49,50^, suggesting that their earliest common ancestor must have existed before this date and a putative protelytropteran is known from the late Carboniferous^51^.

**Table 1.**
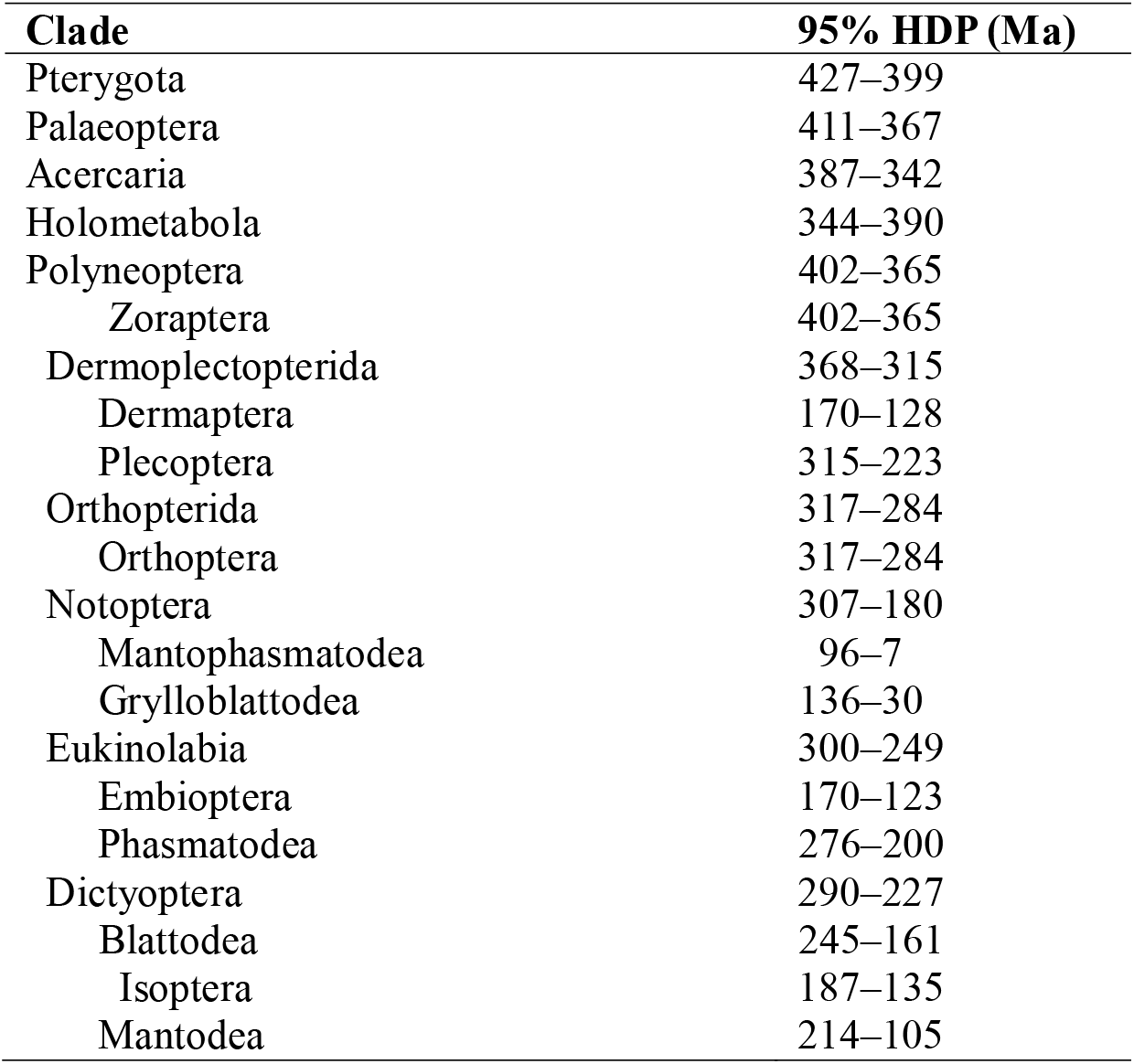
Ninety-five per cent HPD age estimates of divergences between major winged insect (Pterygota) crown groups. Full dated trees are provided in Figs S24 and S25.

The lineages giving rise to the crown-clades Eukinolabia and Dictyoptera originated in the Late Pennsylvanian – Early Triassic (300–249 Ma) and early Permian – Late Triassic (290–227 Ma), respectively. The origin of termites (clade Isoptera of Blattodea) was dated to the Early Jurassic – earliest Cretaceous (187–135 Ma).

### Phylogeny of Zoraptera

With the position of Zoraptera in the insect tree of life now becoming increasingly clear, the last remaining aspect of the ‘Zoraptera problem’ pertains to the internal phylogeny and classification of the order itself. As one of the most morphologically homogeneous insect orders, finding reliable external morphological characters for zorapteran classification has proven notoriously difficult. Zorapterans have been traditionally placed into the single genus *Zorotypus* belonging to the single family Zorotypidae^20^. In the past, some morphological workers have proposed organising extant species into as many as eight separate genera on the basis of wing venation^19^ and morphology of the mouthparts and tarsi^52^, but this alternative classification scheme has not been widely adopted as some of the proposed generic characters are continuous across taxa or highly variable within species^24^. Recent molecular analyses identified multiple clades within the order^16^, leading Kočárek et al.^53^ to divide the order into two families and nine genera. However, the internal relationships within Zoraptera show incongruences between analysed datasets^16,53^, questioning the stability of this new classification. To re-evaluate the contentious intraordinal relationships of Zoraptera, we mined GenBank for publicly available zorapteran sequences. The resultant dataset included 5 molecular markers with a nucleotide alignment of approximately 1,700 bp in length (data occupancy: 65.0%). We also prepared a morphological dataset sampling 10 extinct and extant zorapteran species, with a stonefly as the outgroup.

The molecular analyses run with four models converged on a broadly similar topology of Zoraptera, with the site-heterogeneous CAT-GTR+G and the best-fitting model GTR+F+R3 under a maximum likelihood (ML) framework supporting an identical tree (Fig. 4). In all analyses, Zoraptera formed two main clades with strong to moderate support (BPP = 0.79–1; bootstrap value = 63–100): (i) a clade more or less corresponding to ‘Clade 1’ of Matsumura et al.^16^ (‘Zorotypidae’ of Kočárek et al.^53^) where males possess asymmetrical genitalia, and (ii) a second clade including males with symmetrical genitalia (‘Clade 2’ in^16^, ‘Spiralizoridae’ in^53^). However, relationships among some species and the recently proposed genera were incongruent with previous analyses, rendering Kočárek’s et al.^53^ Zorotypinae, Spermozorinae, and *Spermozoros* paraphyletic. Unlike in Matsumura et al.^16^, we recovered *Z. barberi* as sister to Clade 1 with moderate support, not Clade 2. The position of *Z. novobritannicus* within Spiralizoridae was incongruent among the analyses with different models (Figs S27–30).

**Figure 4.**
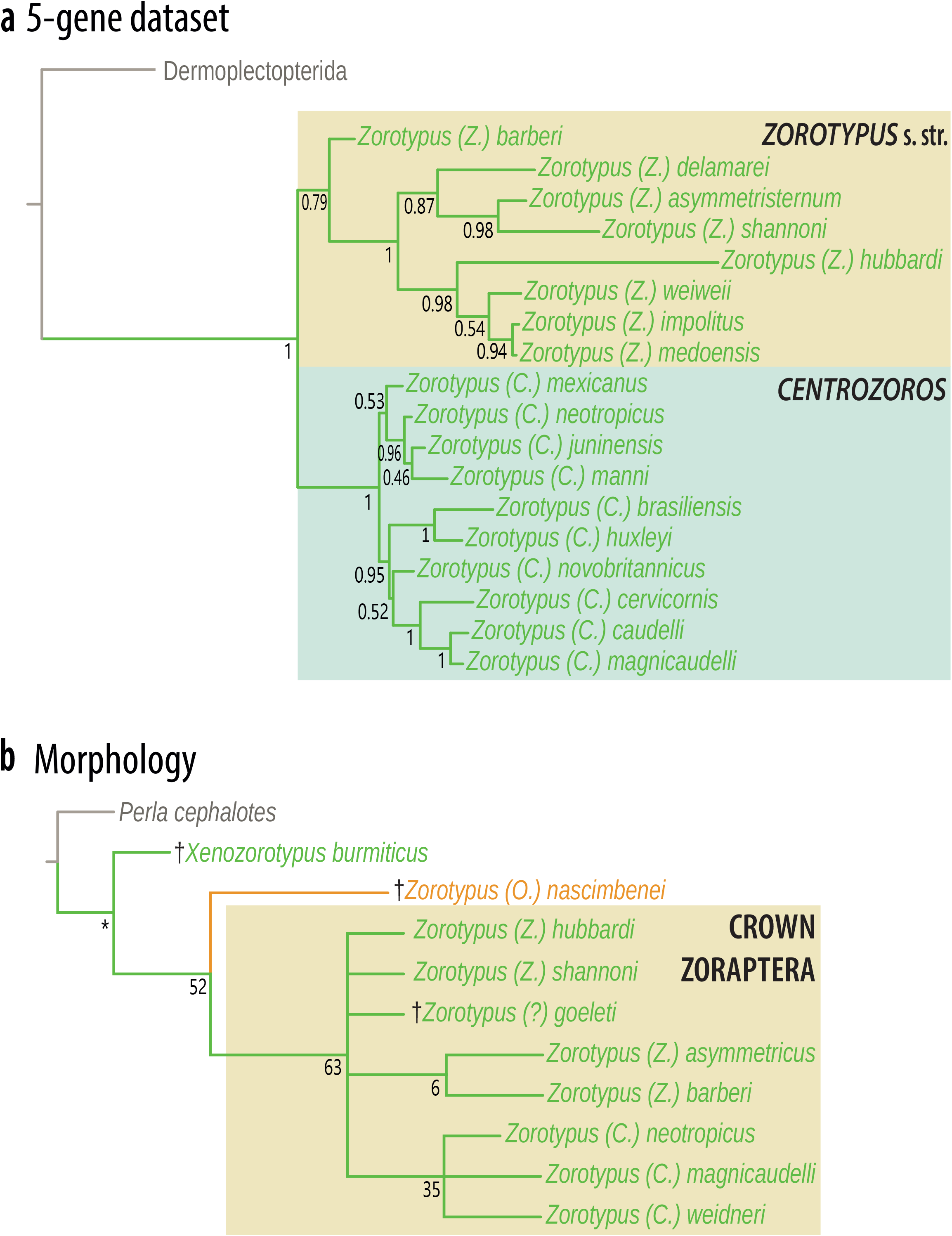
Intraordinal relationships within Zoraptera. **a** Five-gene nucleotide alignment analysed with the CAT-GTR+G model in PhyloBayes. **b** Morphological dataset sampling extinct species, under maximum parsimony in TNT (CI = 0.895; RI = 0.778); character states are mapped in Fig. S31. Support values presented as BPP (**a**) and bootstrap values with 1,000 replicates (**b**).

A morphological phylogenetic analysis recovered *Xenozorotypus* as sister to the remaining zorapterans. The subgenus *Octozoros* is the next diverging clade. Relationships within *Zorotypus* were poorly resolved, with *Z. asymmetricus* + *Z. barberi*, and *Z. neotropicus* + *Z. magnicaudelli* + *Z. weidneri* forming poorly supported clades (bootstrap value = 31–41).

Given that morphological and molecular studies conducted by Matsumura et al.^16^, Kočárek et al.^53^ and in this study support a deep split in the genus *Zorotypus*, into taxa with symmetrical and asymmetrical genitalia, we update zorapteran classification accordingly. We consider extant zorapterans with asymmetrical genitalia as belonging to the subgenus *Zorotypus sensu stricto* (equivalent to ‘Zorotypidae’ of Kočárek et al.^53^), while taxa with symmetrical male genitalia are assigned to the subgenus *Centrozoros* stat. rev. (Tab. 2). This conservative arrangement accommodates uncertainties in *Zorotypus* phylogeny (discussed above) and can be expanded, pending more extensive taxon and gene sampling in future studies. A detailed classification of Zoraptera with a list of species is provided in the Supplementary Information.

**Table 2.**
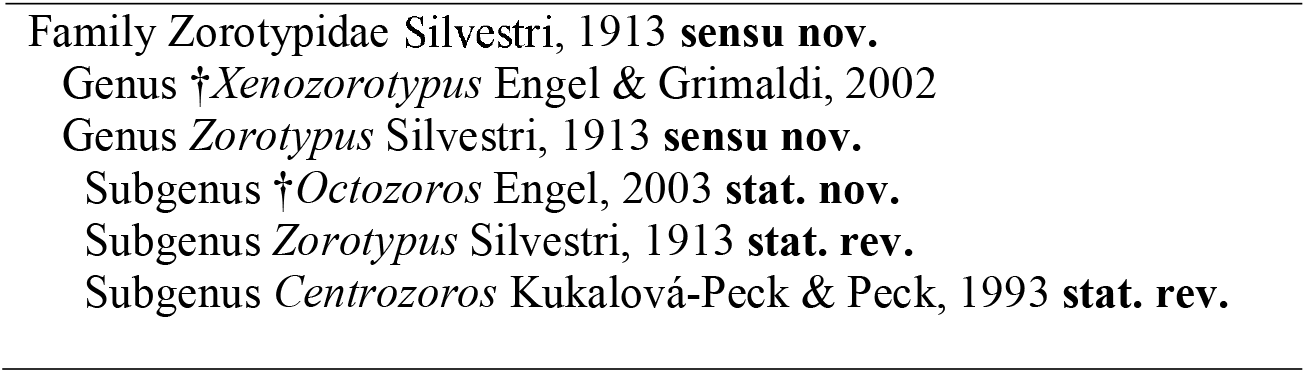
Updated classification of Zoraptera proposed herein, highlighting adopted taxonomic changes. A full checklist of described species and their geographical distribution is provided in the Supplementary Information.

## Discussion

### Phylogenomics, morphology and fossils: congruence in the placement of Zoraptera

Zoraptera represents a typical fast-evolving group, connected by a long branch to the rest of the taxa as shown in many previous analyses^54,55^, suggesting that its placement might be prone to be affected by tree reconstruction artifacts. Zoraptera is recovered as sister to Dermaptera (Haplocercata first) only with less fitting models, both compositionally site-homogeneous models or compositionally site-heterogeneous models with an insufficient number of compositional categories. Analyses with a higher number of substitutional categories (e.g., LG+C60+F+G and CAT-GTR+G) consistently support Zoraptera as the earliest-diverging polyneopteran order, while Dermaptera and Plecoptera form the next branch. As such, our results lend support to the ‘Zoraptera first’ hypothesis, while we regard the Haplocercata hypothesis as a phylogenetic artefact caused by systematic error.

Morphological evidence, from both extant and fossil insects, provides complementary evidence for the placement of Zoraptera. Zorapterans possess numerous plesiomorphies with respect to the remainder of Polyneoptera, such as the morphology of the pro- and mesothoracic spinae, pleural arm, and membranous anapleural suture^56^. A close relationship between Zoraptera and the early-diverging Dermaptera and Plecoptera has been suggested by several morphological studies. Silvestri^17^, who described Zoraptera in 1913, discussed superficial similarities between the order and Dermaptera. Both notably share gregarious habits^57^ and maternal care in some species^58^. Zoraptera, Plecoptera, and Dermaptera share the absence of the muscle *Musculus hypopharyngomandibularis*, a plesiomorphy shared with Palaeoptera^59^. Zorapterans and dermapterans are the only polyneopteran groups with holocentric chromosomes, an apparently plesiomorphic condition otherwise found in Palaeoptera^2,60^. *Zorotypus hubbardi* shares the number of male diploid chromosomes (2*n* = 38) with the dermapteran *Prolabia* (within Polyneoptera, this condition is otherwise present only in some Isoptera and Phasmatodea)^61,62^. Kuznetsova *et al*.^62^ hypothesised an ancestral karyotype of 2*n* = 40 for Zoraptera, which is shared with the early-diverging dermapteran family Labiduridae, and within Polyneoptera is otherwise found only in some Isoptera^61^. Friedrich *et al*.^56^ found similarities in the thoracic skeletal structures of Zoraptera and Plecoptera, such as in the morphology of the prothoracic basisternum which is partly reduced and spoon-shaped in both orders. Both orders likewise possess reduced ovipositors^63^. In the head, Zoraptera and Plecoptera share tormae without mesal extensions and 5 incisivi on the left and right mandibles^59^. Zorapteran winged morphs have a kidney-shaped frontal spot also present in some plecopterans^64^, although the homology of this structure is unclear^21^.

Kukalová-Peck and Peck^19^ observed that the head shape in Zoraptera with antennal articulations close to the ventral cranial margin, represents a plesiomorphy within Insecta, similar to fossil pterygotes from the Palaeozoic. According to the interpretation of Haas and Kukalová-Peck^65^, Zoraptera, Protelytroptera, Dermaptera, and Plecoptera all share a mp-cua cross-vein (arcus). Protelytroptera, Dermaptera, and Plecoptera share a large anal fan on the hind wing in their ground plan.

The Dermaptera + Plecoptera clade is supported by wing base structure, namely the morphology of the ventral basisubcostale, and the articulation between the antemedian notal wing process and first axillary sclerite^66^. Both groups share the absence of male gonostyli and a functional ovipositor (both shared with Zoraptera), and paired male gonopore^66,67^. An early-diverging Dermaptera + Plecoptera clade (dubbed Dermoplectopterida) was recovered in a *18S* analysis^68^ and some early phylotranscriptomic studies^6,40^, and is now supported by our own analyses accounting for common sources of systematic error. We herein formally propose the name Eupolyneoptera Engel, Tihelka, & Cai clade nov. nom. for all Polyneoptera excluding Zoraptera (see the Supplementary Information for a full systematic treatment and synapomorphies).

### The nature of the ancestral polyneopteran

Hennig^20^ regarded the last ancestor of “Paurometabola” (i.e., his concept of Polyneoptera, but excluding Zoraptera and Plecoptera) as probably a ground-dwelling insect with forewings modified into leathery tegmina that lived during the Carboniferous. Recently, molecular studies have attempted to reconstruct morphology, life history, and chromosomal system of the last common ancestor of Polyneoptera^2,61,69^. Reconstructing the root of Polyneoptera is heavily dependent on the resolving the inferred relationships of the early-diverging orders.

Our analyses suggest that the earliest common ancestor of crown-polyneopterans lived in the Devonian. This coincides with the diversification of vascular plants and animals on land^70^. The Devonian saw an abrupt spread of vegetation and increase in plant size^71–73^, which would have provided structured habitats rich in plant biomass such as leaf litter, dead wood, and spaces under bark that are occupied by early-diverging polyneopterans today^2^. These diversified spaces would have made the newly folded wings of these early neopterans beneficial for insects invading plant crevices and other spaces, allowing for access to microhabitats previously inaccessible by palaeopterous insects. It also freed wings, when not in use, to evolve secondary specializations, such as protective covers, etc. (e.g., Dermaptera). Since the three basalmost polyneopteran orders, Zoraptera, Dermaptera, and Plecoptera, all represent opportunistic predators, scavengers, phytophages, and detritivores^43,74^, this suggests such a biology was plesiomorphic to the clade and that the increase in trophic complexity during the Devonian may have opened new niches for the group that gave rise to Polyneoptera. All groups that constitute the diet of extant zorapterans, namely terrestrial fungi, springtails, possible insects, mites, and nematodes make their first appearance in the fossil record in the Devonian^75,76^. Diverse terrestrial fungi first occur in the fossil record in the Devonian^77^; notably the specimen of the stem-zorapteran *Z. nascimbenei* herein possesses specializations for feeding on fungal spores. Since our understanding of insect diversity in the Devonian is severely hampered by the rarity of well-preserved terrestrial sediments of this age^48^, further analyses of the fossil record will be required to test if the basal radiation of insects coincided with the diversification of land plants and animals in the Devonian. However, because unequivocal fossil stem-polyneopterans remain to be identified, the age of total-group Polyneoptera may predate the Devonian.

The ancestral polyneopteran is unlikely to have resembled extant Zoraptera or stem-zorapterans, as numerous characteristics of the order such as the greatly reduced wing venation evidently represent specializations for highly derived, subcortical life in late-stage decaying wood and consequences of miniaturisation, as zorapterans don’t exceed 3 mm in length^6,20,21^. With Zoraptera as sister to the remaining Polyneoptera, it is possible that the ancestral polyneopteran could have exhibited gregarious behaviour and biparental care for offspring^78^, although these are more likely derived specializations of Zoraptera, just as maternal care is indicative of crown-Dermaptera but was absent in stem-Dermaptera^79^. Resolution of the position of stoneflies (Plecoptera), which possess aquatic nymphs, as sister to Dermaptera within Dermoplectopterida also suggests that the ancestral Polyneopteran was likely terrestrial, contrary to the widespread view that the earliest flying insects originated in aquatic habitats^80,81^.

## Conclusion

The relationships among the early-diverging orders of Polyneoptera, one of the three major clades of insects with folding wings (Neoptera), represent one of the most persistent problems in insect evolution. We reanalysed the most complete protein-coding gene dataset available for the group to interrogate common sources of tree reconstruction errors to infer the early evolution of Polyneoptera. We recover a novel topology, with the enigmatic Zoraptera as sister to the remaining polyneopteran orders, and Dermaptera + Plecoptera (Dermoplectopterida), as the next branch, in congruence with some previous molecular and morphological studies. By experimenting with different models of molecular evolution, we show that previous topologies within Polyneoptera are only supported by analyses conducted within a narrow analytical window using poorly fitting substitution models. A newly discovered fossil of the Cretaceous stem zorapteran *Zorotypus nascimbenei* preserves fungal spores in its abdomen, providing insights into the dietary diversity of Mesozoic zorapterans. Together with its specialised mouthparts and the presence of a spore brush on the apex of galea, *Octozoros* provide the earliest direct evidence of feeding on fungal spores in Polyneoptera and further illuminates the ancestral biology of the group. Our results highlight the importance of adequate modelling of molecular evolution and seeking a consensus of molecular and morphological evidence in resolving contentious problems in insect phylogeny.

## Methods

### Dataset assembly and phylogenetic reconstruction

For the phylogenomic analyses, we used the multiple sequence alignment generated by Wipfler et al.^2^, sampling all polyneopteran orders represented by 106 species. With 3,014 protein-coding single-copy genes, it represents the most comprehensive dataset for Polyneoptera compiled to date. To mitigate errors due to poor alignment and incorrect identification of orthologs^30^, the decisive amino acid (AA) dataset was trimmed with BMGE using BLOSUM95 with -h set to 0.4^82^. Trimming reduced the original dataset by 84.7% (from 1,246,506 AA sites to 191,242) and increased data occupancy from 52.2% to 92.0%. The exclusion of highly variable/uncertain regions reduced the average pairwise distance among taxa by 50% (from 0.30 to 0.15). We used the trimmed alignment to generate four matrices with variable taxon sampling. We excluded two and six taxa that behaved as rouge taxa in preliminary analyses, giving datasets with 106, 102, and 100 taxa. The excluded taxa were *Liposcelis* (Psocoptera) and *Pediculus* (Phthiraptera) in the 104-taxon dataset, in addition to *Apachyus* (Dermaptera), *Brachyptera* (Plecoptera), *Leuctra*, and *Perla* (Plecoptera) in the 100-taxon dataset. Wipfler *et al*.^2^ pointed out that the position of Zoraptera may be unstable because of unbalanced taxon sampling, since the order is often represented by only a single species in phylogenetic analyses. To investigate this possibility, we produced a fourth matrix which included only one representative of each order (33 taxa dataset).

The status of Zoraptera as a fast-evolving group prone to systematic bias has been long known^54,55^. To overcome this problem, Wipfler *et al*.^2^ used ModelFinder in IQ-TREE to partition their decisive alignment and analyse individual partitions with best-fitting default site-homogeneous models and the LG4X mixture model. In a recent simulation study, Wang et al.^83^ found that, on a simulated and empirical datasets, partitioning data and analysing partitions with site-homogeneous models is still prone to systematic errors and underperforms in comparison with analysing concatenated datasets with site-heterogeneous models. Hence, we tested the effect of using heterogeneous models of varying computational complexity on polyneopteran phylogeny. We tested the performance of the site-homogeneous models LG+G and GTR+G, variations of which have been used in past polyneopteran phylogenomic studies^2,4^, the LG4X mixture model, and the compositionally site-heterogeneous LG+C60+F+G and CAT-GTR+G, the latter of which has been shown theoretically and empirically to better describe substitution in metazoan genomes^33–35,84^. To compare the fit of the these models to our polyneopteran dataset, we run a 10-fold Bayesian cross-validation analysis in PhyloBayes MPI 1.7^36^ with 10 replicates.

Analyses with the site-homogeneous models LG+G and GTR+G use a single amino acid substitution matrix to describe the biochemical properties of amino acids. Variations of these models have been widely used in previous analyses of polyneopteran phylogeny. In our analyses, they were conducted in IQ-TREE 1.6.3^85^. Analyses employing the LG4X, that accounts for substitution rate heterogeneity across the alignment by estimating four different LG+G substitution matrices^86^, were likewise performed in IQ-TREE. The compositionally site-heterogeneous profile mixture model LG+C60+F+G implements 60 equilibrium frequency categories^87^ and was run in IQ-TREE. Support values for all analyses were generated using 1,000 ultra-fast bootstraps.

Analyses with the site-heterogeneous infinite mixture model CAT-GTR+G were performed in in PhyloBayes by running two independent Markov chain Monte Carlo (MCMC). The bpcomp program was used to evaluate convergence by generating output of the largest (maxdiff) and mean (meandiff) discrepancies observed across all bipartitions. Due to computational constraints, only the 100- and 33-taxon datasets were run to complete convergence (maxdiff_100_ = 0; maxdiff_25_ = 0.097). The remaining analyses did not converge (maxdiff_102,106_ = 1), however the recovered topology was identical to the converged datasets only with a few nodes poorly supported.

### Topology tests

To evaluate support for 20 alternative hypotheses of polyneopteran evolution, we conducted topology tests using the 100-taxon dataset in IQ-TREE using the site-heterogeneous LG+C60 model. This model was used because the best fitting CAT-GTR+G is not implemented in IQ-Tree, but LG+C60+F+G (similarly to CAT-GTR+G) can still accommodate (despite it is not best fit), compositional heterogeneity and would fit the data batter than the other models available in IQ-Tree. To perform the topology tests the Approximately Unbiased (AU) test was used, with 10,000 resamplings performed using the RELL method. We focused on testing historical and recent hypotheses of Zoraptera as well as Dermaptera and Plecoptera phylogeny, since the relationships of these three orders have produced the most incongruence in phylogenetic studies of Polyneoptera^3^.

### Molecular clock analyses

To establish a timescale of polyneopteran evolution congruent we followed the best-practice recommendations for the selection of calibrations outlined by Parham et al.^45^. Each fossil-based node calibration is based on a museum-curated fossil specimen or group of specimens, phylogenetic and age justifications for each are provided in the Supplementary Information. In total, 42 calibrations spread equitably throughout the tree were used, the calibrated nodes are displayed in Fig. S1.

For example, the roached *Qilianiblatta namurensis* from the Tupo Formation in northwestern China^44,88^ was used to calibrate the node representing stem-Dictyoptera. The preserved right forewing shares with extant Blattodea a deeply concave CuP vein^89^, but the RA with branches translocated to RP indicates that the fossil belongs to stem Dictyoptera^44^. Roachoids’ such as *Q. namurensis* were roach-like insects abundant in the Paleozoic and some of the earliest winged insects in the fossil record. While some authors consider Paleozoic roachoids as close to extant Blattodea^90^, they are excluded from crown Blattodea and Mantodea by wing venation character and most notably by the presence of long external ovipositors in females^91^. Previous analyses have shown that excluding Carboniferous roachids from molecular clock analyses leads to underestimates of polyneopteran timescale of diversification^11^. The fossiliferous horizon yielding the specimen has been dated to the Namurian B/C or to the Bashkirian (latest Duckmantian), based on the presence of a characteristic ammonoid and conodont fauna^88,92^. Trümper et al.^92^ proposed an upper bound on the age of the insect-bearing bed of ~315 Ma, which we used as the minimum constraint on the node. The maximum age, 428.9 Ma, is taken from the two oldest Lagerstätten preserving terrestrial animals, the Přídolian Ludford Lane in Shropshire^93^, England and the Pragian Rhynie chert from Aberdeenshire, Scotland

The geological timescale follows Ogg et al.^94^. Molecular clock analyses were run in MCMCTree implemented in PAML 4.7^95,96^. We obtained 200,000 trees with a sampling frequency of 50 and discarded 10,000 as burn-in. Default parameters were set as follows: ‘cleandata=0’, ‘BDparas=1 1 0’, ‘kappa_gamma=6 2’, alpha_gamma=1 1’, ‘rgene_gamma= 2 20’, ‘sigma2_gamma=1 10’, and ‘finetune=1: 0.1 0.1 0.1 0.01 .5’. Convergence was assessed by plotting posterior mean times from the first run against the second run. Analyses were run using a uniform prior distribution between minimum and maximum age constraints. We used soft bounds with maximum and minimum tail probabilities of 2.5%. Analyses both with the autocorrelated rates clock (AC) model and independent rates (IR) clock model, and our results integrate the results from both. To ensure our priors were appropriate, we ran the MCMCTree analysis without sequence data to calculate the effective priors, compared to the specified priors.

### Palaeontology

The studied amber inclusion originates from mines near the summit of the Noije Bum hill in the Hukawng Valley, Kachin State, northern Myanmar. Radiometric dating of the amber deposit has provided an early Cenomanian age, ~99 Ma, while palaeontological evidence suggests that the amber is no older than late Albian^97,98^. The specimen has been cut with a razor blade and polished using diatomite powder. Photographs under normal visible light were taken with a Canon EOS 5D Mark III digital camera, equipped with a Canon MP-E 65 mm macro lens (F2.8, 1–5×), and with an attached Canon MT-24EX twin flash. Epifluorescence photomicrographs were taken using a Zeiss Axio Imager 2 microscope under the eGFP mode (Zeiss Filter Set 10; excitation/emission: 450–490/515–565 nm). Extended depth of field images were digitally compiled in Zerene Stacker 1.04. The type specimen is deposited in the amber collection of the Nanjing Institute of Geology and Palaeontology (NIGP), Chinese Academy of Sciences under the accession number NIGP175112. The amber piece was purchased from a local amber miner in late 2016, complying with the laws of Myanmar and China, and is open to legitimate study^99^.

### Molecular and morphological phylogeny of Zoraptera

To elucidate the intraordinal relationships within Zoraptera, morphological data from extant and fossil species as well as five gene markers were integrated in a total evidence phylogenetic reconstruction. The molecular markers used were the publicly available sequences of the nuclear *18S rRNA*, and *H3*, and the mitochondrial *12S rRNA, 16S rRNA*, and *COI* mined from GenBank. In total, 18 zorapteran taxa identified to species level alongside five plecopteran and dermapteran outgroups were included. Accessions are provided in Supplementary Table 2. The sequences of *COI* and *H3* were unambiguously aligned owing to few gaps and their codon-based structure using the MUSCLE algorithm implemented in the MEGA X^100^. The ribosomal RNAs *12S, 16S*, and *18S*, were aligned in MAFFT using the E-INS-I algorithm^101^. The third codon of *COI* was excluded to reduce data heterogeneity. We analysed our data with the Bayesian site-heterogeneous CAT-GTR+G model implemented in PhyloBayes. We also tested the fit of a range of the default maximum likelihood site-homogeneous models implemented in IQ-TREE. The results are displayed in Tab S3; GTR+F+R3 was the best-fitting model by the corrected Akaike Information Criterion (AICc = 20871.1) and TVM+F+I+G by the Bayesian Information Criterion (BIC = 21152.0), while JC had the poorest fit to the alignment overall (AICc = 23033.1; BIC = 23264.6). We thus used these three ML models to run further analyses of the nucleotide alignment.

For a morphological phylogenetic analysis, we scored 12 morphological characters for 10 ingroup taxa. The principal objective was to address the relationships of the extinct genus *Xenozorotypus*, the extinct subgenus *Octozoros*, and crown-group zorapterans (*Zorotypus s. s*.). Within the crown-group of *Zorotypus*, we sampled representatives of all key clades recovered by molecular analyses^16^. We included the unusual Miocene species *Z. goeleti* which possesses dimeric cerci and may thus represent the sister group to the remainder of *Zorotypus s. s*.^24^. A single plecopteran representative, *Perla cephalotes*, was used as the outgroup. The character matrix is available in Tabs S4 and S5. Maximum parsimony analyses were conducted in TNT v. 1.5^102^ using implied weighting. The recommended concavity value (*K*) of 12 was used, as these have been shown to achieve higher accuracy against homoplastic characters^103^. Collapsing rules were set to ‘none’ and the analysis was run using default settings in ‘New Technology Search’. A majority-rule consensus tree of the resultant four most parsimonious trees was computed. To assess tree support, nonparametric bootstrap analysis was run with 1,000 replicates. Character states were mapped using ASADO v. 1.61^104^.

## Data availability

Analysed files and results have been uploaded to MendeleyData repository (doi: 10.17632/pk747fvxxp.1).

## Authors’ contributions

C.C., E.T. and P.C.J.D. conceived and designed the study, C.C. and E.T. conducted the phylogenomic analyses, E.T. conducted topology tests, the molecular clock analyses, and analyses of zorapteran intraordinal phylogeny, Z.Y. and C.C. processed the photomicrograph data and studied *Z. nascimbenei* morphology. E.T., Z.Y. and C.C. drafted the manuscript, to which J.L.-F., M.S.E., D.H., M.G., O. R.-S., P.C.J.D. and D.P. contributed.

