## Supplemental Information for "Compositional phylogenomic modelling resolves the ‘Zoraptera problem’: Zoraptera are sister to all other polyneopteran insects"

#### Systematic Entomology

Order Zoraptera Silvestri, 1913
Family Zorotypidae Silvestri, 1913
Genus *Zorotypus* Silvestri, 1913
Subgenus *Octozoros* Engel, 2003

*Zorotypus (Octozoros) nascimbenei* Engel & Grimaldi 2002

Figs 3, S2–3

**Material examined.** 1 apterous male, NIGP175112; lowermost Cenomanian, Hukawng Valley, northern Myanmar; deposited in the Nanjing Institute of Geology and Palaeontology, Chinese Academy of Sciences, Nanjing, China.

**Diagnosis.** Body length 1.48–2.51 mm. Antennae with eight antennomeres, antennomeres 2–4 successively longer, 4–6 approximately of similar length, 6–8 successively shorter. Pronotum not constricted, anterior and posterior margins of similar width. Mesonotum lacking a pair of spines on anterolateral corners. Posterior margin of forewing with seven regularly spaced jugate setae along ventral margin. Metafemur with large basal (*spB*) and middle spines (*sp6*), with five smaller apical spines (*sp1–5*), *sp1* and *sp4* longer than *sp2, 3, 5*; metatibia with two spines at apical third and at apex. Tergite X with thin and short median projection.

**Description (apterous male).** Total body length (exclusive of antennae; including cerci) 2.51 mm; antennal length 1.14 mm; length/width of head 0.49/0.41 mm; pronotum 0.34/0.44 mm; mesonotum 0.28/0.41 mm; metanotum 0.22/0.48 mm; metafemoral length 0.45 mm; metatibial length 0.56 mm; abdominal length 1.25 mm; cercus length 0.09 mm.

Integument light reddish-brown and smooth. Head roundly triangular, as broad as pronotum. Compound eyes and ocelli absent. Antenna relatively long, with eight antennomeres; antennomere 1 (scapus) elongate, approximately 2.8× longer than wide, about as long as combined lengths of next two antennomeres; antennomere 2 (pedicel) slightly curved outward, approximately as long as antennomere 3; antennomeres 4–6 each distinctly longer than 3 and of similar length; antennomeres 6–8 successively shorter, each elongate-oval. Postocular margins straight, temples rounded, constricted posteriorly. Mandibles each with large apical and 2–3 smaller sub-apical teeth. Maxillary palpomere 1 short, palpomeres 2, 3, and 5 distinctly elongate, palpomere 4 slightly longer than wide; labial palpus with elongate palpomeres 1 and 3, and short palpomere 2. Pronotum transverse, and slightly broader than head; sides rounded, posterior and anterior widths equivalent (i.e., not constricted); anterior margin slightly curved; setae scattered, of approximately uniform length; mesonotum broader than long, lacking thorn-like spines on anterolateral corners; metanotum broader than long, shorter than mesonotum, anterior margin markedly emarginate; thorax with scattered, short setae. Metafemur expanded, gradually tapering toward apex; seven stout and strongly sclerotized spines present along posterior border of metafemoral ventral surface, slightly angled toward metafemoral apex; basal spine (*spB*) present, about as large as middle spine (*sp6*), both situated on tubercles*, sp1* and *sp4* distinctly longer than *sp2*, *3* and *5*; metatibia distinctly longer than metafemur, slender, slightly dilated toward apex, with two spines at apical third and at apex. Abdominal terga with scattered, minute setae, lacking distinct, transverse rows of setae along posterior margin; terga lacking stiff, erect setae at posterolateral corners; tergite X incised along posterior margin, roundly projected at middle, with thin and short median projection; tergite XI unmodified, with scattered long setae. Cerci elongate, narrowing toward apex; unsegmented; with scattered setae longer than cercus, lacking apical spine-like seta.

Eupolyneoptera Engel, Tihelka, & Cai clade nov.

**Systematic scope.** All Polyneoptera excluding Zoraptera.

**Diagnosis.** Two lateral cervical sclerites (different in Zoraptera and variously simplified in Plecoptera); tarsal plantulae (sometimes termed bladders in Embiodea) present (absent in Zoraptera, reduced or lost various times in Eupolyneoptera); tarsi plesiomorphically pentamerous but reduced several times independently across clade (pentamerous tarsi in stem-group Dermaptera, Plecoptera, and Phasmatodea); hind wings typically larger than forewings, with enlarged vannus (anal fan) folding along jugal and vannal folds (reduced in Embiodea and Euisoptera, and lost entirely in extant Notoptera, but present in stem-group Notoptera), vannus typically pleated (except when anal region secondarily reduced, *i.e.*, Embiodea, Euisoptera); malpighian tubules numerous, far greater than six (Zoraptera with six); abdominal ganglia unconcentrated (Zoraptera concentrated into single mass); male genitalia with cluster of accessory glands, mesomeres do not fuse to form aedeagus, intromittent organ by eversion of ductus ejaculatorius; ovipositor either vestigial apomorphically (stem-group Plecoptera, Dermaptera, Embiodea, and Dictyoptera have well-developed ovipositors) or third valvulae forming part of ovipositor penetrating apparatus (rather than as sheath).

**Etymology.** A combination of Greek *eu*– (*εὐ*–, meaning, "true" or "good") and Polyneoptera [itself formed of *polús* (*πολύς*, meaning, "many"), *néos* (*νέος*, meaning, "new"), and *pterá* (*πτερά*, plural of *πτερόν*, meaning, "wings")].

Eteopolyneoptera Engel, Tihelka, & Cai clade nov.

**Systematic scope.** Orthoptera, Mecynoptera (Notoptera + Eukinolabia), and Dictyoptera.

**Diagnosis.** Trochantin detached from episternum, separated by membrane (attached, wholly or at least partially, with episternal sulcus in Plecoptera and Dermaptera); hind wing vannus typically pleated (except when anal region secondarily reduced, *i.e.*, Embiodea); male styli present (suppressed in Plecoptera and Dermaptera, and secondarily and independently in Acrididae and Phasmatodea + Embiodea).

**Etymology.** The clade name is a combination of Greek *ἐτεός* (*eteós*, meaning, "true" or "genuine") and Polyneoptera (for derivation, *vide supra*).

#### Classification of Zoraptera

##### FAMILY ZOROTYPIDAE Silvestri, 1913 **sensu nov.**

Zorotypidae Silvestri, 1913: 195. Type genus: *Zorotypus* Silvestri, 1913.
Spermozorinae Kočárek, Horká, & Kundrata, 2020: 9: Type genus *Spiralozoros* Kočárek, Horká, & Kundrata, 2020. **syn. nov.**
Spiralozoridae Kočárek, Horká, & Kundrata, 2020: 10: Type genus *Spiralozoros* Kočárek, Horká, & Kundrata, 2020. **syn. nov.**
Latinozorinae Kočárek, Horká, & Kundrata, 2020: 12: Type genus *Spiralozoros* Kuklová-Peck & Peck, 1993. **syn. nov.**

The recent proposal[1] to split this minute family of largely externally homogenous insects into two families and four subfamilies serves no good purpose, particularly as the family Spiralozoridae as proposed is found to be paraphyletic herein, as is their concept of Zorotypinae. The genera in their system are similarly oversplit, differentiated by the most minute and unremarkable of characters such as body size, colour, pronotal shape and morphology of the metafemora. We conservatively consider the order Zoraptera to consist of a single family, Zorotypidae, and two genera, *Xenozorotypus* and *Zorotypus*. We treat *Zorotypus* as composed of three subgenera: the basal *Octozoros* known exclusively from Cretaceous ambers, alongside the extant *Centrozoros* and *Zoroytpus sensu stricto*. *Centrozoros* males have symmetrical genitalia while they are asymmetrical in *Zoroytpus s. str.* This conservative arrangement takes into consideration the known uncertainties in *Zorotypus* phylogeny and is open to future revision, pending more extensive taxon and gene sampling.

###### GENUS †*XENOZOROTYPUS* Engel & Grimaldi, 2002

*Xenozorotypus* Engel & Grimaldi, 2002: xxx. Type species: *Xenozorotypus burmiticus* Engel & Grimald, 2002, by original designation.

*X. burmiticus* Engel & Grimaldi, 2002.
[Myanmar: Cretaceous amber from the Hukawng Valley, Kachin]

###### GENUS *ZOROTYPUS* Silvestri, 1913 **sensu. nov.**

*Zorotypus* Silvestri, 1913: 196. Type species: *Zorotypus nascimbenei* Engel & Grimaldi, 2002, by original designation.

###### SUBGENUS †*OCTOZOROS* Engel, 2003 **stat. nov.**

*Zorotypus* (*Octozoros*) Engel, 2003: 148. Type species: *Zorotypus guineensis* Silvestri, 1913, by original designation.

*Z. (O.) acanthothorax* Engel & Grimaldi, 2002. (= syn. *Z. (O). hukawngi* **syn. nov.**)
[Myanmar: Cretaceous amber from the Hukawng Valley, Kachin]

*Z. (O.) cenomanianus* Yin, Cai, & Huang 2017. (= syn. *Z. (O). robustus* Liu, Zhang, Cai, & Li, 2018)
[Myanmar: Cretaceous amber from the Hukawng Valley, Kachin]

*Z. (O.) hirsutus* Mashimo, 2018.
[Myanmar: Cretaceous amber from the Hukawng Valley, Kachin]

*Z. (O.) hudae* Kaddumi, 2005.
[Jordan: Cretaceous amber from the Zarqa River Basin]

*Z. (O.) pecten* Mashimo, Müller, & Beutel, 2019
[Myanmar: Cretaceous amber from the Hukawng Valley, Kachin]

*Z. (O.) pusillus* Chen & Su, 2019
[Myanmar: Cretaceous amber from the Hukawng Valley, Kachin]

*Z. (O.) nascimbenei* Engel & Grimaldi, 2002

[Myanmar: Cretaceous amber from the Hukawng Valley, Kachin]

##### SUBGENUS *ZOROTYPUS* Silvestri, 1913 **stat. rev.**

*Zorotypus* Silvestri, 1913: 196. Type species: *Zorotypus nascimbenei* Engel & Grimaldi, 2002, by original designation.

*Z. (Z.) amazonensis* Rafael & Engel, 2006.
[Brazil]

*Z. (Z.) asymmetricus* Kočárek, 2017 **comb. restit.**
[Brunei]

*Z. (Z.) asymmetristernum* Mashimo, 2019.
[Kenya]

*Z. (Z.) caxiuana* Rafael, Godoi, & Engel, 2008.
[Brazil]

*Z. (Z.) delamarei* Paulian, 1949.
[Madagascar]

*Z. (Z.) guineensis* Silvestri, 1913.
[Guinea, Ghana, Ivory Coast]

*Z. (Z.) huangi* Yin & Li, 2017 **comb. restit.**
[China: Yunnan]

*Z. (Z.) hubbardi* Caudell, 1918, **comb. restit.**
[USA]

*Z. (Z.) impolitus* Mashimo, Engel, Dallai, Beutel & Machida, 2013) **comb. restit.**
[Peninsular Malaysia]

*Z. (Z.) medoensis* Huang, 1976 **comb. restit.**
[China: Tibet]

*Z. (Z.) shannoni* Gurney, 1938.
[Brazil]

*Z. (Z.) sinensis* Huang, 1974 **comb. restit.**
[China: Tibet]

*Z. (Z.) vinsoni* Paulian, 1951.
[Mauritius]

*Z. (Z.) weiweii* Wang & Cai, 2016 **comb. restit.**
[Borneo]

##### SUBGENUS *CENTROZOROS* Kukalová-Peck & Peck, 1993 **stat. rev.**

*Centrozoros* (Kukalová-Peck & Peck, 1993): 342. Type species: *Zorotypus gurneyi* Choe, 1989, by original designation.

*Z. (C.) barberi* Gurney, 1938 **comb. restit.**
[Panama, Costa]

*Z. (C.) brasiliensis* Silvestri, 1946 **comb. restit.**
[Brazil]

*Z. (C.) buxtoni* Karny, 1932, **comb. restit.**

[Samoa]

*Z. (C.) caudelli* Karny, 1922, **comb. restit.**
[Peninsular Malaysia, Sumatra]

*Z. (C.) cervicornis* Mashimo, Yoshizawa and Engel, 2013 **comb. restit.**
[Peninsular Malaysia, Borneo]

*Z. (C.) ceylonicus* Silvestri, 1913, **comb. restit.**
[Sri Lanka]

*Z. (C.) cramptoni* Gurney, 1938) **comb. restit.**
[Guatemala]

*Z. (C.) gurneyi* (Choe, 1989) **comb. restit.**
[Costa Rica, Panama]

*Z. (C.) hainanensis* Yin, andWu, 2015 **comb. restit.**
[China: Hainan]

*Z. (C.) hamiltoni* (New, 1978)
[Barbados, Colombia]

*Z. (C.) huxleyi* Bolívar Pieltain and Coronado, 1963 **comb. restit.**
[Brazil, Peru, Guyana, Ecuador]

*Z. (C.) magnicaudelli* Mashimo, Engel, Dallai, Beutel and Machida, 2013 **comb. restit.**
[Peninsular Malaysia, Borneo]

*Z. (C.) manni* (Caudell, 1923)
[Bolivia, Peru]

*Z. (C.) mexicanus* (Bolívar y Pieltain, 1940)
[Mexico]

*Z. (C.) neotropicus* Silvestri, 1916
[Costa Rica]

*Z. (C.) novobritannicus* Terry and Whiting, 2012 **comb. restit.**
[Papua New Guinea]

*Z. (C.) philippinensis* Gurney, 1938 **comb. restit.**
[Philippines]

*Z. (C.) silvestrii* Karny, 1927 **comb. restit.**
[Indonesia: Mentawai Islands]

*Z. (C.) snyderi* (Caudell, 1920)
[USA; Jamaica]

*Z. (C.) weidneri* New, 1978 **comb. restit.**
[Brazil]

*Z. (C.) zimmermani* Gurney, 1939 **comb. restit.**
[Fiji]

SUBGENUS ***INCERTAE SEDIS***

*Z. congensis* van Ryn Tournel, 1971.
[Congo]

*Z. javanicus* Silvestri, 1913.
[Indonesia: Java]

*Z. juninensis* Engel, 2000.
[Peru]

*Z. lawrencei* New, 1995.
[Christmas Island]

*Z. leleupi* Weidner, 1976.
[Galapagos]

*Z. longicercatus* Caudell, 1927.
[Jamaica]

*Z. newi* (Chao & Chen, 2000).
[China: Taiwan]

*Z. sechellensis* Zompro, 2005.
[Seychelles]

*Z. swezeyi* Caudell, 1922.
[Hawaii]

#### Fossil calibrations

**1 Stem-Archaeognatha** (382.7 – 428.9 Ma), node 1

***1.1 Fossil taxon and specimen.*** Archaeognatha gen. spec. [329-AR4 (figured specimen): American Museum of Natural History, New York, USA]. Brown Mountain locality, Middle Devonian Panther Mountain Formation, near Gilboa, New York [2].

***1.2 Phylogenetic justification.*** The specimen is represented by a head fragment with large and dorsally adjacent eyes. This arrangement of the eyes is an autapomorphy of archaeognathans and is present in modern representatives [3]. The fragmentary nature of the specimen makes it difficult to determine if it represents a crown- or stem-group archaeognathan. We consider the latter as more likely since crown-archaeognathans are not known from the Palaeozoic (see 3.5). Consequently, it is most conservative to treat the Gilboa specimen as a stem archaeognathan, a placement that we follow here. Monophyly of the group is supported by molecular [4,5] and morphological data [6].

***1.3 Minimum age and justification.*** The arthropod-bearing horizon has been correlated with mid-Givetian [7]. The minimum age is obtained from the upper boundary of the Givetian [8].

***1.4 Soft maximum age and justification.*** The maximum age is taken from the two oldest Lagerstätten preserving terrestrial animals, the Přídolian Ludford Lane in Shropshire [9], England and the Pragian Rhynie chert from Aberdeenshire, Scotland (see 1.3). While both fossil deposits preserve trigonotarbids [10,11], an extinct order of pulmonate arachnids [12,13], alongside other land plants and arthropods, no jumping bristletails have been recovered from the deposits despite intensive sampling [9,14]. The Ludford Bone Bed was radiometrically dated to 420.0 ± 8.9 Ma [15]. We thus consider 428.9 Ma as a suitable maximum constraint on the node.

***1.5 Discussion.*** *Gaspea palaeoentognathae* (*nomen nudum*) reported from the Late Emsian Gaspe Bay fossil bed in Quebec, Canada by Labandeira et al. [16] is slightly older, but the exceptional three-dimensional preservation of the fossil has led Jeram et al. [9] to suggest that it may represent a modern contaminant. The separation between the eyes, is not apparent from the published images, which prevents us from determining if the specimen falls within crown-Archaeognatha. We therefore refrain from using this fossil as a calibration point in favour of the so-far-undisputed Gilboa specimen [17]. Arguments for why the Gilboa specimen we used as a calibration point likely does not represent a Recent contaminant have been discussed in detail by Shear et al. [7].

Besides the Devonian record of Archaeognatha, the extinct bristletail suborder Monura (treated as a separate order by some authors) including the single family Dasyleptidae ranges from the Carboniferous and is represented mainly by juveniles and shed exuvae [18–22]. The latest reliable monurans are from the Middle Triassic [23,24].

**2 Crown-Archaeognatha** (126.3 – 165 Ma), node 2

***2.1 Fossil taxon and specimen.*** *Cretaceomachilis libanensis* Sturm and Poinar, 1998 (Meinertellidae) [Nr. 194/35 (holotype): Milki collection, American University of Beirut, Beirut, Lebanon]. Lebanese amber, Jouar Es-Souss outcrop in Bkassine, south Lebanon [25,26].

***2.2 Phylogenetic justification.*** *Cretaceomachilis libanensis* is known from a single male specimen preserved in amber. It can be assigned to the extant family Meinertellidae based mainly on the dorsal basis of maxillary palps with a longitudinal projection and small abdominal sternites [27].

Specifically, Sturm and Poinar [25] suggested an affinity with the Recent *Machiloides*-group of genera [25]. Zhang et al. [27] recently described a congeneric from Burmese amber and revised the generic diagnosis of *Cretaceomachilis*, confirming its placement within Meinertellidae with a formal phylogenetic analysis. Together with other jumping bristletails from Lebanese amber, *C. libanensis* represents the earliest reliable crown-archaeognathan in the fossil record. The monophyly of Machilidae and Meinertellidae is supported by morphology [28] and a recent phylogenetic analysis of morphological characters incorporating fossil and extant taxa [27], but was questioned in a mitogenomic study [29], albeit with variable degrees of statistical support.

***2.3 Minimum age and justification.*** While Sturm and Poinar [49] did not initially indicate the precise origin of the amber, Azar [26] and Maksoud and Azar [30] later specified that the holotype originated from the Jouar Es-Souss outcrop in Bkassine in south Lebanon. The outcrop belongs to the 'lower interval' of the Lower Cretaceous "Grèsdu Liban" that Maksoud et al. conservatively attributed to the Early Barremian based on fossil evidence [31]. We thus use the upper boundary of the Barremian, 126.3 Ma, as the minimum age constraint on the node [8].

***2.4 Soft maximum age and justification.*** Crown-archaeognathans are absent from the Late Jurassic Daohugou beds. Ar/Ar and SHRIMP U-Pb dating indicated that the age of the intermediate-acid volcanic rock overlaying the Daohugou fossiliferous beds is approximately 164–165 Ma, and so the age of the fossil deposit is older than or equal to 165 Ma, corresponding to the Callovian of the Middle Jurassic [32], an age widely adopted by palaeoentomologists [33]. Indeed, the fossil insects from the Daohugou beds may be correlated to those from the slightly younger deposit in Karatau (Kazakhstan) [34], but there is strong evidence indicating that the age of the Daohugou beds may be older [35]. As such, 165 Ma provides a conservative maximum age constraint on the node.

***2.5 Discussion.*** Crown-Archaeognatha consists of two extant families grouped into the suborder Machilida that are represented in the Recent fauna by some 500 described species. The Machilidae are found mostly in the northern hemisphere, while the Meinertellidae are predominantly distributed in the southern hemisphere [28,36]. While members of the stem-archaeognathan lineage Monura were diverse in the Palaeozoic and into the Triassic [23], the fossil record of Machilida is confined to the Mesozoic and Cenozoic, and is known mostly from amber inclusions. Montagna et al. [37,38] recently described a putative crown-archaeognathan (Machilidae) from the Middle Triassic of Italy, but the fossil in question is in fact a misidentified mayfly nymph. Thus, the earliest reliable record of crown Archaeognatha is from Lebanese amber. The archaeognathan fauna from Lebanese amber is already diverse and includes four species in three genera quite similar to modern taxa [25,39].

**3 Crown-Zygentoma** (112.6 – 428.9 Ma), node 3

***3.1 Fossil taxon and specimen.*** Lepismatidae gen. spec. [B99 (figured specimen): Naturmuseum Senckenberg, Frankfurt am Main, Germany; 1998 III/4 (figured specimen): Bayerischen Staatssammlung für Paläontologie und historische Geologie, Munich, Germany; SMNS66535 (figured specimen): State Museum of Natural History Stuttgart, Stuttgart, Germany]. Nova Olinda quarry, Nova Olinda Member of the Lower Cretaceous (Aptian) Crato Formation, northeastern Brazil [40,41].

***3.2 Phylogenetic justification.*** The fossil can be assigned to Zygentoma, based on the compound eyes widely separated; flattened ovoid coxae; the absence of wings; and three long terminal filaments lacking scales [3,42]. The presence of unreduced compound eyes represents a plesiomorphic character that is also present in the family Lepidotrichidae (sometime divided into two families: the fossil Lepidotrichidae, and the extant Tricholepidiidae) [3] , implying that the fossil belongs to crown Zygentoma.

***3.3 Minimum age and justification.*** An Aptian age of the Crato Formation is supported by the ostracod fauna [43] and pollen [44–46]. The Nova Olinda Member is presently believed to lie at the Aptian/Albian boundary [41]. The minimum age thus comes from the upper boundary of the Aptian [8].

***3.4 Soft maximum age and justification.*** As in 1.4.

**4 Stem-Odonata** (315 – 428.9 Ma), node 4

***4.1 Fossil taxon and specimen.*** *Oligotypus huangheensis* Ren, Nel, and Prokop, 2008 (†Meganeuridae) [CNU-NX2006003 (holotype); CNUNX1-433;  CNU-NX1-435; CNU-NX1-453; CNU-NX1-407; CNUNX1-431; CNU-NX1-432; CNU-NX1-434; CNU-NX1-436; CNU-NX1-455: Key Lab of Insect Evolution and Environmental Changes, College of Life Sciences, Capital Normal University, Beijing, China]. Xiaheyan village locality, Tupo Formation, Zhongwei County, northwestern China [47,48].

***4.2 Phylogenetic justification.*** *Oligotypus huangheensis* is known from wing fragments of varying degrees of preservation. The fusion of the veins CuP and CuA to a single oblique vein, distinctly stronger than the crossveins supports its placement in Meganisoptera [47]. Meganisoptera (‘Protodonata’ or ‘griffenflies’) is an extinct lineage most likely forming a stem group to true Odonata [49–51]. They differ from crown-odonates by the absence of the nodus and pterostigma on the wings, males lacking secondary genitalia, and sometimes very large wingspans [52]. Monophyly of Odonata, as well as of Anisozygoptera, Anisoptera, and Zygoptera, is supported by molecular analyses [5,53], and a total-evidence phylogenetic study [54].

***4.3 Minimum age and justification.*** The fossiliferous horizon yielding the specimen has been dated to the Namurian B/C or to the Bashkirian (latest Duckmantian), based on the presence of a characteristic ammonoid and conodont fauna [55,56]. Trümper et al. [56] proposed an upper bound on the age of the insect-bearing bed of ~315 Ma, which we follow herein.

***4.4 Soft maximum age and justification.*** As in 1.4.

**5 Crown-Odonata** (228.5 – 314.6 Ma), node 5

***5.1 Fossil taxon and specimen.*** *Triassolestodes asiaticus* Pritykina, 1981 (†Triassolestidae) [PIN2240/1783 (holotype): Paleontological Institute of the Academy of Science of Russia, Moscow, Russia]. Dzhayloucho, Upper Triassic, Ladinian-Carnian Madygen Formation, south of Fergana Valley, Kyrgyzstan [57].

***5.2 Phylogenetic justification.*** The fossil can be assigned to *Triassolestodes* on the basis of possessing the following autapomorphic characters: fusion of AA to MP + Cu; and the presence of a large posteriorly opened and transverse subdiscoidal space [58,59]. A phylogenetic analysis supported the fossil’s placement within the Isophlebioptera-Triassolestidae clade [59]. A family-level phylogenetic analysis recovered Triassolestidae within crown-group Odonata [54], and Kohli et al. [51] recommended *T. asiaticus* as a calibration point for crown-Odonata (Zygoptera + Epiprocta).

***5.3 Minimum age and justification.*** Kohli et al. [51] assigned the fossil to the Ladinian based on the flora found at the locality [60]. The minimum age is based on the base of the Carnian which has been dated to 228.5.0 Ma [8].

***5.4 Soft maximum age and justification.*** The maximum age constrain for the node is provided by the age of Mazon Creek in Illinois, USA. Mazon Creek [61]. None of the younger well-explored late Palaeozoic Lagerstätten, including Commentry in France, Midco in Oklahoma, Elmo in Kansas, Obora in Czechia, or Tshekarda in Russia, have yielded crown odonates, so we consider the maximum age of the Mazon Creek fauna as an appropriate soft maximum constraint.

**6 Crown-Ansioptera** (162.5 – 314.6 Ma), node 6

***6.1 Fossil taxon and specimen.*** *Sinacymatophlebia mongolica* Nel and Huang, 2009 (†Cymatophlebiidae) [NIGP 148312 (holotype); NIGP 148313 (paratype): Nanjing Institute of Geology and Palaeontology, Nanjing, China]. Locality near the Daohugou Village, Wuhua Township, Ningcheng County, Chifeng City, Middle Jurassic Haifanggou Formation, Inner Mongolia, northeast China [65].

***6.2 Phylogenetic justification.*** The fossil, known from a male hind wing and a fragment of the thorax and abdomen, can be assigned to crown-Anisoptera based on the presence of several wing venation synapomorphies: vein Rsp1 present; RP1 and RP2 basally parallel up to the pterostigma; and RP3/4 and MA undulating [51,65,66].

***6.3 Minimum age and justification.*** The fossil was collected from the Daohugou beds of the Haifanggou Formation, Inner Mongolia, northeastern China. The precise age of the Daohugou beds has been controversial [33,34,67,68]. 39Ar–40Ar and SHRIMP U-Pb dating has provided an age of 165 ± 2.5 Ma of the acid volcanic rock overlaying the Daohugou fossiliferous horizon yielding insects [32,69]. This corresponds to the Callovian of the Middle Jurassic and is further corroborated by the composition of the fossil insect community, which is similar to the slightly younger deposit in Karatau, Kazakhstan [34]. This provides a minimum age of 162.5 Ma.

***6.4 Soft maximum age and justification.*** As in 5.4.

**7 Crown-Siphlonuridae** (240.5 – 314.6 Ma), node 7

***7.1 Fossil taxon and specimen.*** *Triassonurus doliiformis* Sinitshenkova and Papier, 2005 (Siphlonuridae) [9304 (holotype): Louis Grauvogel collection, Ringendorf, Bas-Rhin, France]. Arzviller, Upper Buntsandstein Moselle, France [70].

***7.2 Phylogenetic justification.*** Affinity of this incomplete nymph with the extant family Siphlonuridae is suggested by the following combination of characters [70,71]: body large and non-flattened; head longer than prothorax; mesothorax massive and with significantly shorter metathorax; forewing pads large, wide, and almost completely covering the hind ones; legs short and slender; abdominal segments lacking sharp denticles; tergalia large and rounded; cerci and paracercus long. A total evidence phylogenetic study incorporating 5 genes and 101 morphological characters recovered the family Siphlonuridae as monophyletic [72]. We use the fossil to calibrate crown Schistonota, represented in our study by *Ephemera danica*.

***7.3 Minimum age and justification.*** The Grès à Meules unit of the Grès-a-Voltzia Formation belongs to the last stage of the fluviatile facies present in the Buntsandstein [73]. Bourquin et al. [74,75] correlated the Grès-a-Voltzia Formation to the middle Anisian stage based on sequence stratigraphy. The uppermost boundary of the Anisian is 241.5 ± 1 Ma [8], giving a minimum age of 240.5 Ma.

***7.4 Soft maximum age and justification.*** As in 5.4.

***7.5 Discussion.*** *Triassonurus* is the earliest representative of the extant family Siphlonuridae [76], sharing conspicuous similarities with larvae of Recent members of the family [77], and also of crown Ephemeroptera.

**8 Stem-Plecoptera** (315 – 428.9 Ma), node 8

***8.1 Fossil taxon and specimen.*** *Gulou carpenteri* Béthoux, Cui, Kondratieff, Stark, and Ren, 2011 (†Gulouidae) [CNU-NX1-143 (holotype); CNU-NX1-137 to CNU-NX1-142; CNU-NX1-144 to CNU-NX1-159: Key Lab of Insect Evolution and Environmental Changes, College of Life Science, Capital Normal University, Beijing, China]. Xiaheyan village, Zhongwei city, Early Pennsylvanian Tupo Formation, Ningxia Hui Autonomous Region, China [78,79].

***8.2 Phylogenetic justification.*** Béthoux et al. [78] originally placed *Gulou* into a new family, Gulouidae, within stem Plecoptera. This was based on the basal origin of the RP vein; narrow area between RA and RP for a long distance; and CuA with a few extremely distal branches. However, *Gulou* lacks the ra-rp specialized cross-vein and possesses a branched MP, which are not present in any extant stoneflies, and shares other morphological similarities with Permian stem-Plecoptera. The placement of *Gulou* within stem-Plecoptera was questioned by Aristov [80] who synonymized Gulouidae with Emphylopteridae (Cnemidolestodea). This was corrected by Schubnel et al. [79], who pointed out several morphological incongruences regarding the shape of the veins CuA and CuP in Aristov's treatment, and moved Gulouidae back to stem-Plecoptera.

***8.3 Minimum age and justification.*** As in 4.3.

***8.4 Soft maximum age and justification.*** As in 1.4.

**9 Crown-Plecoptera** (174.2 – 241.5 Ma), node 9

***9.1 Fossil taxon and specimen.*** *Dobbertiniopteryx capniomimus* Ansorge, 1993 (Capniidae) [MfN LDA 740 (holotype): Museum für Naturkunde, Berlin, Germany]. Closed clay pit near Dobbertin, Mecklenburg, Germany [81,82].

***9.2 Phylogenetic justification.*** Known from an isolated forewing, *D. capniomimus* can be assigned to the extant family Capniidae based on the presence of diagnostic wing venation characters [81,82]. Sinitshenkova [83] questioned the assignment of *Dobbertiniopteryx* to Capniidae and considered an alternative placement within the extinct family Baleyopterygidae. However, a more recent discovery of an articulated *Dobbertiniopteryx* body fossils from the younger Daohugou Biota has confirmed the placement of the genus in Capniidae [84].

***9.3 Minimum age and justification.*** The fossil has been found in carbonate concretions within clay of the “Green Series” belonging to the *Harpoceras exaratum* subzone of the *Harpoceras falciferum* ammonite zone, which corresponds to the Lower Toarcian [85]. The minimum age is thus taken from the upper boundary of the Toarcian, 174.2 Ma [8].

***9.4 Soft maximum age and justification.*** We use the maximum age of the Ladinian-Carnian Madygen Formation in Kyrgyzstan, the world’s richest Triassic insect Konservat-Lagerstätte [60]. No crown group plecopterans are known from the deposit.

***9.5 Discussion.*** While some Permian fossils have been assigned to crown-Plecoptera, these require re-examination. At present, *D. capniomimus* is the earliest reliable crown stonefly [86].

**10 Crown-Dermaptera** (126.3 – 167.5 Ma), node 10

***10.1 Fossil taxon and specimen.*** *Rhadinolabis phoenicica* Engel, Ortega-Blanco, and Azar, 2011 (family *incertae sedis*) [1013 (holotype); 1018 (paratype): Muséum national d'Histoire naturelle (National Museum of Natural History), Paris, France]. Lebanese amber, Hammana-Mdeyrij outcrop, Caza Baabda, Mouhafazit Jabal Loubnan, Lebanon [87].

***10.2 Phylogenetic justification.*** *Rhadinolabis phoenicica*, known from amber inclusions, can be assigned to Neodermaptera based on its trimerous tarsi and absence of a well-developed ovipositor. While its familiar affinity remains uncertain due to insufficient preservation of key characters, it is excluded from Eudermaptera on the basis of the second tarsomere not notably expanded and projecting strongly beneath the third tarsomere [87]. We thus conservatively use *Rhadinolabis* to calibrate the node representing crown-Dermaptera.

***10.3 Minimum age and justification.*** The Hammana-Mdeyrij outcrop belongs to the ‘upper interval’ of the Grès du Liban [30]. The upper interval was assigned to the Late Barremian based on fossil evidence [31]. The upper boundary of the Barremian, 126.3 Ma [8], is thus used as the minimum constraint on the node.

***10.4 Soft maximum age and justification.*** The maximum constrain on the node is provided by the age of the Daohugou biota in Inner Mongolia, northeastern China [32,69].

***10.5 Discussion.*** The earliest stem-dermapterans appear in the fossil record during the Late Triassic [88,89]. The Palaeozoic extinct order Protelytroptera may also include stem-dermapterans [90–93]. All Jurassic earwigs belong to extinct suborders [94,95]. Zhao *et al.* considered the enigmatic mid-Jurassic genus *Atopderma* as a likely neodermapteran [96], although the fragmentary nature of the specimens renders their assignment uncertain [97]. The earliest, definitive Neodermaptera are from the earliest Cretaceous. Rasnitsyn and Quicke [98] figured a putative neodermapteran from the Lower Cretaceous Zaza Formation in Transbaikalian Russia (~134.7 Ma), slightly older than *Rhadinolabis*. By the mid-Cretaceous, crown-Dermaptera evidently became diverse as evidenced by their representatives known from Burmese, French, and Spanish ambers [99].

**11 Stem-Orthoptera** (315 – 428.9 Ma), node 11

***11.1 Fossil taxon and specimen.*** *Xixia huban* Gu, Béthoux, and Ren, 2013 (order Cnemidolestodea, family *incertae sedis*) [CNU-NX1-381 (holotype); CNUNX1- 380–392: Key Lab of Insect Evolution and Environmental Changes, College of Life Science, Capital Normal University, Beijing, China].

Xiaheyan village, Zhongwei city, Early Pennsylvanian Tupo Formation, Ningxia Hui Autonomous Region, China [100].

***11.2 Phylogenetic justification.*** *Xixia huban*, known from isolated fore- and hind wing fragments, can be placed into the superorder Archaeorthoptera on the basis of possessing the characteristic fusion of CuA with the with the anterior branch of CuP [101,102]. It can further be placed into Cnemidolestodea, as indicated by the the ScP reaching RA; and MP diverging obliquely from M and reaching the stem of CuA + CuPa [103]. Since Archaeorthoptera has been regarded as a stem group to Orthoptera based on wing venation characters [101], we use the fossil to calibrate the stem of Orthoptera. Monophyly of Orthoptera is supported by molecular data [5,104], morphology [103], and a combination of both [105].

***11.3 Minimum age and justification.*** As in 4.3.

***11.4 Soft maximum age and justification.*** As in 1.4.

**12 Stem-Ensifera** (280.0 – 314.6 Ma), node 12

***12.1 Fossil taxon and specimen.*** *Raphogla rubra* Béthoux, Nel, Lapeyrie, Gand, and Galtier, 2002 (Raphoglidae) [Ld LAP 415ab (holotype): Lapeyrie collection, Musée Fleury, Lodève, France]. Locality F21 D, “Le Moural D”, Salagou Formation, Lodève Basin, Hérault, France [106].

***12.2 Phylogenetic justification.*** This isolated wing can be placed among Enserifa based on wing venation characters [106]. Affinity with Gryllidea and Tettigoniidea is suggested by the notably wide area between the anterior margin and Sc; RS moderately long basal of a short fusion with the anterior branch MA1a of MA; and MP + CuA1 with only one simple anterior branch [106]. Given that *Raphogla* probably represents a sister group to Grylloidea and Tettigonioidea [106], it has been used to calibrate stem Enserifa.

***12.3 Minimum age and justification.*** The Octon Member that underlies the Merifons Member where the fossil was found has been U–Pb zircon dated to the Artinskian (284 ± 4 Ma) [107]. This gives a conservative minimum age of 280.0 Ma.

***12.4 Soft maximum age and justification.*** As in 5.4.

**13 Crown-Hagloidea** (191.4 – 241.5 Ma), node 13

***13.1 Fossil taxon and specimen.*** *Aboilus tuzigouensis* Lin and Huang, 2006 (Prophalangopsidae) [NIGP 133703 (holotype): Nanjing Institute of Geology and Palaeontology, Nanjing, China]. Tuzigou locality, Lower Sinemurian Badaowan Formation, Xinjiang Province, northwestern China [108].

***13.2 Phylogenetic justification.*** *Aboilus tuzigouensis,* known from an isolated tegmina, can be assigned to the family Prophalangopsidae based on the widened anal part of the forewing and presence of a long secondary longitudinal vein connecting MP+CuA1 with CuA2 [109]. While its placement in the subfamily Aboilinae has been questioned [110], it can nonetheless still be used to calibrate the node representing crown Hagloidea.

***13.3 Minimum age and justification.*** The Badaowan Formation has been biostratigraphically constrained to the latest Rhaetian–Sinemurian [111]. The upper boundary of the Sinemurian, 191.4 Ma [8], thus provides a conservative minimum age constrain one the node.

***13.4 Soft maximum age and justification.*** As in 10.4.

**14 Crown-Caelifera** (249.8 – 295.0 Ma), node 14

***15.1 Fossil taxon and specimen.*** *Praelocustopsis mirabilis* Sharov, 1968 (†Locustavidae) [PIN 2010/2 (holotype): Paleontological Institute of the Academy of Science of Russia, Moscow, Russia]. Bugarikta locality, at the right bank of the Lower Tunguska, Bugarikta Formation, Krasnoyarsk Krai, Russia [112].

***14.2 Phylogenetic justification.*** *Praelocustopsis mirabilis,* is known from a pair of fore- and hind wings. It placement in the extinct family Locustavidae is supported by the long and narrow forewing with a well-developed costal vein and reduced anal field and the cross-connection from M to CuA (= stem of MP) transverse [112–114]. The extinct Locustavidae have been regarded as close to the extant early-diverging caeliferan family Eumastacidae [112,115]. We conservatively use the fossil to calibrate the node representing crown Caelifera.

***14.3 Minimum age and justification.*** The dating of the Bugarikta Formation remains contentious. Sharov [112] regarded the insect-bearing horizon as Lower Triassic but did not provide closer stratigraphic details. Ostracod, conchostracan, and fish faunas point towards an Early Triassic age, while palynomorphs supports a Late Permian age [116]. The presence of the ray-finned fish genus *Eoperleidus* seems to suggest that the Formation may lie close to the Permian-Triassic boundary [117]. We conservatively use the uppermost boundary of the lowermost Triassic stage, the Induan at 249.8 Ma [8], as the minimum constraint in the node.

***14.4 Soft maximum age and justification.*** The maximum age constraint is based on the age of the Sakmarian–Artinskian locality Obora in Moravia, Czech Republic [118]. Together with Elmo in Kansas and Tshekarda in Russia, these three localities represent the best-explored and most productive insect Lagerstätten of the Permian. The lower boundary of the Sakmarian is 295.0 Ma [8].

***14.5 Discussion.*** *P. mirabilis* represents the earliest uncontested member of the family Locustavidae, and thus the earliest caeliferan. The older *Legendreia magnifica* from the middle Permian Yinping Formation of China (~259 Ma) has been suggested by Béthoux to represent a stem-caeliferan [119], but see Huang et al. [120] for an opposing view.

**15 Stem-Dictyoptera (315 – 428.9 Ma), node 15**

***15.1 Fossil taxon and specimen.*** *Qilianiblatta namurensis* Zhang, Schneider, and Hong, 2012 (Dictyoptera, family indet.) [GMCB 04GNX1001-1 (holotype); GMCB 97×128: Geological Museum of China, Beijing, China; CNU-NX1-303: Key Lab of Insect Evolution and Environmental Changes, College of Life Sciences, Capital Normal University, Beijing, China]. Xiaheyan village locality, Tupo Formation, Zhongwei County, northwestern China [55,121].

***15.2 Phylogenetic justification.*** The preserved right forewing shares with extant Blattodea a deeply concave CuP vein [122], but the RA with branches translocated to RP indicates that the fossil belongs to stem Dictyoptera [121]. In line with Wolfe et al. [71], we conservatively assign the fossil to stem Dictyoptera, and thereby among crown-Polyneoptera [122–124].

***15.3 Minimum age and justification.*** As in 4.3.

***15.4 Soft maximum age and justification.*** As in 1.4.

***15.5 Discussion.*** 'Roachoids' such as *Q. namurensis* were roach-like insects abundant in the Palaeozoic and some of the earliest winged insects in the fossil record. While some authors consider Palaeozoic roachoids as close to extant Blattodea [125], they are excluded from crown Blattodea and Mantodea by wing venation character and most notably by the presence of long external ovipositors in females [126]. This therefore also makes *Q. namurensis* suitable for calibrating the node representing stem-Dictyoptera (node 17).

A slightly older but less well preserved fossil assigned to Archaeorthoptera has been reported from the Lowermost Namurian of the Czech Republic [127], dating back to 328.48 ± 0.19 Ma [128]. Misof et al. [5] considered this fossil to represent a stem-Pterygota, without providing a justification, while Wolfe et al. [71] considered the fossil as too poorly preserved for use as a calibration point. Dvořák et al. [129] recently re-examined the specimen and found that it probably is not an insect at all, but most likely a fragment of a fish fin.

**16 Stem-Grylloblattodea** (315 – 428.9 Ma), node 16

***16.1 Fossil taxon and specimen.*** *Sinonamuropteris ningxiaensis* Peng, Hong, and Zhang, 2005 (†Sinonamuropteridae) [91NZ1/035 (holotype): Geological Museum of China, Beijing, China; CNU-NX1-161 (neotype): Key Lab of Insect Evolution & Environmental Changes, College of Life Sciences, Capital Normal University, Beijing, China]. Xiaheyan village locality, Tupo Formation, Zhongwei County, northwestern China [130–132].

***16.2 Phylogenetic justification.*** The family Sinonamuropteridae was described by Peng et al. [130] with four genera and nine species and referred to the extinct order Diaphanopterodea. Based on new specimens preserving compete wings from the type locality, Cui et al. [132] synonymised the nine original species under *S. ningxiaensis*. Wing venation of Sinonamuropteridae indicates that the family belongs to Notoptera (Grylloblattodea + Mantophasmatodea); the wings possess of a CuA forked into two main stems (CuA1 and CuA2); and a m-cua arculus in the forewings [132]. It has been compared to the uncontested Permian notopteran *Chelopterum peregrinum* that also has completely preserved wings [132]. *Sinonamuropteris* represents a stem group to Grylloblattodea + Mantophasmatodea since modern ice crawlers and rock crawlers lack wings [3]. We thus use *Sinonamuropteris* to calibrate stem-Grylloblattodea + Mantophasmatodea. The monophyly of Grylloblattidae is supported by molecular analyses [5,133] as well as morphological characters [3].

***16.3 Minimum age and justification.*** As in 4.3.

***16.4 Soft maximum age and justification.*** As in 1.4.

**17 Crown-Mantophasmatodea** (162.5 – 314.6 Ma), node 17

***17.1 Fossil taxon and specimen.*** *Juramantophasma sinica* Huang, Nel, Zompro, and Waller, 2008 (Mantophasmatidae) [NIGP 142171a, b (holotype): Nanjing Institute of Geology and Palaeontology,

Chinese Academy of Sciences, Nanjing, China]. Locality near the Daohugou Village, Wuhua Township, Ningcheng County, Chifeng City, Middle Jurassic Haifanggou Formation, Inner Mongolia, north-east China [134].

***17.2 Phylogenetic justification.*** *Juramantophasma sinica* is known from a dorsoventral compression fossil preserving an adult female. It displays several apomorphies of Mantophasmatidae, namely the third tarsomere with a sclerotized elongated dorsal process; enlarged and fan-like pretarsal arolia with a clearly visible row of dorsal setae; last tarsomere connected to the penultimate one at a right angle; and female gonoplacs short and claw-shaped [134,135]. The fossil differs from extant Mantophasmatidae by the absence of ventroapical spines on the tibiae, which are characteristic of Mantophasmatinae [136]. It can however be placed into the extinct subfamily †Raptophasmatinae known otherwise only from Baltic amber [136,137]. This makes *Juramantophasma* a member of crown-group Mantophasmatodea. The monophyly of Mantophasmatodea is supported by a molecular phylogeny based on 1,300 bp of mitochondrial DNA [138] and by a total evidence phylogenetic study based on three genes and 125 morphological characters [105].

***17.3 Minimum age and justification.*** As in 6.3.

***17.4 Soft maximum age and justification.*** As in 5.4.

**18 Stem-Phasmatodea** (162.5 – 295.0 Ma), node 18

***18.1 Fossil taxon and specimen.*** *Adjacivena rasnitsyni* Shang, Béthoux, and Ren, 2011 (†Susumaniidae) [CNU-PHA-NN2009001 (holotype); CNU-PHA-NN2009002 (paratype): Key Lab of Insect Evolution and Environmental Changes, College of Life Sciences, Capital Normal University, Beijing, China]. Locality near the Daohugou Village, Wuhua Township, Ningcheng County, Chifeng City, Middle Jurassic Haifanggou Formation, Inner Mongolia, north-east China [139].

***18.2 Phylogenetic justification.*** *Adjacivena rasnitsyni*, known from compression fossils of a male and a female preserving the wings and genitalia, displays the following apomorphies of Phasmatodea: ovipositor concealed by an operculum (female specimens); and tergum 10 bearing a hook (vomer) on the venter (male specimen) [140–142]. It is placed outside of crown Phasmatodea by the following combination of wing characters: narrow area between MA2 and MP + CuA1 present; MA2 approaching MP + CuA1 a short distance before the middle of the forewing; MP + CuA1 forked in the forewing; MA1 fused over a moderate distance with RP in the hind wing; MA2 fused with MA1 distal to its divergence from RP + MA1 in the hind wing [139]. The phylogenetic position of *A. rasnitsyni* has recently been supported by a formal phylogenetic analysis [160]. The monophyly of Phasmatodea is strongly supported by transcriptome data [5,143], ribosomal and H3 sequences [105], as well as morphological characters [144,145].

***18.3 Minimum age and justification.*** As in 6.3.

***18.4 Soft maximum age and justification.*** As in 14.4.

***18.5 Discussion.*** The early evolution of Phasmatodea lies in murky waters; although a considerable number of putative stem-phasmatodeans have been reported from the Permian and Mesozoic by workers from the 1960s until the turn of the millennium [98,112,146,147], mostly on the basis of fragmentary wings. Whether these fossils represent true relatives of modern stick and leaf insects has been questioned [145,148]. Tilgner [149] reviewed the fossil record of phasmatodeans and concluded that all the putative stem-group fossils from the Palaeozoic and Mesozoic reported at the time were diagnosed based on wing venation characters that are not unique to the order Phasmatodea and are plesiomorphic. It was not until recently that some members of the extinct family †Susumaniidae were recognised as true stem-phasmatodeans. Specimens from the Middle Jurassic Jiulongshan [139] and Early Cretaceous Yixian Formation in China [150,151] were described with preserved vomers, male clasping genital structures that represent an apomorphy of Phasmatodea [140–142]. These fossils can thus be identified as the earlier stem-phasmatodeans in the fossil record. Other specimens assigned to †Susumaniidae are known, extending up to the Eocene [152], but because the fossil family is not defined by any definitive autapomorphy [145], these fossils should be treated with caution.

Another stem-phasmatodean reported recently from the Haifanggou Formation, *Aclistophasma echinulatum* [153], would be equally suited to calibrate the node.

**19 Crown-Embiodea (Clothodidae)** (98.17 – 167.5 Ma), node 19

***19.1 Fossil taxon and specimen.*** *Atmetoclothoda orthotenes* Engel and Huang, 2016 (Clothodidae) [HP-B-4164, NIGP 162534 (holotype): Nanjing Institute of Geology and Palaeontology, Nanjing, China]. Burmese amber, Hukawng Valley, Myitkyina District, Kachin State, northern Myanmar [154].

***19.2 Phylogenetic justification.*** *Atmetoclothoda orthotenes*, known from a male preserved as an amber inclusion, belongs to the extant family Clothodidae as indicated by the MA forked; CuA multibranched; extensive crossvenation; undivided tenth tergum; completely symmetrical male terminalia; and the elongate and completely symmetrical cercomeres [154]. It can thus be used to calibrate the node separating Clothodidae from the remaining webspinners.

***19.3 Minimum age and justification.*** The fossil is preserved in Burmese amber mined in the Hukawng Valley in northern Myanmar. Volcanoclastic matrix from the amber-bearing horizon was dated radiometrically at 98.79 ± 0.62 Ma [155], which is in line with the age predicted based on palaeontological evidence [156]. However, the zircon date should be taken as a lower limit for the age of the amber [157] since the method made no use of chemical abrasion which typically results into younger ages [158,159]. A reliable upper limit on the age of Burmese amber is provided by a juvenile *Puzosia* ammonite trapped in the amber, which indicates that the deposit is at most late Albian in age [160]. Here we use 98.17 Ma as the conservative minimum age of Burmese amber.

***19.4 Soft maximum age and justification.*** As in 10.4.

***19.5 Discussion.*** The fossil record of crown-group webspinners starts in the Cretaceous. Other embiodeans from Burmese amber [reviewed in 175] are equally suited for calibrating the node representing crown-Embiodea.

**20 Crown-Embiodea (Oligotomidae)** (98.17 – 167.5 Ma), node 20

***20.1 Fossil taxon and specimen.*** *Litoclostes delicatus* Engel and Huang, 2016 (Oligotomidae) [HP-B-4164, NIGP 162535 (holotype): Nanjing Institute of Geology and Palaeontology, Nanjing, China]. Burmese amber, Hukawng Valley, Myitkyina District, Kachin State, northern Myanmar [154].

***20.2 Phylogenetic justification.*** *Litoclostes delicatus*, known from a male preserved as an amber inclusion, is attributed to the extant family Oligotomidae based on its rather simplified and weak venation with MA unforked; submedian and subapical metabasitarsal plantulae present; simple left basal cercomere; and echinulation on the basal cercomeres absent [154]. It can thus be used to calibrate separating Oligotomidae from the remaining webspinners.

***20.3 Minimum age and justification.*** As in 19.3.

***20.4 Soft maximum age and justification.*** As in 10.4.

**21 Crown-Anisomorphini (Pseudophasmatidae)** (43.47 Ma – 130.8 Ma), node 21

***21.1 Fossil taxon and specimen.*** *Eophasmodes oregonense* Clark-Sellick, 1994 (ichnotaxon) [UF 15768-6437 (holotype); UF 15768-6364; 15768-6366; 225-8686 (paratypes): Florida Museum of Natural History, University of Florida, Gainesville, Florida, USA]. John Day National Fossil Monument Nut Beds, Eocene Clarno Formation, Oregon, USA [162] (Clark-Sellick 1994).

***21.2 Phylogenetic justification.*** The eggs possess a distinct operculum and micropylar plate, features that are apparently unique to the eggs of Phasmatodea [163]. The operculum is tilted ventrally, a character which is known only in Anisomorphini [162]. The fossil is thus used to calibrate the Anisomorphini – Pseudophasmatini split recovered by analyses of transcriptome data [143,164].

***21.3 Minimum age and justification.*** Radiometric dating has yielded a range of ages for the Clarno Nut Bed ranging from 43.76 ± 0.29 Ma to 48.32 ± 0.11 Ma [165–167] and this age is further corroborated by the mammal fauna known from the locality [168]. This provides a minimum age for the node, 43.47 Ma.

***21.4 Soft maximum age and justification.*** The maximum constraint on the clade is provided by the maximum age of Lebanese amber, that has been dated as late Barremian to early Aptian [31]. This is because none of the Cenozoic and Early Cretaceous Lagerstätten (e.g., Baltic amber, Messel pit, Canadian amber, Taimyr amber, New Jersey amber, Burmese amber, French amber, Spanish amber, Crato Formation, Yixian Formation) preserve any members of the group.

**22 Crown-Lonchodidae** (98.17 – 167.5 Ma), node 22

***22.1 Fossil taxon and specimen.*** *Echinosomiscus primoticus* Engel and Wang, 2016 (family incertae sedis) [NIGP 163536 (holotype); CNU-PHA-MA2017001 (paratype): Nanjing Institute of Geology and Palaeontology, Chinese Academy of Sciences, Nanjing, China]. Burmese amber, Hukawng Valley, Myitkyina District, Kachin State, northern Myanmar [148].

***22.2 Phylogenetic justification.*** *Echinosomiscus primoticus*, known from a male specimen preserved as an amber inclusion, can be placed into Euphasmatodea on the basis of the following apomorphies: having the abdominal sternum I fused with the metasternum; thorn field present on the abdominal tergum 10; and pentamerous tarsi [141,169]. It can be placed among the 'areolate lineages' on the basis of lacking the area apicalis on the tibiae. It shares with extant Lonchodinae the slender body form and long antennae [170]. Crucially, the tenth abdominal tergite is divided into moveable hemitergites, a character that has traditionally been considered as the hallmark of Lonchodinae [171–175], although it is absent in the ambiguous taxon *Neohirasea* [141]. Extant Lonchodinae are restricted to Southeast Asia and Australia, which is notable since most of the fauna and flora of Burmese amber is apparently of an Australian origin [176]. Admittedly, a similar morphology of the 10th abdominal tergum is also found in Clitumninae [141], but the fossil differs from this subfamily in having the profemora trapezoid in cross-section, not triangular [170]. Moreover, a morphological phylogenetic analysis recovered *Echinosomiscus* as a sister group to the lonchodine taxon *Eurycantha*, albeit with a rather limited taxon sampling [177]. *Echinosomiscus primoticus* falls outside of stem-Lonchodinae, it differs in the structure of the antennae, head, and abdomen and as such has been placed into its own subfamily †Echinosomiscinae [148]. We thus provisionally treat the fossil as a sister group to Lonchodinae. The monophyly of Lonchodinae, although controversial in the past, is well-supported by the latest transcriptome analyses [143,164].

***22.3 Minimum age and justification.*** As in 19.3.

***22.4 Soft maximum age and justification.*** As in 10.4.

***22.5 Discussion.*** Other members of Euphasmatodea [178] and even supposed euphasmatodean eggs [179] have been reported from Burmese amber, although their exact phylogenetic position remains uncertain [177]. These mid-Mesozoic stick insects together show that Euphasmatodea began to diversify by 100 Ma. The plant-seed genus *Knoblochia* known from the Late Cretaceous of central Europe has been recently reinterpreted as possible phasmatodean eggs [180], but this assignment seems unlikely as the supposed eggs lack a distinct micropylar plate [163] that is characteristic of the order.

**23 Crown-Dictyoptera** (228.5 Ma – 295.0 Ma), node 23

***23.1 Fossil taxon and specimen.*** *Oothecichnus duraznensis* Cariglino, Lara, and Zavattieri, 2020 (ichnotaxon) [IANIGLA-PI 3130 (holotype); IANIGLA-PI 3129, IANIGLA-PI 3131, IANIGLA-PI 3132, IANIGLA-PI 3133, IANIGLA-PI 3136, IANIGLA-PI 3137, IANIGLA-PI 3138 (paratypes): Instituto Argentino de Nivología, Glaciología y Ciencias Ambientales, Mendoza, Argentina]. del Durazno locality, Mendoza, Triassic Potrerillos Formation, Quebrada, Argentina [181].

***23.2 Phylogenetic justification.*** The identity of the fossils as oothecae (egg cases) is confirmed by their general morphology and well as ultrastructural and chemical analysis [181]. Oothecae are a feature unique to crown Dictyoptera [182,183]. It is likely that stem dictyopterans, namely Palaeozoic roachoids, did not produce oothecae, as the females in these taxa had elongate external ovipositors (see 9.5). The fossils are thus suitable for calibrating the node representing crown-Dictyoptera.

***23.3 Minimum age and justification.*** The fossils originate from the uppermost strata of the Potrerillos Formation in Mendoza, Argentina. U\Pb SHRIMP dating of the tuff beds from the base of the Potrerillos Formation yielded ages of 239.2 ± 4.5 Ma and 239.7 ± 2.2 Ma ages [184]. A Carnian age of the insect-bearing upper bed is supported by palynological evidence [185,186]. The upper boundary of the Carnian, ~228.5 Ma [8], thus provides a minimum age for the node.

***23.4 Soft maximum age and justification.*** As in 14.4.

**24 Crown-Mantodea** (112.6 – 241.5 Ma), node 24

***24.1 Fossil taxon and specimen.*** *Cretophotina santanensis* Lee, 2014 (Chaeteessidae) [(SMNS 67583 (holotype): State Museum of Natural History Stuttgart, Stuttgart, Germany]. Nova Olinda quarry, Nova Olinda Member of the Lower Cretaceous (Aptian) Crato Formation, northeastern Brazil [187].

***24.2 Phylogenetic justification.*** *Cretophotina santanensis* is known from a body with wings preserved as a compression. It is identified a mantodean by wing characters [187]. It likely represents a stem group to the extant family Chaeteessidae [71], which may be the basalmost extant mantis family [188]. We conservatively use *C. santanensis* to calibrate the node representing crown Mantodea.

***24.3 Minimum age and justification.*** As in 3.3.

***24.4 Soft maximum age and justification.*** As in 9.4.

***24.5 Discussion.*** The fossil record of crown-group mantises is sparse. An oft-cited Mesozoic representative is *Ambermantis* [189], whose systematic position was however questioned [190]. *Protohierodula crabbi* from the Priabonian Insect Limestone of the Isle of Wight, UK is another possible crown-mantis [191], although wing-venation characters are not unequivocal and the fossil has been more recently treated as *incertae sedis* [192,193]. Further uncontested crown-mantises are from the Palaeocene of Menat.

**25 Crown-Blaberidae** (59.2 – 130.8 Ma), node 25

***25.1 Fossil taxon and specimen.*** *"Gyna" obesa* Piton, 1940 (Blaberidae) [MNHN.F.R06689 (holotype): Muséum national d'Histoire naturelle (National Museum of Natural History), Paris, France]. South-east of the village of Menat, Menat Basin, Puyde-Dôme, France [194–196].

***25.2 Phylogenetic justification.*** Piton [194] placed this single fossil known from the Paleocene of Meant into the extant blaberid genus *Gyna*. Evangelista et al. [196] redescribed the fossil and concluded that the generic and subfamilial position of the fossil is not substantiated. However the apomorphic shape of the subgenital plate still supported attribution to Blaberidae [195,196]. We therefore use the fossil to calibrate the node representing crown Blaberidae.

***25.3 Minimum age and justification.*** Mammalian biostratigraphy, namely the presence of the genus *Plesiadapis* suggests a Selandian age [197], whose upper boundary at 59.2 Ma [8], provides the minimum age for the node.

***25.4 Soft maximum age and justification.*** As in 21.4.

**26 Crown-Blattodea** (126.3 – 241.5 Ma), node 26

***26.1 Fossil taxon and specimen.*** *Cretaholocompsa montsecana* Martinez- Delclòs, 1993 (Corydiidae) [LC-1704-IEI (holotype): Fundació Pública Institut d'Estudis llerdencs en Lleida, Lleida, Spain]. La Cabrua outcrop, Sierra del Montsec, Barremian La Pedrera de Rubies Formation, Montsec, Spain [195,198].

***26.2 Phylogenetic justification.*** The species is known from a compression fossil preserving part of the head, thorax, mesothoracic legs, and tegmina. It can be assigned to Holocompsinae (Corydiidae) based on the presence of the following synapomorphic characters: apical portion of tegmina without venation; radius (R) shortened, with anterior branches; CuA simple and incomplete; and CuP basally or medially with an angulate plical furrow (Rehn 1951; Evangelista et al. 2017). The presence of numerus anal veins, that are not found in recent Holocompsinae [196,198], indicate that *Cretaholocompsa* represents a stem group to the subfamily, thus assigning it to crown-Blattodea. An analysis of 2,370 nuclear protein-coding genes, that recovered a monophyletic Blattodea, found Corydiidae to form a sister group to Tryonicidae + Blattidae + Lamproblattidae + Cryptocercidae + Isoptera [195].

***26.3 Minimum age and justification.*** The La Pedrera de Rubies Formation has been dated to the Barremian based on the presence of the charophyte *Atopochara trivolvis triquetra* [199,200]. Although earlier studies have indicated an earlier age of the fossiliferous strata [200], we use a more conservative age of 126.3 Ma based on the upper boundary of the Barremian [8].

***26.4 Soft maximum age and justification.*** As in 9.4.

**27 Stem Isoptera** (130.8 – 167.5 Ma), node 27

***27.1 Fossil taxon and specimen.*** *Valditermes brenanae* Jarzembowski, 1981 (Mastotermitidae) [In. 64588 (holotype); In. 64589-93 (paratypes): Natural History Museum, London, UK]. Clockhouse Brickworks locality, Surrey, Hauterivian–Barremian Weald Formation, southern England [201].

***27.2 Phylogenetic justification.*** *Valditermes*, known from isolated wings, possesses the humeral suture on the forewings, a character unique to Isoptera [202]. The placement of *Valditermes* is further supported by two phylogenetic analyses [203,204]. Both studies have recovered the genus as a stem group to the family Mastotermitidae. Mastotermitidae are sister to all remaining extant termites [204,205]. The monophyly of Isoptera is supported by molecular analyses [124,206]. However, Evangelista et al. [195] noted ambiguities in the character scoring for the taxon in previous phylogenetic studies, and thus conservatively used it to calibrate stem-Isoptera. We follow the same placement here.

***27.3 Minimum age and justification.*** The presence of the ostracod *Cypridea tuberculate* in sediments of the Lower Weald Clay place it into the *Theriosynoecum fittoni* Zone [207]. Palynomorphs from the Lower Weald Clay indicate that it lies at the boundary of the Hauterivian and Barremian [208]. The upper boundary of the Hauterivian, 130.8 Ma [8], thus provides the minimum age of the horizon.

***27.4 Soft maximum age and justification.*** As in 10.4.

**28 Crown-Mastotermitidae** (98.17 – 167.5 Ma), node 28

***28.1 Fossil taxon and specimen.*** *Mastotermes monostichus* Zhao, Eggleton, and Ren, 2019 (Mastotermitidae) [CNU-TER-BU-2017004 (holotype): Key Lab of Insect Evolution and Environmental Changes, College of Life Sciences, Capital Normal University, Beijing, China]. Burmese amber, Hukawng Valley, Myitkyina District, Kachin State, northern Myanmar [209].

***28.2 Phylogenetic justification.*** *Mastotermes monostichus*, known an adult entombed in amber, can be placed into the family Mastotermitidae on the basis of the sclerotised M vein as well as Sc, R, and

Rs. Assignment to the extant genus *Mastotermes* is furthermore supported by a formal cladistic analysis [209]. *Mastotermes monostichus* is therefore suitable for calibrating the node representing crown-Mastotermitidae.

***28.3 Minimum age and justification.*** As in 19.3.

***28.4 Soft maximum age and justification.*** As in 10.4.

**29 Stem-Neoisoptera** (98.17 – 167.5 Ma), node 29

***29.1 Fossil taxon and specimen.*** *Archeorhinotermes rossi* Krishna and Grimaldi, 2003 (Mastotermitidae) [BMNH In. 20160 (holotype): Natural History Museum, London, UK]. Burmese amber, Hukawng Valley, Myitkyina District, Kachin State, northern Myanmar [209].

***29.2 Phylogenetic justification.*** *Archeorhinotermes rossi*, known from a single amber inclusion, has been initially placed into the neoisopteran family Rhinotermitidae by Krishna and Grimaldi [210]. In a subsequent morphological phylogenetic analysis, *Archeorhinotermes* was recovered as the stem group to the rest of the Neoisoptera [203]. This placement is supported by the frontal gland developed into distinct fontanelle, and costalized forewings [195]. We thus used *A. rossi* to calibrate the split between Neoisoptera and the remaining termites.

***29.3 Minimum age and justification.*** As in 19.3.

***29.4 Soft maximum age and justification.*** As in 10.4.

**30 Stem-Hemiptera** (319.9 – 428.9 Ma), node 30

***30.1 Fossil taxon and specimen.*** *Protoprosbole straeleni* Laurentiaux, 1952 (Protoprosbolidae) [IRSNB a9885 (holotype): Institut Royal des Sciences Naturelles de Belgique, Brussels, Belgium]. Monceau-Fontaine colliery, shaft 10, Charbonnage de Monceau-Fontaine, Charleroi Coal Basin, Belgium [211,212].

***30.2 Phylogenetic justification.*** The systematic position *P. straeleni*, known from a forewing, has been debated for over sixty years since its description [212]. It shares with Hemiptera the cua-cup touching CuP; and the flexion or nodal line following the course of RA. It however differs from extant hemipterans by the presence of three veins in the anal area [212]. We thus use *Protoprosbole* to calibrate the stem of Hemiptera. Monophyly of the order is supported by phylogenomic analyses and morphology [5,169,182,213].

***30.3 Minimum age and justification.*** The deposit has been assigned to the *Bilinguites superbilinguis* R2c2 subzone, corresponding to the late Namurian B [214]. While the boundary of the late Namurian and early Westphalian lacks a precise isotopic date, Pointon et al. [215] estimated the base of the Westphalian at *ca.* 319.9 Ma.

***30.4 Soft maximum age and justification.*** As in 1.4.

**31 Stem Euhemiptera** (306.7 – 428.9 Ma), node 31

***31.1 Fossil taxon and specimen.*** *Aviorrhyncha magnifica* Nel, Bourgoin, Engel, and Szwedo, 2013 (†Aviorrhynchidae) [Avion n° 2 (holotype): Muséum national d'Histoire naturelle (National Museum of Natural History), Paris, France]. Avion outcrop (Moscovian), Department of Pas-de-Calais, France [216].

***31.2 Phylogenetic justification.*** *Aviorrhyncha magnifica* known from a forewing, shares with Euhemiptera (i.e., all living Hemiptera except Sternorrhyncha) the presence of an ambient vein (synapomorphy); a well-developed, concave CP7 (synapomorphy); a concave ScP fused with R very close to wing base and re-emerging distally as a veinlet between R and anterior wing margin (proposed character of Euhemiptera); a convex vein AA1+2 at least partly fused with the concave CuP5. It has therefore been used in the past to calibrate the node representing stem-Euhemiptera [76]. The monophyly of Euhemiptera is supported by phylogenomic datasets [213]. We thus use *Aviorrhyncha* to calibrate the split between Sternorrhyncha and the remaining Hemiptera.

***31.3 Minimum age and justification.*** The fossil was found in the "Terril No. 7" layer, which consists of rocks from the slag heaps of coal mines 3 and 4 of Liévin in the Avion outcrop [216]. These coal mines were correlated to the Westphalian C/D, or Bolsovian/Asturian [216]. The upper boundary of the Moscovian, 306.7 Ma [8], provides the minimum age.

***31.4 Soft maximum age and justification.*** As in 1.4.

***31.5 Discussion.*** An alternative candidate for calibrating the node would be the yet undescribed representative of crown-Sternorrhyncha from the same Avion outcrop [217].

**32 Crown Holometabola** (306.7 – 428.9 Ma), node 32

***32.1 Fossil taxon and specimen.*** *Westphalopsocus pumilio* Azar, Nel, Engel, and Bourgoin, 2013 (†Westphalopsocidae) [MGPV Avion 4 (holotype): Patrick Roques collection, Entomological Laboratory, Muséum national d'Histoire naturelle, Paris, France]. Avion outcrop (Moscovian, Department of Pas-de-Calais, France [216].

***32.2 Phylogenetic justification.*** The specimen, preserved as a single forewing, can be placed into Paraneoptera based on the presence of a cua-cup vein between CuP and M+CuA, which represents an autapomorphy of the group [216]. Assignment to Psocodea is supported by the combined presence of an aerola postica; crossveins of RA in the distal portion of the costal area; anal veins fusing apically and forming an inverted 'Y-vein'; ScP completely separated from R until their distal fusion near their apices (autapomorphy of Psocodea); and the elongate shape of the forewing with the anterior and posterior margins parallel [216]. Despite these characters, Aristov [218] moved the taxon to the family Spanioderidae belonging to the extinct order Cnemidolestida, but this was shown to be due to a misinterpretation of wing-venation characters [219]. Notwithstanding the absence of the rest of the body, we treat the fossil as a stem-psocodean, since all Palaeozoic fossils belonging to this group bear features that are never present in extant bark lice, book lice, and true lice. These include pentamerous tarsi, short cerci, and a rather generalized wing venation pattern [146]. The monophyly of Psocodea (Psocoptera + Phthiraptera) is well supported by morphological [220] and molecular evidence [5,221].

***32.3 Minimum age and justification.*** As in 31.3.

***32.4 Soft maximum age and justification.*** As in 1.4.

**33 Crown-Hymenoptera** (228.5 – 314.6 Ma), node 33

***33.1 Fossil taxon and specimen.*** *Triassoxyela foveolata* Rasnitsyn, 1964 (Xyelidae) [PIN 2070/1 (holotype): Paleontological Institute of the Academy of Science of Russia, Moscow, Russia]. Dzhayloucho, Upper Triassic, Ladinian-Carnian Madygen Formation, south of Fergana Valley, Kyrgyzstan [222].

***33.2 Phylogenetic justification.*** *Triassoxyela foveolata,* known from an exoskeleton with wings, can be assigned to the extant family Xyelidae on the basis of wing venation characters discussed by Rasnitsyn [222]. The placement of the fossil as a stem-group member of Xyelidae was confirmed by a total-evidence phylogenetic analysis [223], thus making it suitable for calibrating crown-Hymenoptera.

***33.3 Minimum age and justification.*** As in 5.3.

***33.4 Soft maximum age and justification.*** As in 5.4.

**34 Crown-Apocrita** (163.1 – 241.5 Ma), node 34

***34.1 Fossil taxon and specimen.*** *Cleistogaster buriatica* Rasnitsyn, 1975 (†Cleistogastridae) [PIN 3000/1003 (holotype): Paleontological Institute of the Academy of Science of Russia, Moscow, Russia]. Novospasskoye village site, Ichetuy Formation, Buryat, Russia [224].

***34.2 Phylogenetic justification.*** *Cleistogaster buriatica,* known from an exoskeleton with wings, was initially placed into the extinct subfamily Cleistogastrinae within the Megalyridae by Rasnitsyn [224]. Shaw [225] and later Perrichot [226] considered the taxon as a separate family, Cleistogastridae. While wing-venation characters are similar to modern Trigonalyidae, *Cleistogaster* differs in the presence of a very long ovipositor [227]. In a phylogenetic analysis, *Cleistogaster* was recovered as sister to Ichneumonidae, thus placing it into crown-Apocrita [223,228].

***34.3 Minimum age and justification.*** The Novospasskoye locality is usually considered Upper to Middle Jurassic based on biostratigraphy [229]. We thus use 163.1 Ma as a conservative constraint on the node, representing the upper boundary of the Callovian [8].

***34.4 Soft maximum age and justification.*** As in 9.4.

**35 Stem-Coleoptera** (290.1 Ma – 314.6 Ma), node 35

***35.1 Fossil taxon and specimen.*** *Coleopsis archaica* Kirejtshuk, Poschmann, and Nel, 2014 (Coleopsidae) [ZfB 3315 (holotype): Zentrum fur Biodokumentation des Saarlandes, Schiffweiler, Germany]. Rotliegend, Meisenheim Formation, Saarland, Germany [230,231].

***35.2 Phylogenetic justification.*** *Coleopsis archaica*, preserved as a compression of a body in ventral aspect, can be assigned to Coleoptera based on the presence of sclerotised forewings (elytra) with 'window' punctures arranged in rows which are typical of early beetles from the Permian [232] and the genitalia apparently retracted into the abdomen. Based on the elytral punctation and the presence of putative 'veins' (although the true identity of these structures remains rather ambiguous),

*Coleopsis* can be placed into Protocoleoptera, a group that has been recovered as a stem group to Coleoptera in a morphological phylogenetic analysis [233]. The monophyly of Coleoptera has not been seriously challenged, either on morphological or on molecular grounds [5,234,235].

***35.3 Minimum age and justification.*** Poschmann and Schindler [236] reported the biostratigraphically important cockroach *Spiloblattina odernheimensis* from the locality, which indicates a Sakmarian age [237,238]. However, amphibian remains have been interpreted as evidence for an Asselian date [239]. We conservatively follow the younger Sakmarian date. The minimum boundary of the Sakmarian has been defined at 290.1 Ma [8].

***35.4 Soft maximum age and justification.*** As in 5.4.

***35.5 Discussion.*** *Adiphlebia lacoana*, regarded by Béthoux [240] as the oldest stem-beetle, has been recommended as a suitable calibration fossil [71]. However, re-examination of additional material has revealed that the species lacks coleopteran apomorphies and that the “box-like veins” on its forewings are in fact merely lumps of clay and not remnants of a biological structure [216,241,242]. Likewise, the systematic position of the purportedly oldest coleopterid (Strepsiptera + Coleoptera), *Stephanastus polinae* from the Gzhelian of France [216], is unclear. Beutel et al. [243] called into question the original interpretation of the wing-venation characters of the holotype, pointed out uncertainties regarding its wing-folding pattern, and concluded that it cannot be excluded from the enigmatic extinct order Protelytroptera. This makes *Coleopsis archaica* the oldest reliable stem-beetle.

**36 Stem-Neuroptera** (272.3 Ma – 314.6 Ma), node 36

***36.1 Fossil taxon and specimen.*** *Elmothone martynovae* Carpenter, 1976 (†Permithonidae) [5585 (holotype): Museum of Comparative Zoology, Harvard University, Cambridge, USA]. Elmo, Carlton Limestone Member of the Wellington Formation, Dickinson County, central Kansas, USA [244].

***36.2 Phylogenetic justification.*** The specimen, represented by a forewing, has been recovered as a stem group to the extant Neuroptera in a morphological phylogenetic analysis [245]. However, the monophyly of Permithonidae has not yet been subjected to a formal cladistic investigation [246].

***36.3 Minimum age and justification.*** The Elmo locality has been biostratigraphy correlated with the Leonardian regional stage based on conchostracan fossils, which corresponds to the Artinskian to Kungurian [247,248]. Thus, the minimum age of *E. martynovae* is provided by the minimum boundary of the Kungurian, 272.3 Ma [8].

***36.4 Soft maximum age and justification.*** As in 6.4.

***36.5 Discussion.*** A putative stem-neuropteran from the Sakmarian–Artinskian locality Obora in Moravia, Czech Republic was mentioned by Kukalová [249] but remains undescribed. Some authors consider the permoberothid *Permoberotha villosa* [250] from the Permian Elmo Lagerstätten in Kansas as the oldest stem-neuropteran [182,251,252]. The species has been assigned to the extinct Permian-Jurassic order Glosselytrodea by Carpenter [146], but the systematic position of this enigmatic group within Eumetabola is unclear [253].

**37 Crown-Raphidioptera** (209.6 Ma – 314.6 Ma), node 37

***37.1 Fossil taxon and specimen.*** Raphidioptera sp. (family indet.) [ID unassigned (figured specimen):

Paleontological Institute of the Academy of Science of Russia, Moscow, Russia]. Schönbachsmühle quarry near Ebelsbach, Late Carnian Hassberge Formation, Bavaria, Germany [244].

***37.2 Phylogenetic justification.*** This undescribed fossil can be assigned to the order Raphidioptera based on its elongate and flat head, and distinctly elongate prothorax (both apomorphic [3]) [254]. We thus use the fossil to calibrate the split between Raphidioptera and Megaloptera.

***37.3 Minimum age and justification.*** The Hassberge Formation (previously known as Coburger Sandstein and Blasensandstein) has been regarded as latest Carnian to early Norian [255]. We thus use the upper boundary of the Norian, ~209.6 Ma [8], as a conservative minimum bound on the node.

***37.4 Soft maximum age and justification.*** As in 5.4.

**38 Crown-Panorpida** (313.7 – 428.9 Ma), node 38

***38.1 Fossil taxon and specimen.*** *Westphalomerope maryvonneae* Nel, Roques, Nel, Prokop, and Steyer, 2007 [MNHN-LP-R.55181 (holotype): Patrick Roques collection, Entomological Laboratory, Muséum national d'Histoire naturelle (National Museum of Natural History), Paris, France]. Pas-de-Calais, Bruay-la-Bussière, Faisceau de Modeste, Veine Maroc, France [256].

***38.2 Phylogenetic justification.*** The single hindwing described as *W. maryvonneae* can be attributed to the extinct family Protomeropidae based on the CuA directly connected with M and negative, but with a convex, heavily sclerotized fold running in front of it and disappearing in the wing membrane [257]. It possesses a putative protomeropid apomorphy “ScP distally fused with RA” [258]. The family Protomeropidae has not been included in any formal phylogenetic analysis [71] but can be placed within total-group Mecoptera since the males possess Carpenter’s organs, which have been considered as a probable apomorphy of crown-Mecoptera [259]. This places *Westphalomerope* within crown-Panorpida and makes it suitable for calibrating the split between panorpids and Amphiesmenoptera (Trichoptera + Lepidoptera).

***38.3 Minimum age and justification.*** The specimen originates from the “Terril no. 5” horizon of the “Faisceau de Modeste” locality in Bruayla-Bussière [256]. The locality has been correlated with the Westphalian A stage [256]. The upper boundary of the Westphalian B constrained by U-Pb zircon dating is 313.78 Ma ± 0.08 Ma [215], providing a minimum age of 313.7 Ma.

***38.4 Soft maximum age and justification.*** As in 1.4.

**39 Crown-Trichoptera** (313.7 – 428.9 Ma), node 39

***39.1 Fossil taxon and specimen.*** *Folindusia* (ichnotaxon) [NIGP162044: Nanjing Institute of Geology and Palaeontology, Chinese Academy of Sciences, Nanjing, China]. Huayuangou outcrop of the Carnian Karamay Formation, Karamay City, Xinjiang, northwestern China [260].

***39.2 Phylogenetic justification.*** The cases can be recognised as belonging to Trichoptera based on their characteristic cylindrical shape. They appear to be constructed from small pebbles, possibly reinforced with silk [261].

***39.3 Minimum age and justification.*** The age of the Karamay entomofauna is considered to be Carnian based on megaspore fossils [262,263]. A minimum age of 228.5 Ma is given by the upper boundary of the Carnian [8].

***39.4 Soft maximum age and justification.*** As in 1.4.

***39.5 Discussion.*** *Necrotaulius proximus* and *Prorhyacophila furcata* (Necrotauliidae) described from the Ladinian-Carnian Madygen Formation, south of Fergana Valley, Kyrgyzstan [261,264] appear to represent the earliest body fossil record of stem-Trichoptera. However, the fossil record of caddisfly larval cases is longer than that of their builders, almost certainly due to the higher preservation potential of the former. Mouro et al. [265] described putative caddisfly cases from Asselian–Sakmarian marine deposits of Brazil. However, it is not possible to exclude other potential builders of these structures [71]. In particular, the marine palaeoenvironment strongly suggests that they most likely belong to polychaetes. The earliest undisputed trichopteran larval case is thus *Folindusia* along with *Terrindusia* from Karamay, followed by mid-Jurassic cases from Transbaikalian Russia [266].

**40 Stem-Mecoptera** (268.8 – 314.6 Ma), node 40

***40.1 Fossil taxon and specimen.*** *Pseudonannochorista willmanni* Novokshonov, 1994 (†Permochoristidae) [PIN966/21: Paleontological Institute of the Academy of Science of Russia, Moscow, Russia]. Kaltan, Kemerovo, Permian Mitina Formation, Russia [267,268].

***40.2 Phylogenetic justification.*** *Pseudonannochorista willmanni*, known from wings, belongs to Pseudonannochoristinae, a subfamily of Permochoristidae [268,269]. While this family has been determined as paraphyletic in a phylogenetic analysis, *Pseudonannochorista* can still be used to calibrate the node representing stem-Mecoptera [245].

***40.3 Minimum age and justification.*** The age of the insect-bearing Mitino Horizon at the Kaltan locality in the Kuznetsk Basin is Roadian, based on the ammonoid fauna [71,270]. The upper boundary of the Roadian, 268.8 Ma [8], can thus be used to constrain the node.

***40.4 Soft maximum age and justification.*** As in 5.4.

**41 Stem-Siphonaptera** (162.5 – 314.6 Ma), node 41

***41.1 Fossil taxon and specimen.*** *Hadropsylla sinica* Huang, Engel, Cai, and Nel, 2013 (†Pseudopulicidae) [NIGP 154257 (holotype); NIGP 154258 (paratype): Nanjing Institute of Geology and Palaeontology, Nanjing, China]. Locality near the Daohugou Village, Wuhua Township, Ningcheng County, Chifeng City, Middle Jurassic Haifanggou Formation, Inner Mongolia, northeast China [271].

***41.2 Phylogenetic justification.*** *Hadropsylla sinica,* known from compression fossils from Daohugou, are placed into Siphonaptera by their specialised tarsal attachment structures, lack of wings, and siphonate mouthparts [271–273]. While *H. sinica* and other Jurassic fleas lack some apomorphies of modern fleas [274], the abovementioned characters unambiguously assign them to the siphonapteran stem group [273].

***41.3 Minimum age and justification.*** As in 6.3.

***41.4 Soft maximum age and justification.*** As in 5.4.

**42 Crown-Diptera** (240.5 – 314.6 Ma), node 42

***42.1 Fossil taxon and specimen.*** *Grauvogelia arzvilleriana* Krzemiński, Krzemińska, and Papier, 1994 (†Grauvogeliidae) [5514 (holotype): Grauvogel & Gall collection, University of Strasbourg, Strasbourg, France]. Arzviller, Grès à Voltzia Formation, France [275].

***42.2 Phylogenetic justification.*** *Grauvogelia arzvilleriana*, known from an isolated forewing, is the oldest definitive fossil that can be assigned to Diptera [275,276]. The placement was confirmed by a morphological phylogenetic analysis which recovered *Grauvogelia* as a stem group of Psychodomorpha [277]. The stem of Psychodomorpha has been recovered within crown-Diptera by a molecular analysis [278].

***42.3 Minimum age and justification.*** As in 7.3.

***42.4 Soft maximum age and justification.*** As in 5.4.

### Supplementary Figures

**Figure S1.** Distribution of calibration points on the phylogeny of Polyneoptera.

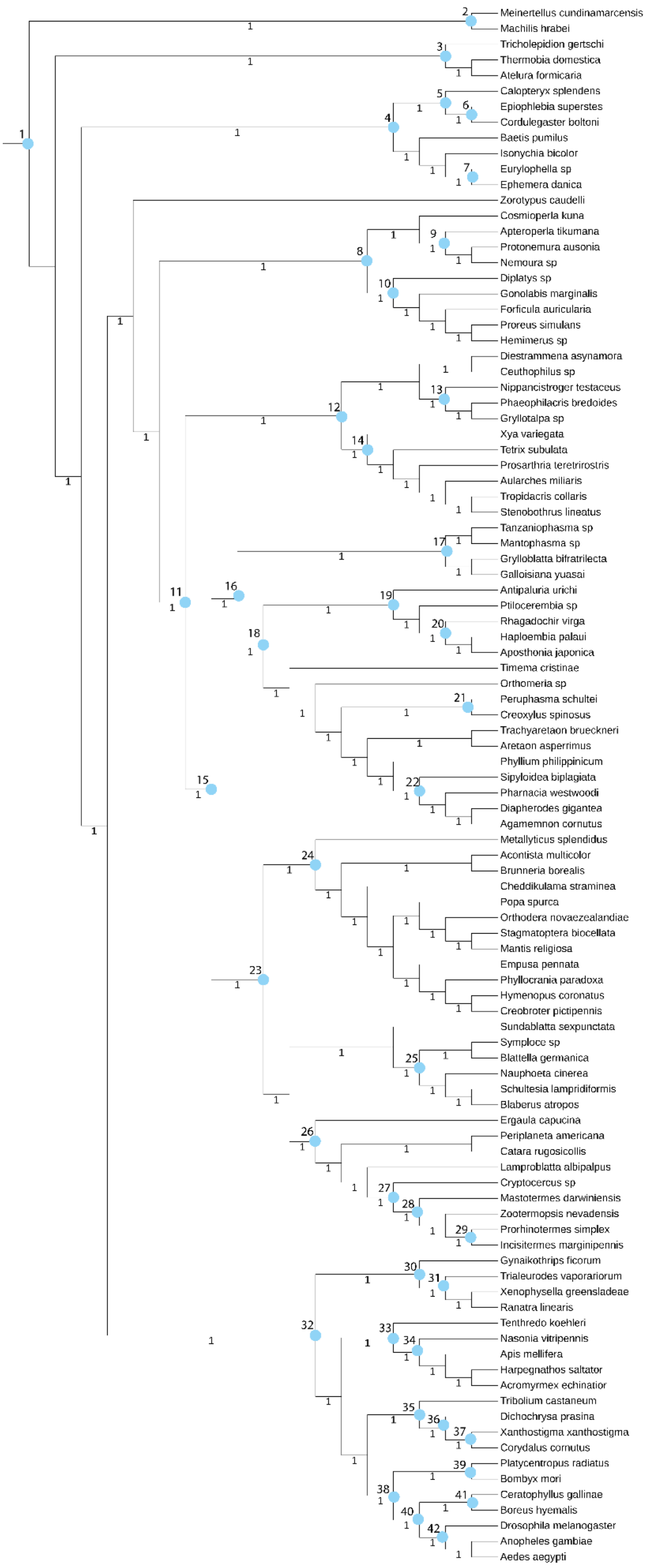

**Figure S2.** Habitus view of *Zorotypus nascimbenei* comb. nov. in mid-Cretaceous amber from northern Myanmar (~99 Ma; NIGP175112) under green fluorescence in (**a**) dorsal and (**b**) ventral views. Abbreviations: abd, abdomen; ce, cercus; mp, maxillary palpomere, msf, mesofemur; mst, mesothorax; mtt; metathorax; mttf, metafemur; pf, profemur; pt, prothorax. Scale bars: 200 µm.

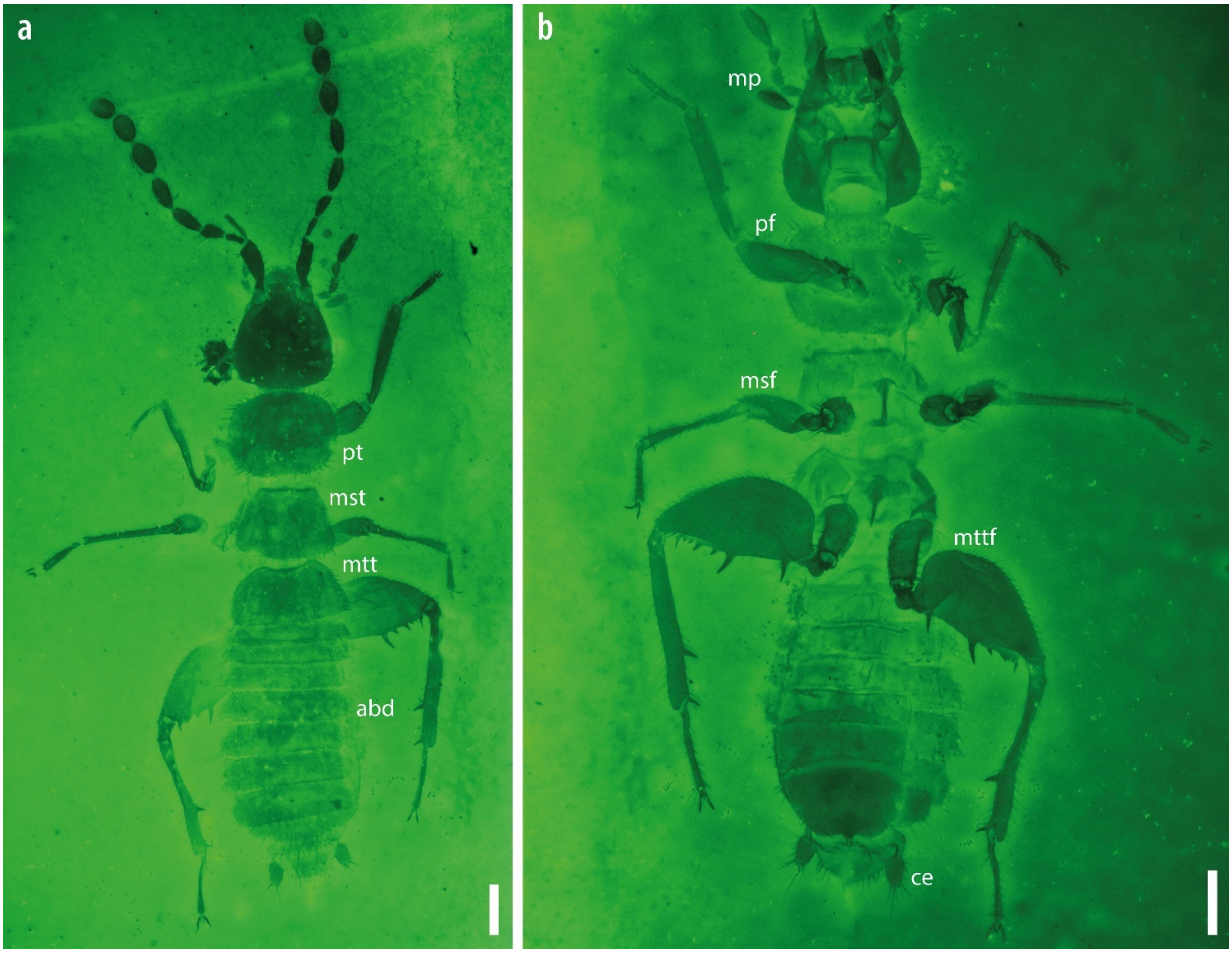

**Figure S3.** Morphological details of *Zorotypus nascimbenei* comb. nov. from mid-Cretaceous amber from northern Myanmar (~99 Ma; NIGP175112). **a** Head in ventral view. **b** Labial palp. **c** Prothoracic leg. **d** Mesothoracic leg. **e** Abdominal apex. **f** Metathoracic leg. Abbreviations: a1, antennomere 1 (scape); g, galea; lp, labial palp; md, mandible; mp, maxillary palp; msta1–2, mesotarsi 1–2; msti, mesotibia; mtta1–2, metatarsi 1–2; mttf, metafemur; mtti, metatibia; pta1–2, protarsi 1–2; pti, protibia. Scale bars: 100 μm (a, d), 50 μm (b, c, e, f).

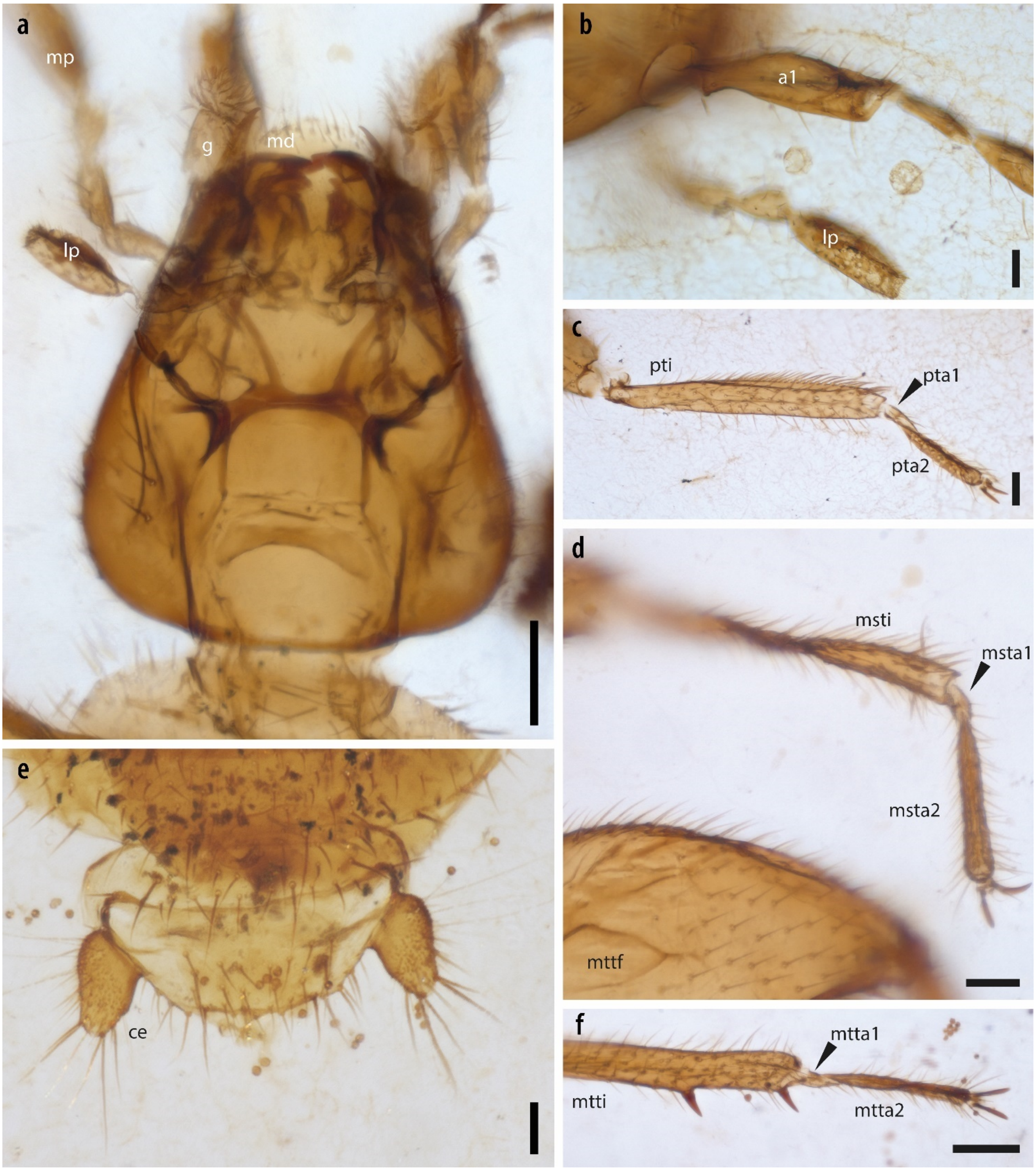

**Figure S4.** Phylogeny of Polyneoptera inferred with the site-heterogeneous model CAT-GTR+G implemented in PhyloBayes, on the 106-taxon dataset.

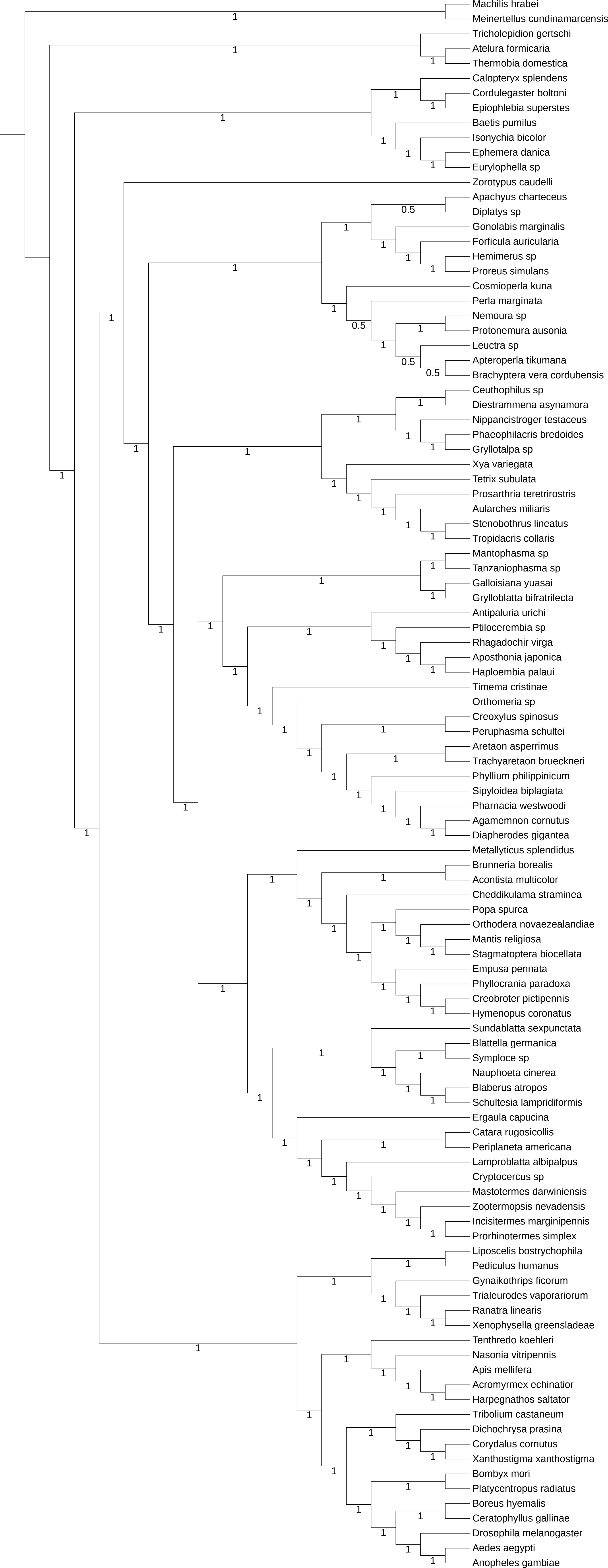
Machilis hrabei
Meinertellus cundinamarcensis
Tricholepidion gertschi
Atelura formicaria
Thermobia domestica
Calopteryx splendens
Cordulegaster boltoni
Epiophlebia superstes
Baetis pumilus
Isonychia bicolor
Ephemera danica
Eurylophella sp
Zorotypus caudelli
Apachyus charteceus
Diplatys sp
Gonolabis marginalis
Forficula auricularia
Hemimerus sp
Proreus simulans
Cosmioperla kuna
Perla marginata
Nemoura sp
Protonemura ausonia
Leuctra sp
Apteroperla tikumana
Brachyptera vera cordubensis
Ceuthophilus sp
Diestrammena asynamora
Nippancistroger testaceus
Phaeophilacris bredoides
Gryllotalpa sp
Xya variegata
Tetrix subulata
Prosarthria teretrirostris
Aularches miliaris
Stenobothrus lineatus
Tropidacris collaris
Mantophasma sp
Tanzaniophasma sp
Galloisiana yuasai
Grylloblatta bifratrilecta
Antipaluria urichi
Ptilocerembia sp
Rhagadochir virga
Aposthonia japonica
Haploembia palaui
Timema cristinae
Orthomeria sp
Creoxylus spinosus
Peruphasma schultei
Aretaon asperrimus
Trachyaretaon brueckneri
Phyllium philippinicum
Sipyloidea biplagiata
Pharnacia westwoodi
Agamemnon cornutus
Diapherodes gigantea
Metallyticus splendidus
Brunneria borealis
Acontista multicolor
Cheddikulama straminea
Popa spurca
Orthodera novaezealandiae
Mantis religiosa
Stagmatoptera biocellata
Empusa pennata
Phyllocrania paradoxa
Creobroter pictipennis
Hymenopus coronatus
Sundablatta sexpunctata
Blattella germanica
Symploce sp
Nauphoeta cinerea
Blaberus atropos
Schultesia lampridiformis
Ergaula capucina
Catara rugosicollis
Periplaneta americana
Lamproblatta albipalpus
Cryptocercus sp
Mastotermes darwiniensis
Zootermopsis nevadensis
Incisitermes marginipennis
Prorhinotermes simplex
Liposcelis bostrychophila
Pediculus humanus
Gynaikothrips ficorum
Trialeurodes vaporariorum
Ranatra linearis
Xenophysella greensladeae
Tenthredo koehleri
Nasonia vitripennis
Apis mellifera
Acromyrmex echinatior
Harpegnathos saltator
Tribolium castaneum
Dichochrysa prasina
Corydalus cornutus
Xanthostigma xanthostigma
Bombyx mori
Platycentropus radiatus
Boreus hyemalis
Ceratophyllus gallinae
Drosophila melanogaster
Aedes aegypti
Anopheles gambiae

**Figure S5.** Phylogeny of Polyneoptera inferred with the site-heterogeneous model CAT-GTR+G implemented in PhyloBayes, on the 102-taxon dataset.

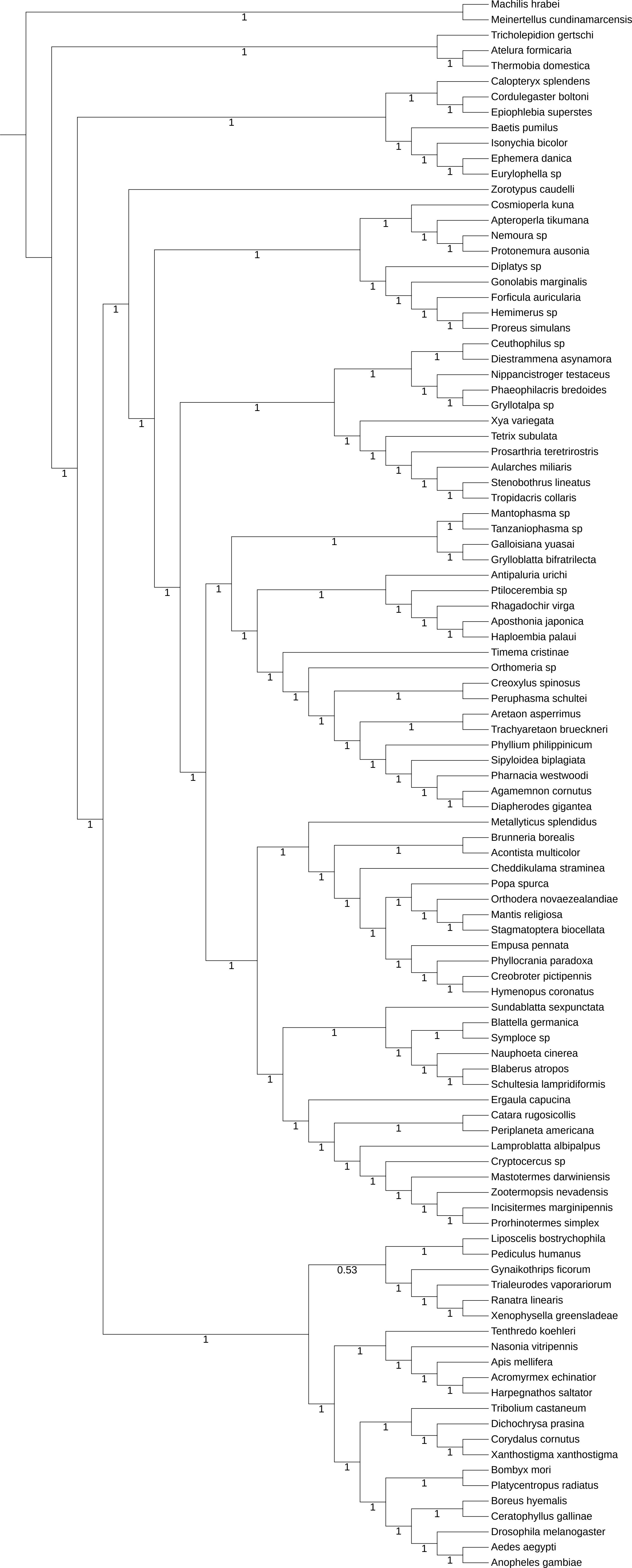
Machilis hrabei
Meinertellus cundinamarcensis
Tricholepidion gertschi
Atelura formicaria
Thermobia domestica
Calopteryx splendens
Cordulegaster boltoni
Epiophlebia superstes
Baetis pumilus
Isonychia bicolor
Ephemera danica
Eurylophella sp
Zorotypus caudelli
Cosmioperla kuna
Apteroperla tikumana
Nemoura sp
Protonemura ausonia
Diplatys sp
Gonolabis marginalis
Forficula auricularia
Hemimerus sp
Proreus simulans
Ceuthophilus sp
Diestrammena asynamora
Nippancistroger testaceus
Phaeophilacris bredoides
Gryllotalpa sp
Xya variegata
Tetrix subulata
Prosarthria teretrirostris
Aularches miliaris
Stenobothrus lineatus
Tropidacris collaris
Mantophasma sp
Tanzaniophasma sp
Galloisiana yuasai
Grylloblatta bifratrilecta
Antipaluria urichi
Ptilocerembia sp
Rhagadochir virga
Aposthonia japonica
Haploembia palaui
Timema cristinae
Orthomeria sp
Creoxylus spinosus
Peruphasma schultei
Aretaon asperrimus
Trachyaretaon brueckneri
Phyllium philippinicum
Sipyloidea biplagiata
Pharnacia westwoodi
Agamemnon cornutus
Diapherodes gigantea
Metallyticus splendidus
Brunneria borealis
Acontista multicolor
Cheddikulama straminea
Popa spurca
Orthodera novaezealandiae
Mantis religiosa
Stagmatoptera biocellata
Empusa pennata
Phyllocrania paradoxa
Creobroter pictipennis
Hymenopus coronatus
Sundablatta sexpunctata
Blattella germanica
Symploce sp
Nauphoeta cinerea
Blaberus atropos
Schultesia lampridiformis
Ergaula capucina
Catara rugosicollis
Periplaneta americana
Lamproblatta albipalpus
Cryptocercus sp
Mastotermes darwiniensis
Zootermopsis nevadensis
Incisitermes marginipennis
Prorhinotermes simplex
Liposcelis bostrychophila
Pediculus humanus
Gynaikothrips ficorum
Trialeurodes vaporariorum
Ranatra linearis
Xenophysella greensladeae
Tenthredo koehleri
Nasonia vitripennis
Apis mellifera
Acromyrmex echinatior
Harpegnathos saltator
Tribolium castaneum
Dichochrysa prasina
Corydalus cornutus
Xanthostigma xanthostigma
Bombyx mori
Platycentropus radiatus
Boreus hyemalis
Ceratophyllus gallinae
Drosophila melanogaster
Aedes aegypti
Anopheles gambiae
0.53

**Figure S6.** Phylogeny of Polyneoptera inferred with the site-heterogeneous model CAT-GTR+G implemented in PhyloBayes, on the 100-taxon dataset.

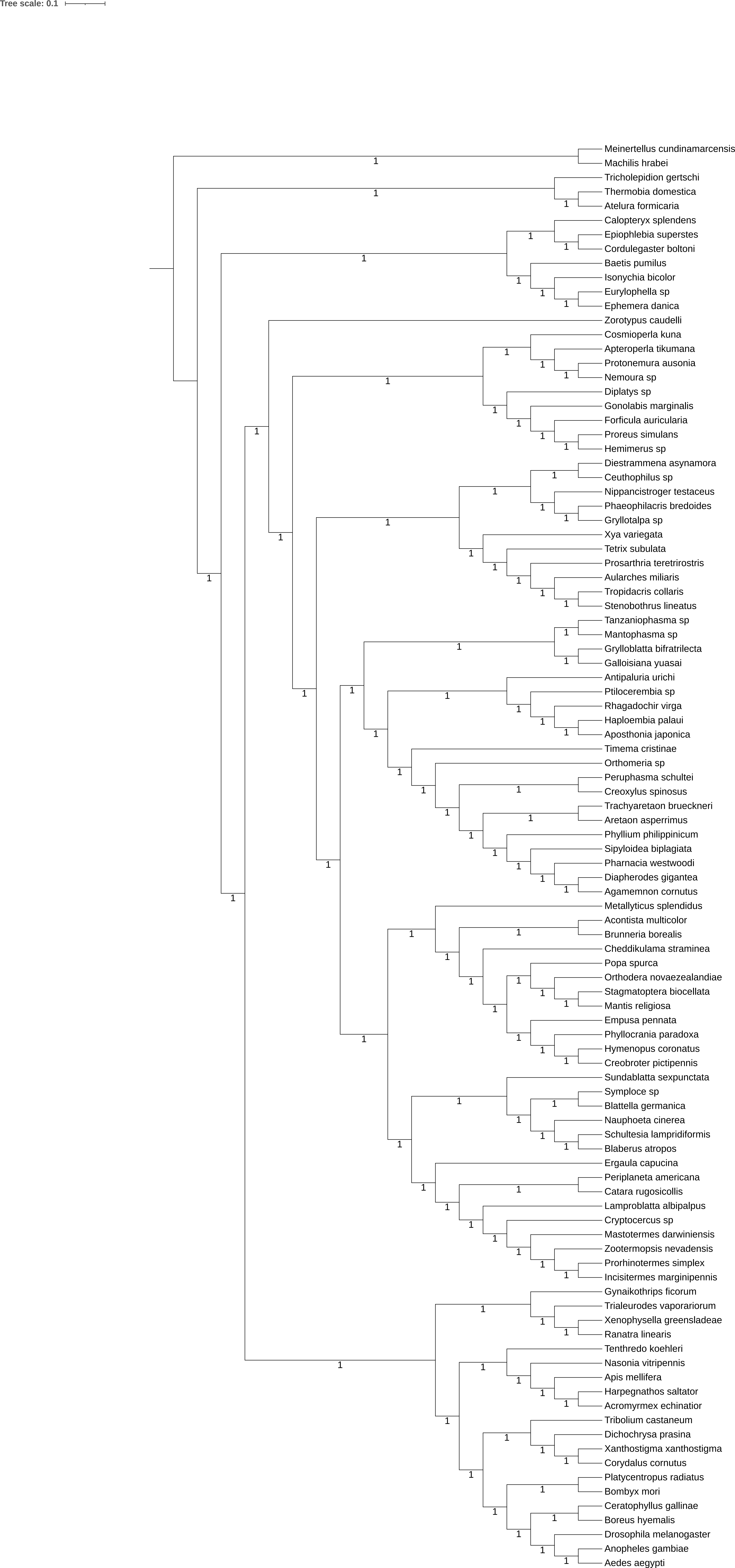
Tree scale: 0.1
Meinertellus cundinamarcensis
Machilis hrabei
Tricholepidion gertschi
Thermobia domestica
Atelura formicaria
Calopteryx splendens
Epiophlebia superstes
Cordulegaster boltoni
Baetis pumilus
Isonychia bicolor
Eurylophella sp
Ephemera danica
Zorotypus caudelli
Cosmioperla kuna
Apteroperla tikumana
Protonemura ausonia
Nemoura sp
Diplatys sp
Gonolabis marginalis
Forficula auricularia
Proreus simulans
Hemimerus sp
Diestrammena asynamora
Ceuthophilus sp
Nippancistroger testaceus
Phaeophilacris bredoides
Gryllotalpa sp
Xya variegata
Tetrix subulata
Prosarthria teretrirostris
Aularches miliaris
Tropidacris collaris
Stenobothrus lineatus
Tanzaniophasma sp
Mantophasma sp
Grylloblatta bifratrilecta
Galloisiana yuasai
Antipaluria urichi
Ptilocerembia sp
Rhagadochir virga
Haploembia palaui
Aposthonia japonica
Timema cristinae
Orthomeria sp
Peruphasma schultei
Creoxylus spinosus
Trachyaretaon brueckneri
Aretaon asperrimus
Phyllium philippinicum
Sipyloidea biplagiata
Pharnacia westwoodi
Diapherodes gigantea
Agamemnon cornutus
Metallyticus splendidus
Acontista multicolor
Brunneria borealis
Cheddikulama straminea
Popa spurca
Orthodera novaezealandiae
Stagmatoptera biocellata
Mantis religiosa
Empusa pennata
Phyllocrania paradoxa
Hymenopus coronatus
Creobroter pictipennis
Sundablatta sexpunctata
Symploce sp
Blattella germanica
Nauphoeta cinerea
Schultesia lampridiformis
Blaberus atropos
Ergaula capucina
Periplaneta americana
Catara rugosicollis
Lamproblatta albipalpus
Cryptocercus sp
Mastotermes darwiniensis
Zootermopsis nevadensis
Prorhinotermes simplex
Incisitermes marginipennis
Gynaikothrips ficorum
Trialeurodes vaporariorum
Xenophysella greensladeae
Ranatra linearis
Tenthredo koehleri
Nasonia vitripennis
Apis mellifera
Harpegnathos saltator
Acromyrmex echinatior
Tribolium castaneum
Dichochrysa prasina
Xanthostigma xanthostigma
Corydalus cornutus
Platycentropus radiatus
Bombyx mori
Ceratophyllus gallinae
Boreus hyemalis
Drosophila melanogaster
Anopheles gambiae
Aedes aegypti

**Figure S7.** Phylogeny of Polyneoptera inferred with the site-heterogeneous model CAT-GTR+G implemented in PhyloBayes, on the 33-taxon dataset.

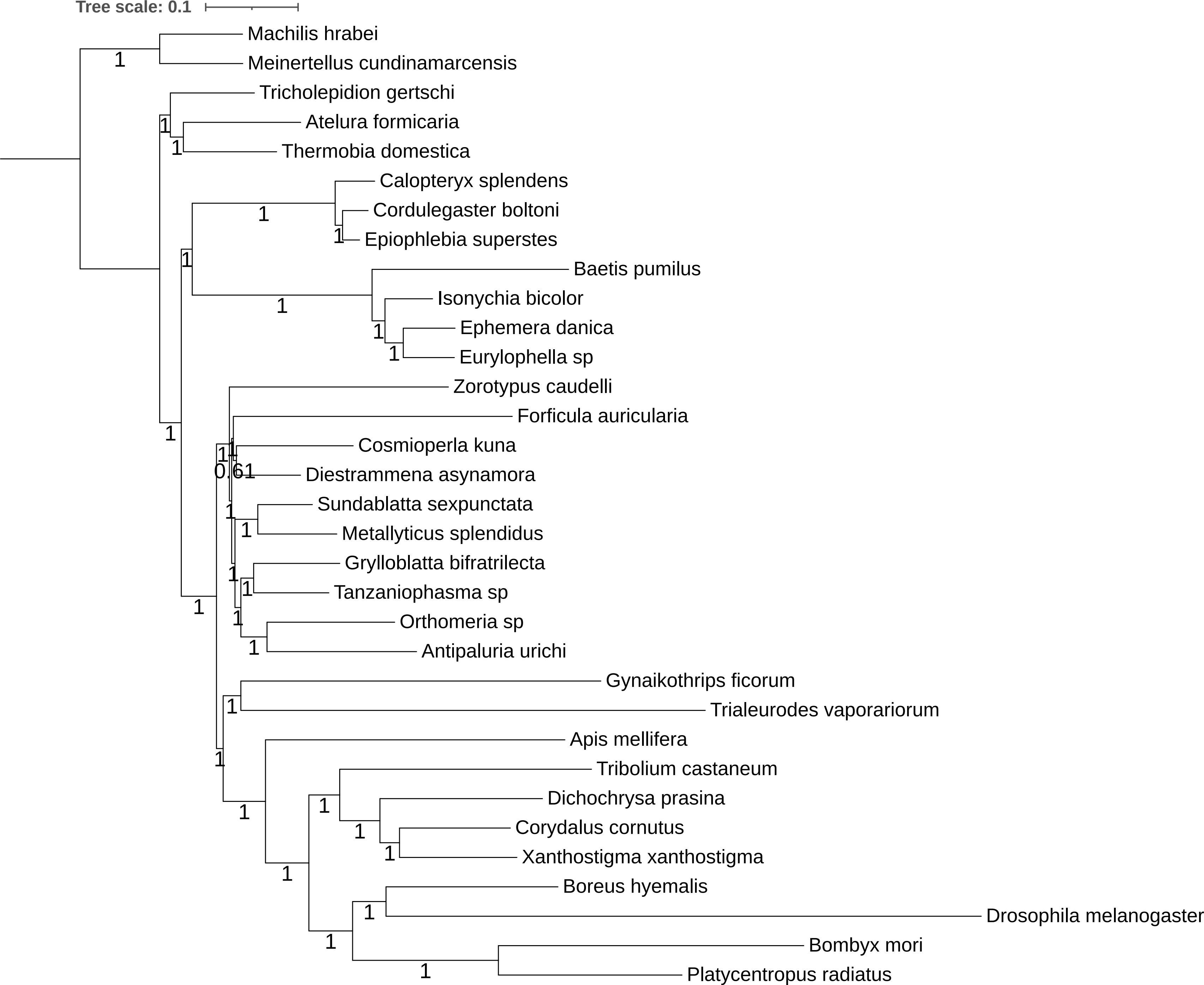
Tree scale: 0.1
Machilis hrabei
Meinertellus cundinamarcensis
Tricholepidion gertschi
Atelura formicaria
Thermobia domestica
Calopteryx splendens
Cordulegaster boltoni
Epiophlebia superstes
Baetis pumilus
Isonychia bicolor
Ephemera danica
Eurylophella sp
Zorotypus caudelli
Forficula auricularia
Cosmioperla kuna
Diestrammena asynamora
Sundablatta sexpunctata
Metallyticus splendidus
Grylloblatta bifratrilecta
Tanzaniophasma sp
Orthomeria sp
Antipaluria urichi
Gynaikothrips ficorum
Trialeurodes vaporariorum
Apis mellifera
Tribolium castaneum
Dichochrysa prasina
Corydalus cornutus
Xanthostigma xanthostigma
Boreus hyemalis
Drosophila melanogaster
Bombyx mori
Platycentropus radiatus
0.61

**Figure S8.** Phylogeny of Polyneoptera inferred with the site-heterogeneous model LG+C60+F+G implemented in IQ-TREE, on the 106-taxon dataset.

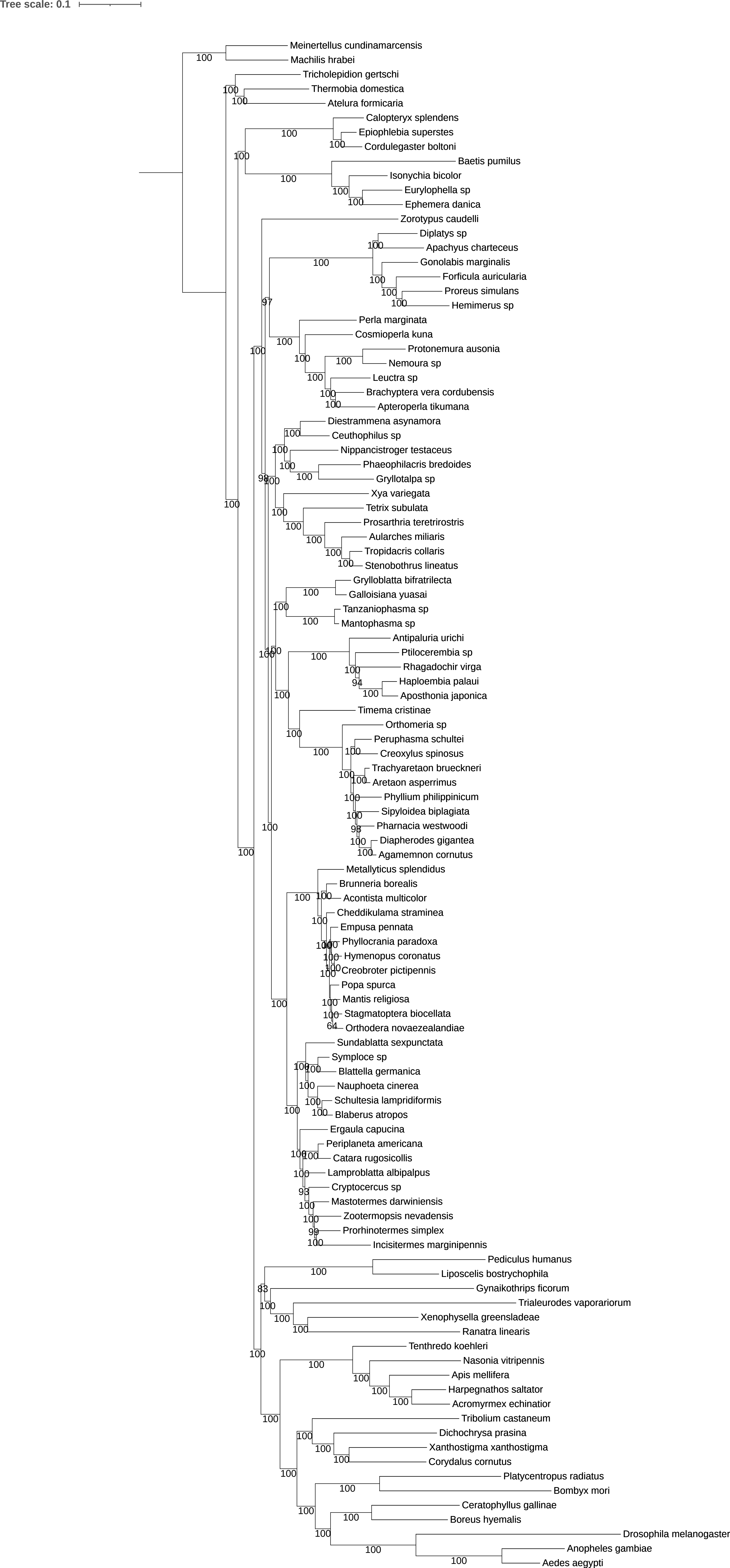
Tree scale: 0.1
Meinertellus cundinamarcensis
Machilis hrabei
Tricholepidion gertschi
Thermobia domestica
Atelura formicaria
Calopteryx splendens
Epiophlebia superstes
Cordulegaster boltoni
Baetis pumilus
Isonychia bicolor
Eurylophella sp
Ephemera danica
Zorotypus caudelli
Diplatys sp
Apachyus charteceus
Gonolabis marginalis
Forficula auricularia
Proreus simulans
Hemimerus sp
Perla marginata
Cosmioperla kuna
Protonemura ausonia
Nemoura sp
Leuctra sp
Brachyptera vera cordubensis
Apteroperla tikumana
Diestrammena asynamora
Ceuthophilus sp
Nippancistroger testaceus
Phaeophilacris bredoides
Gryllotalpa sp
Xya variegata
Tetrix subulata
Prosarthria teretrirostris
Aularches miliaris
Tropidacris collaris
Stenobothrus lineatus
Grylloblatta bifratrilecta
Galloisiana yuasai
Tanzaniophasma sp
Mantophasma sp
Antipaluria urichi
Ptilocerembia sp
Rhagadochir virga
Haploembia palaui
Aposthonia japonica
Timema cristinae
Orthomeria sp
Peruphasma schultei
Creoxylus spinosus
Trachyaretaon brueckneri
Aretaon asperrimus
Phyllium philippinicum
Sipyloidea biplagiata
Pharnacia westwoodi
Diapherodes gigantea
Agamemnon cornutus
Metallyticus splendidus
Brunneria borealis
Acontista multicolor
Cheddikulama straminea
Empusa pennata
Phyllocrania paradoxa
Hymenopus coronatus
Creobroter pictipennis
Popa spurca
Mantis religiosa
Stagmatoptera biocellata
Orthodera novaezealandiae
Sundablatta sexpunctata
Symploce sp
Blattella germanica
Nauphoeta cinerea
Schultesia lampridiformis
Blaberus atropos
Ergaula capucina
Periplaneta americana
Catara rugosicollis
Lamproblatta albipalpus
Cryptocercus sp
Mastotermes darwiniensis
Zootermopsis nevadensis
Prorhinotermes simplex
Incisitermes marginipennis
Pediculus humanus
Liposcelis bostrychophila
Gynaikothrips ficorum
Trialeurodes vaporariorum
Xenophysella greensladeae
Ranatra linearis
Tenthredo koehleri
Nasonia vitripennis
Apis mellifera
Harpegnathos saltator
Acromyrmex echinatior
Tribolium castaneum
Dichochrysa prasina
Xanthostigma xanthostigma
Corydalus cornutus
Platycentropus radiatus
Bombyx mori
Ceratophyllus gallinae
Boreus hyemalis
Drosophila melanogaster
Anopheles gambiae
Aedes aegypti

**Figure S9.** Phylogeny of Polyneoptera inferred with the site-heterogeneous model LG+C60+F+G implemented in IQ-TREE, on the 102-taxon dataset.

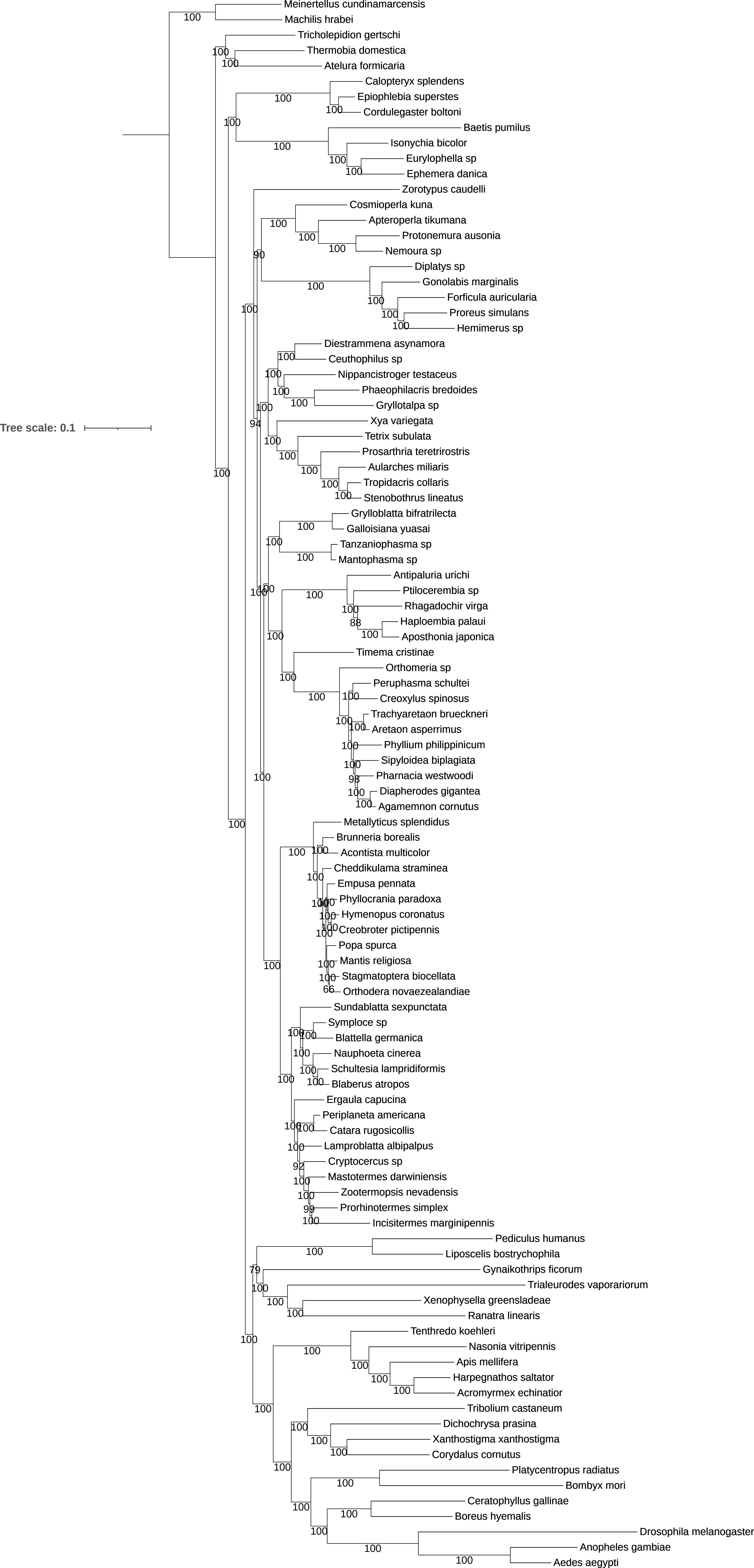

Tree scale: 0.1
Meinertellus cundinamarcensis
Machilis hrabei
Tricholepidion gertschi
Thermobia domestica
Atelura formicaria
Calopteryx splendens
Epiophlebia superstes
Cordulegaster boltoni
Baetis pumilus
Isonychia bicolor
Eurylophella sp
Ephemera danica
Zorotypus caudelli
Cosmioperla kuna
Apteroperla tikumana
Protonemura ausonia
Nemoura sp
Diplatys sp
Gonolabis marginalis
Forficula auricularia
Proreus simulans
Hemimerus sp
Diestrammena asynamora
Ceuthophilus sp
Nippancistroger testaceus
Phaeophilacris bredoides
Gryllotalpa sp
Xya variegata
Tetrix subulata
Prosarthria teretrirostris
Aularches miliaris
Tropidacris collaris
Stenobothrus lineatus
Grylloblatta bifratrilecta
Galloisiana yuasai
Tanzaniophasma sp
Mantophasma sp
Antipaluria urichi
Ptilocerembia sp
Rhagadochir virga
Haploembia palaui
Aposthonia japonica
Timema cristinae
Orthomeria sp
Peruphasma schultei
Creoxylus spinosus
Trachyaretaon brueckneri
Aretaon asperrimus
Phyllium philippinicum
Sipyloidea biplagiata
Pharnacia westwoodi
Diapherodes gigantea
Agamemnon cornutus
Metallyticus splendidus
Brunneria borealis
Acontista multicolor
Cheddikulama straminea
Empusa pennata
Phyllocrania paradoxa
Hymenopus coronatus
Creobroter pictipennis
Popa spurca
Mantis religiosa
Stagmatoptera biocellata
Orthodera novaezealandiae
Sundablatta sexpunctata
Symploce sp
Blattella germanica
Nauphoeta cinerea
Schultesia lampridiformis
Blaberus atropos
Ergaula capucina
Periplaneta americana
Catara rugosicollis
Lamproblatta albipalpus
Cryptocercus sp
Mastotermes darwiniensis
Zootermopsis nevadensis
Prorhinotermes simplex
Incisitermes marginipennis
Pediculus humanus
Liposcelis bostrychophila
Gynaikothrips ficorum
Trialeurodes vaporariorum
Xenophysella greensladeae
Ranatra linearis
Tenthredo koehleri
Nasonia vitripennis
Apis mellifera
Harpegnathos saltator
Acromyrmex echinatior
Tribolium castaneum
Dichochrysa prasina
Xanthostigma xanthostigma
Corydalus cornutus
Platycentropus radiatus
Bombyx mori
Ceratophyllus gallinae
Boreus hyemalis
Drosophila melanogaster
Anopheles gambiae
Aedes aegypti

**Figure S10.** Phylogeny of Polyneoptera inferred with the site-heterogeneous model LG+C60+F+G implemented in IQ-TREE, on the 100-taxon dataset.

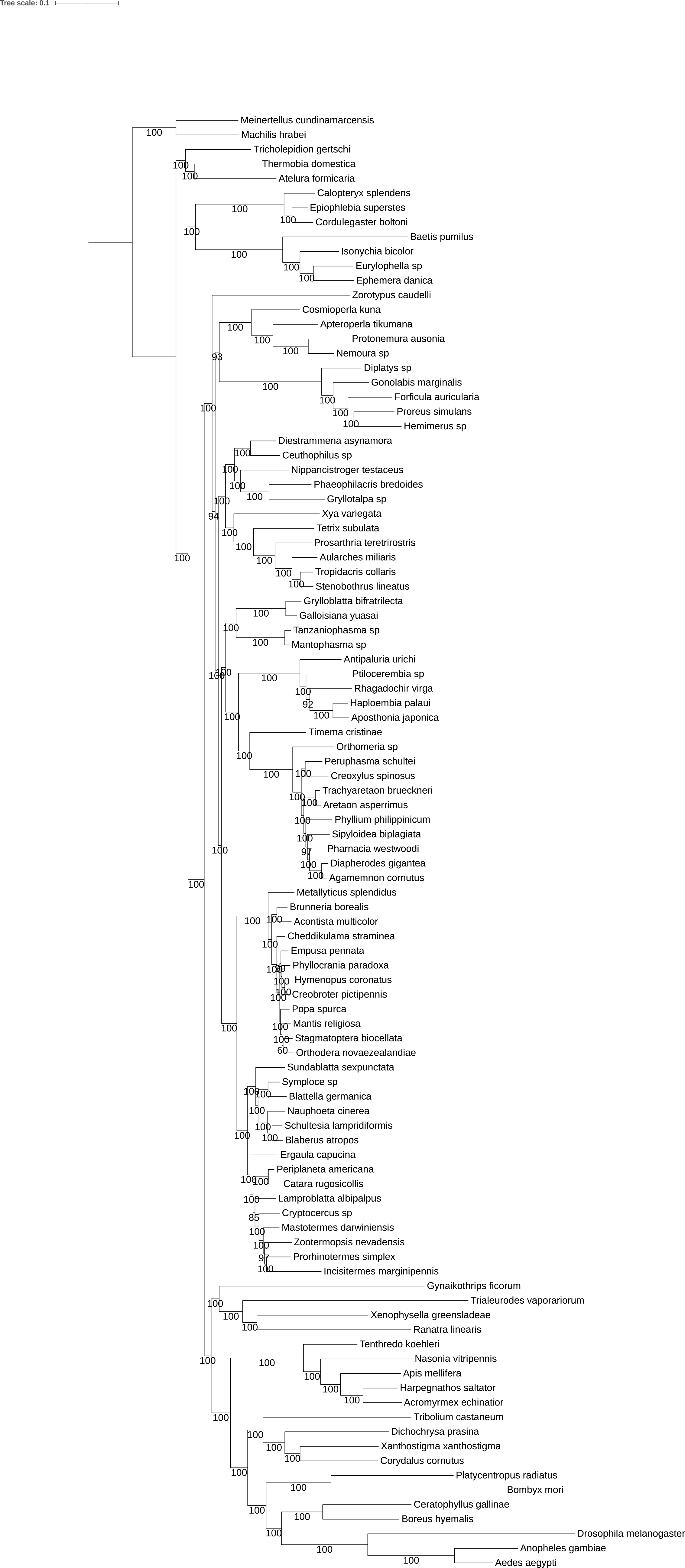

Tree scale: 0.1
Meinertellus cundinamarcensis
Machilis hrabei
Tricholepidion gertschi
Thermobia domestica
Atelura formicaria
Calopteryx splendens
Epiophlebia superstes
Cordulegaster boltoni
Baetis pumilus
Isonychia bicolor
Eurylophella sp
Ephemera danica
Zorotypus caudelli
Cosmioperla kuna
Apteroperla tikumana
Protonemura ausonia
Nemoura sp
Diplatys sp
Gonolabis marginalis
Forficula auricularia
Proreus simulans
Hemimerus sp
Diestrammena asynamora
Ceuthophilus sp
Nippancistroger testaceus
Phaeophilacris bredoides
Gryllotalpa sp
Xya variegata
Tetrix subulata
Prosarthria teretrirostris
Aularches miliaris
Tropidacris collaris
Stenobothrus lineatus
Grylloblatta bifratrilecta
Galloisiana yuasai
Tanzaniophasma sp
Mantophasma sp
Antipaluria urichi
Ptilocerembia sp
Rhagadochir virga
Haploembia palaui
Aposthonia japonica
Timema cristinae
Orthomeria sp
Peruphasma schultei
Creoxylus spinosus
Trachyaretaon brueckneri
Aretaon asperrimus
Phyllium philippinicum
Sipyloidea biplagiata
Pharnacia westwoodi
Diapherodes gigantea
Agamemnon cornutus
Metallyticus splendidus
Brunneria borealis
Acontista multicolor
Cheddikulama straminea
Empusa pennata
Phyllocrania paradoxa
Hymenopus coronatus
Creobroter pictipennis
Popa spurca
Mantis religiosa
Stagmatoptera biocellata
Orthodera novaezealandiae
Sundablatta sexpunctata
Symploce sp
Blattella germanica
Nauphoeta cinerea
Schultesia lampridiformis
Blaberus atropos
Ergaula capucina
Periplaneta americana
Catara rugosicollis
Lamproblatta albipalpus
Cryptocercus sp
Mastotermes darwiniensis
Zootermopsis nevadensis
Prorhinotermes simplex
Incisitermes marginipennis
Gynaikothrips ficorum
Trialeurodes vaporariorum
Xenophysella greensladeae
Ranatra linearis
Tenthredo koehleri
Nasonia vitripennis
Apis mellifera
Harpegnathos saltator
Acromyrmex echinatior
Tribolium castaneum
Dichochrysa prasina
Xanthostigma xanthostigma
Corydalus cornutus
Platycentropus radiatus
Bombyx mori
Ceratophyllus gallinae
Boreus hyemalis
Drosophila melanogaster
Anopheles gambiae
Aedes aegypti

**Figure S11.** Phylogeny of Polyneoptera inferred with the site-heterogeneous model LG+C60+F+G implemented in IQ-TREE, on the 33-taxon dataset.

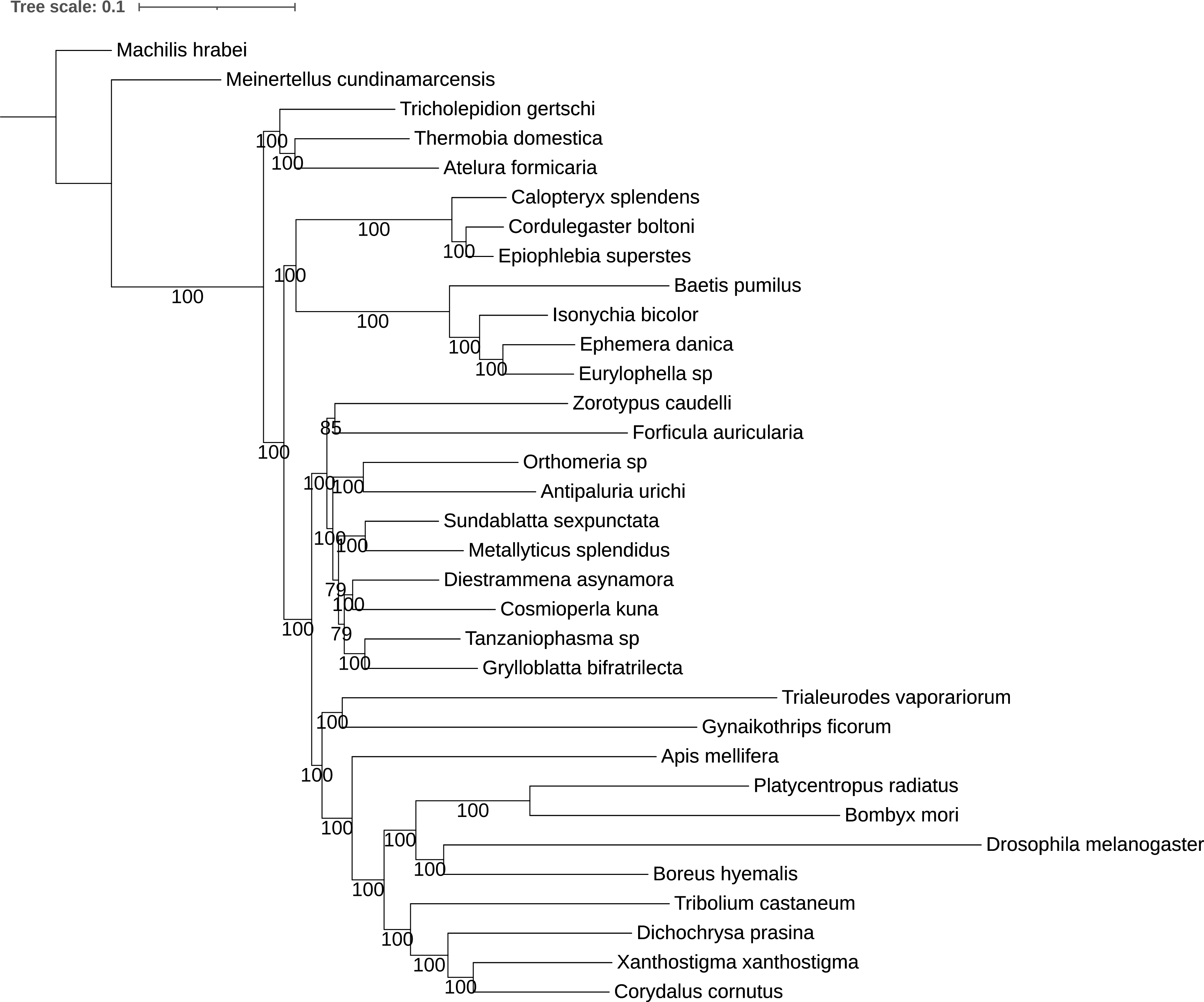
Tree scale: 0.1
Machilis hrabei
Meinertellus cundinamarcensis
Tricholepidion gertschi
Thermobia domestica
Atelura formicaria
Calopteryx splendens
Cordulegaster boltoni
Epiophlebia superstes
Baetis pumilus
Isonychia bicolor
Ephemera danica
Eurylophella sp
Zorotypus caudelli
Forficula auricularia
Orthomeria sp
Antipaluria urichi
Sundablatta sexpunctata
Metallyticus splendidus
Diestrammena asynamora
Cosmioperla kuna
Tanzaniophasma sp
Grylloblatta bifratrilecta
Trialeurodes vaporariorum
Gynaikothrips ficorum
Apis mellifera
Platycentropus radiatus
Bombyx mori
Drosophila melanogaster
Boreus hyemalis
Tribolium castaneum
Dichochrysa prasina
Xanthostigma xanthostigma
Corydalus cornutus

**Figure S12.** Phylogeny of Polyneoptera inferred with the multi-matrix mixture model LG4X implemented IQ-TREE, on the 106-taxon dataset.

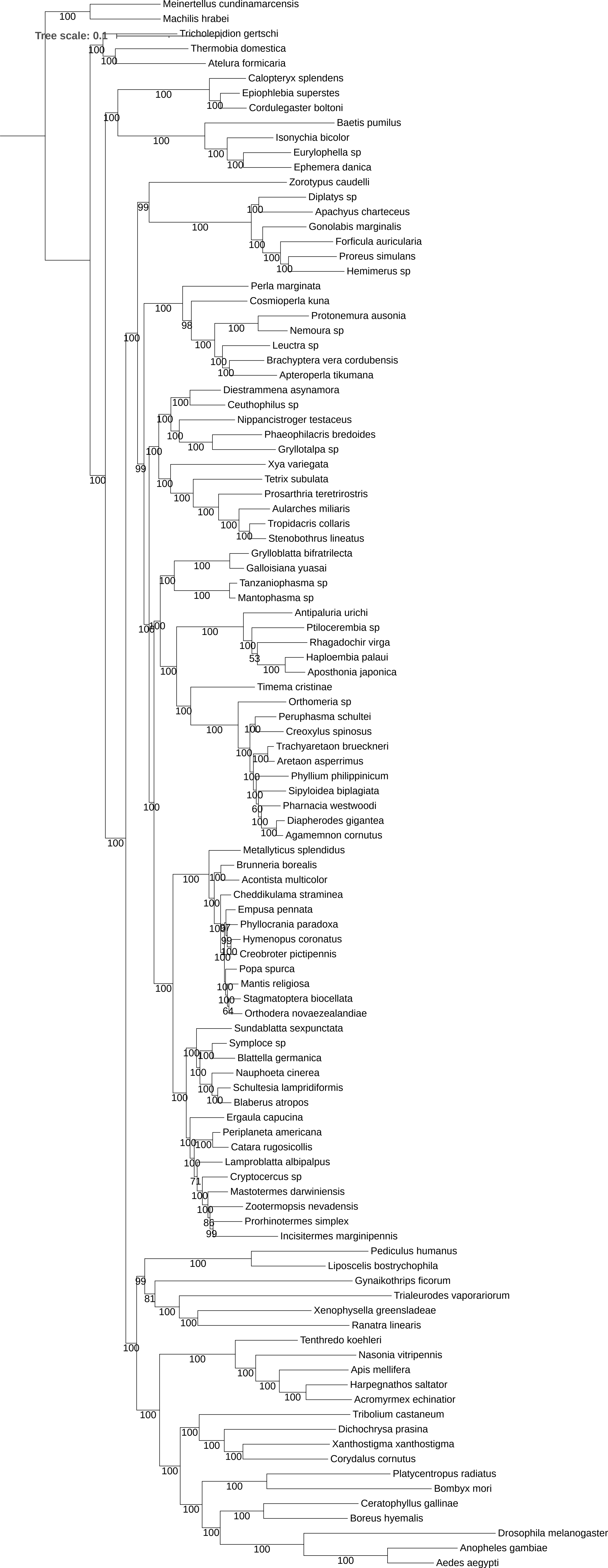
Tree scale: 0.1
Meinertellus cundinamarcensis
Machilis hrabei
Tricholepidion gertschi
Thermobia domestica
Atelura formicaria
Calopteryx splendens
Epiophlebia superstes
Cordulegaster boltoni
Baetis pumilus
Isonychia bicolor
Eurylophella sp
Ephemera danica
Zorotypus caudelli
Diplatys sp
Apachyus charteceus
Gonolabis marginalis
Forficula auricularia
Proreus simulans
Hemimerus sp
Perla marginata
Cosmioperla kuna
Protonemura ausonia
Nemoura sp
Leuctra sp
Brachyptera vera cordubensis
Apteroperla tikumana
Diestrammena asynamora
Ceuthophilus sp
Nippancistroger testaceus
Phaeophilacris bredoides
Gryllotalpa sp
Xya variegata
Tetrix subulata
Prosarthria teretrirostris
Aularches miliaris
Tropidacris collaris
Stenobothrus lineatus
Grylloblatta bifratrilecta
Galloisiana yuasai
Tanzaniophasma sp
Mantophasma sp
Antipaluria urichi
Ptilocerembia sp
Rhagadochir virga
Haploembia palaui
Aposthonia japonica
Timema cristinae
Orthomeria sp
Peruphasma schultei
Creoxylus spinosus
Trachyaretaon brueckneri
Aretaon asperrimus
Phyllium philippinicum
Sipyloidea biplagiata
Pharnacia westwoodi
Diapherodes gigantea
Agamemnon cornutus
Metallyticus splendidus
Brunneria borealis
Acontista multicolor
Cheddikulama straminea
Empusa pennata
Phyllocrania paradoxa
Hymenopus coronatus
Creobroter pictipennis
Popa spurca
Mantis religiosa
Stagmatoptera biocellata
Orthodera novaezealandiae
Sundablatta sexpunctata
Symploce sp
Blattella germanica
Nauphoeta cinerea
Schultesia lampridiformis
Blaberus atropos
Ergaula capucina
Periplaneta americana
Catara rugosicollis
Lamproblatta albipalpus
Cryptocercus sp
Mastotermes darwiniensis
Zootermopsis nevadensis
Prorhinotermes simplex
Incisitermes marginipennis
Pediculus humanus
Liposcelis bostrychophila
Gynaikothrips ficorum
Trialeurodes vaporariorum
Xenophysella greensladeae
Ranatra linearis
Tenthredo koehleri
Nasonia vitripennis
Apis mellifera
Harpegnathos saltator
Acromyrmex echinatior
Tribolium castaneum
Dichochrysa prasina
Xanthostigma xanthostigma
Corydalus cornutus
Platycentropus radiatus
Bombyx mori
Ceratophyllus gallinae
Boreus hyemalis
Drosophila melanogaster
Anopheles gambiae
Aedes aegypti

**Figure S13.** Phylogeny of Polyneoptera inferred with the multi-matrix mixture model LG4X implemented IQ-TREE, on the 102-taxon dataset.

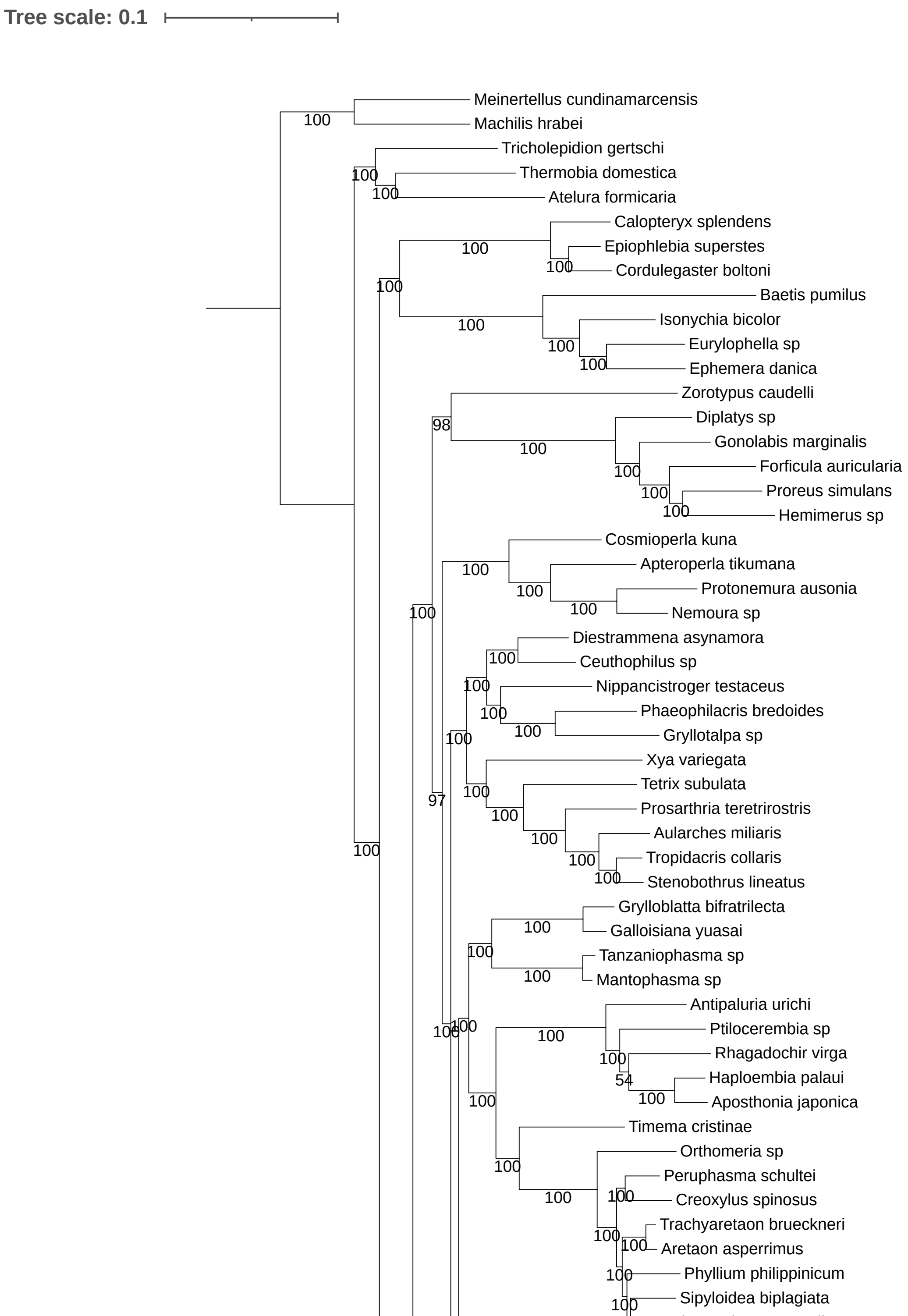

Tree scale: 0.1
Meinertellus cundinamarcensis
Machilis hrabei
Tricholepidion gertschi
Thermobia domestica
Atelura formicaria
Calopteryx splendens
Epiophlebia superstes
Cordulegaster boltoni
Baetis pumilus
Isonychia bicolor
Eurylophella sp
Ephemera danica
Zorotypus caudelli
Diplatys sp
Gonolabis marginalis
Forficula auricularia
Proreus simulans
Hemimerus sp
Cosmioperla kuna
Apteroperla tikumana
Protonemura ausonia
Nemoura sp
Diestrammena asynamora
Ceuthophilus sp
Nippancistroger testaceus
Phaeophilacris bredoides
Gryllotalpa sp
Xya variegata
Tetrix subulata
Prosarthria teretrirostris
Aularches miliaris
Tropidacris collaris
Stenobothrus lineatus
Grylloblatta bifratrilecta
Galloisiana yuasai
Tanzaniophasma sp
Mantophasma sp
Antipaluria urichi
Ptilocerembia sp
Rhagadochir virga
Haploembia palaui
Aposthonia japonica
Timema cristinae
Orthomeria sp
Peruphasma schultei
Creoxylus spinosus
Trachyaretaon brueckneri
Aretaon asperrimus
Phyllium philippinicum
Sipyloidea biplagiata

**Figure S14.** Phylogeny of Polyneoptera inferred with the multi-matrix mixture model LG4X implemented IQ-TREE, on the 100-taxon dataset.

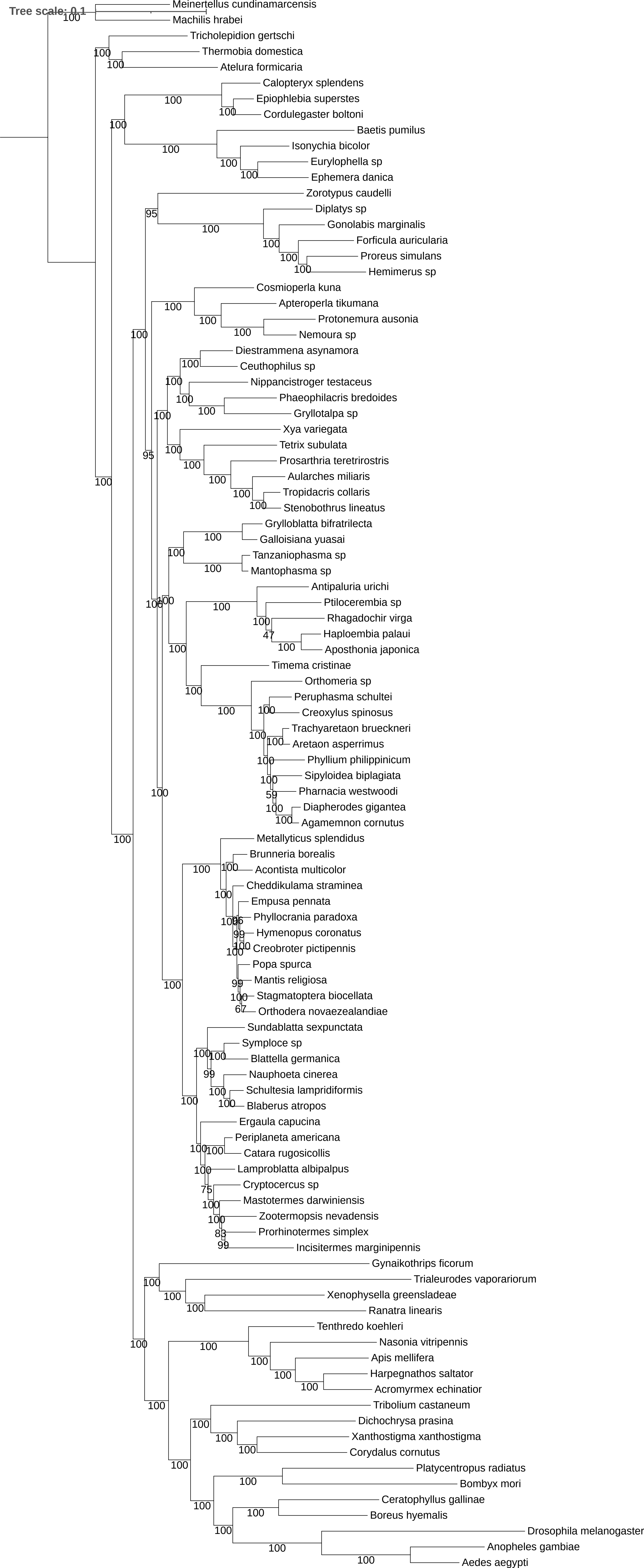
Tree scale: 0.1
Meinertellus cundinamarcensis
Machilis hrabei
Tricholepidion gertschi
Thermobia domestica
Atelura formicaria
Calopteryx splendens
Epiophlebia superstes
Cordulegaster boltoni
Baetis pumilus
Isonychia bicolor
Eurylophella sp
Ephemera danica
Zorotypus caudelli
Diplatys sp
Gonolabis marginalis
Forficula auricularia
Proreus simulans
Hemimerus sp
Cosmioperla kuna
Apteroperla tikumana
Protonemura ausonia
Nemoura sp
Diestrammena asynamora
Ceuthophilus sp
Nippancistroger testaceus
Phaeophilacris bredoides
Gryllotalpa sp
Xya variegata
Tetrix subulata
Prosarthria teretrirostris
Aularches miliaris
Tropidacris collaris
Stenobothrus lineatus
Grylloblatta bifratrilecta
Galloisiana yuasai
Tanzaniophasma sp
Mantophasma sp
Antipaluria urichi
Ptilocerembia sp
Rhagadochir virga
Haploembia palaui
Aposthonia japonica
Timema cristinae
Orthomeria sp
Peruphasma schultei
Creoxylus spinosus
Trachyaretaon brueckneri
Aretaon asperrimus
Phyllium philippinicum
Sipyloidea biplagiata
Pharnacia westwoodi
Diapherodes gigantea
Agamemnon cornutus
Metallyticus splendidus
Brunneria borealis
Acontista multicolor
Cheddikulama straminea
Empusa pennata
Phyllocrania paradoxa
Hymenopus coronatus
Creobroter pictipennis
Popa spurca
Mantis religiosa
Stagmatoptera biocellata
Orthodera novaezealandiae
Sundablatta sexpunctata
Symploce sp
Blattella germanica
Nauphoeta cinerea
Schultesia lampridiformis
Blaberus atropos
Ergaula capucina
Periplaneta americana
Catara rugosicollis
Lamproblatta albipalpus
Cryptocercus sp
Mastotermes darwiniensis
Zootermopsis nevadensis
Prorhinotermes simplex
Incisitermes marginipennis
Gynaikothrips ficorum
Trialeurodes vaporariorum
Xenophysella greensladeae
Ranatra linearis
Tenthredo koehleri
Nasonia vitripennis
Apis mellifera
Harpegnathos saltator
Acromyrmex echinatior
Tribolium castaneum
Dichochrysa prasina
Xanthostigma xanthostigma
Corydalus cornutus
Platycentropus radiatus
Bombyx mori
Ceratophyllus gallinae
Boreus hyemalis
Drosophila melanogaster
Anopheles gambiae
Aedes aegypti

**Figure S15.** Phylogeny of Polyneoptera inferred with the multi-matrix mixture model LG4X implemented IQ-TREE, on the 33-taxon dataset.

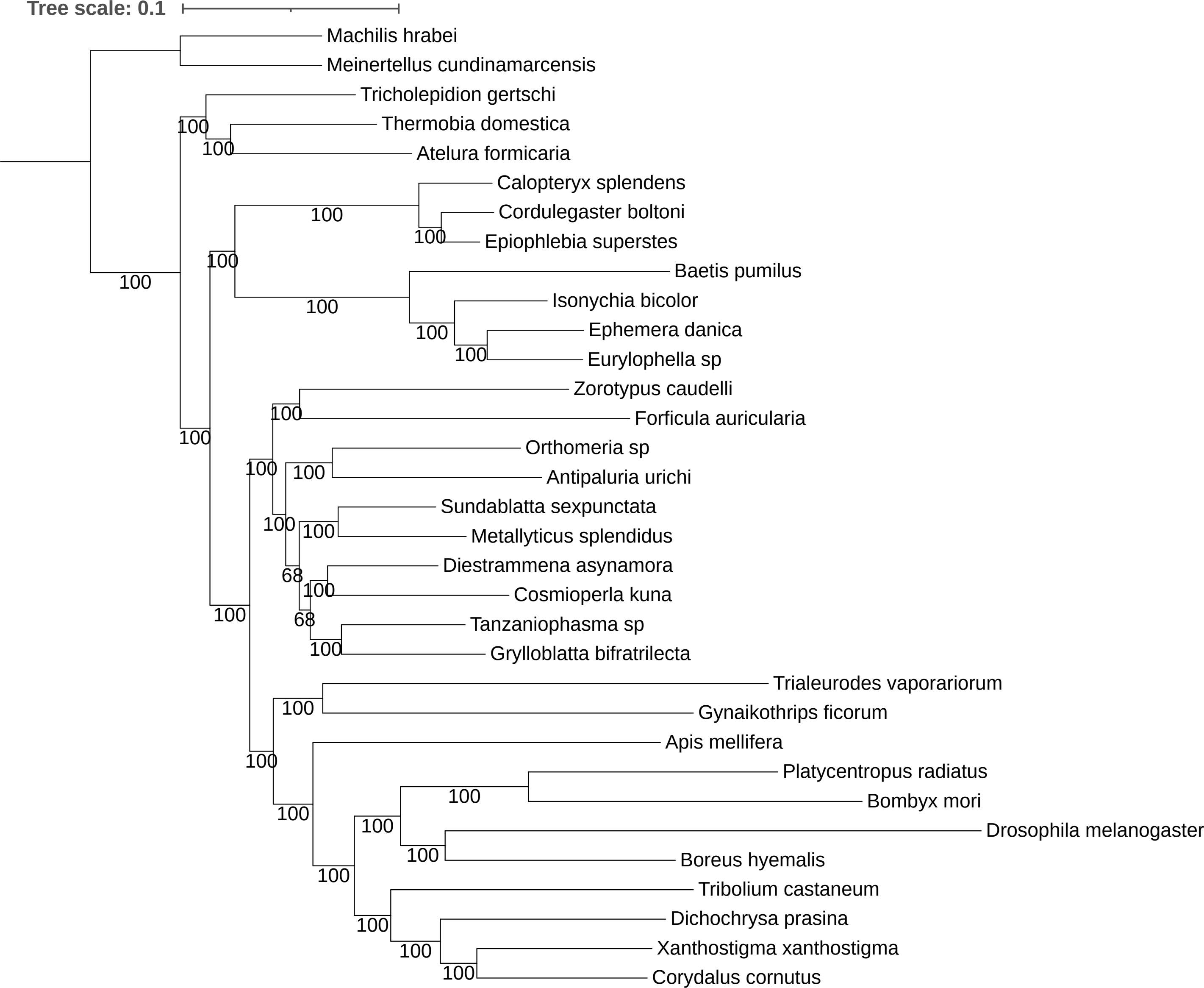

Tree scale: 0.1
Machilis hrabei
Meinertellus cundinamarcensis
Tricholepidion gertschi
Thermobia domestica
Atelura formicaria
Calopteryx splendens
Cordulegaster boltoni
Epiophlebia superstes
Baetis pumilus
Isonychia bicolor
Ephemera danica
Eurylophella sp
Zorotypus caudelli
Forficula auricularia
Orthomeria sp
Antipaluria urichi
Sundablatta sexpunctata
Metallyticus splendidus
Diestrammena asynamora
Cosmioperla kuna
Tanzaniophasma sp
Grylloblatta bifratrilecta
Trialeurodes vaporariorum
Gynaikothrips ficorum
Apis mellifera
Platycentropus radiatus
Bombyx mori
Drosophila melanogaster
Boreus hyemalis
Tribolium castaneum
Dichochrysa prasina
Xanthostigma xanthostigma
Corydalus cornutus
100
68

**Figure S16.** Phylogeny of Polyneoptera inferred with the site-homogeneous model LG implemented IQ-TREE, on the 106-taxon dataset.

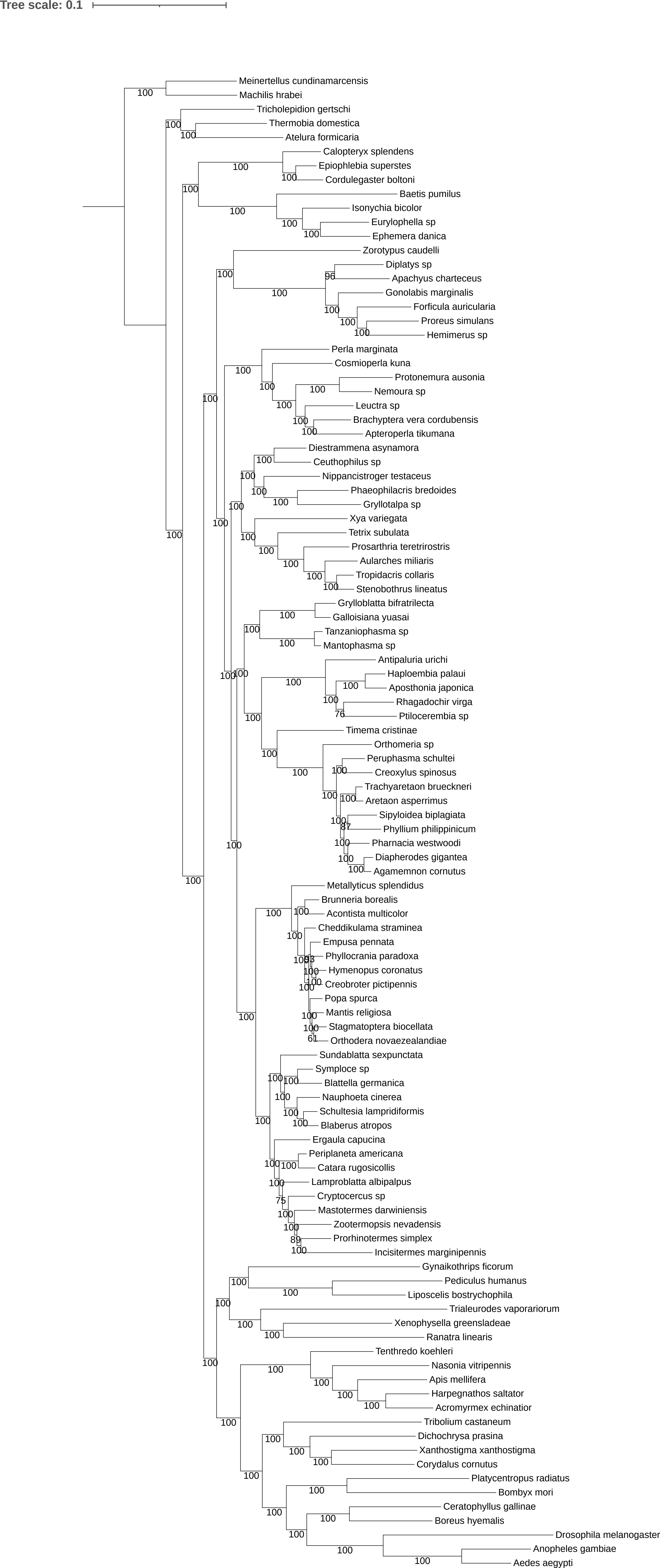
Tree scale: 0.1
Meinertellus cundinamarcensis
Machilis hrabei
Tricholepidion gertschi
Thermobia domestica
Atelura formicaria
Calopteryx splendens
Epiophlebia superstes
Cordulegaster boltoni
Baetis pumilus
Isonychia bicolor
Eurylophella sp
Ephemera danica
Zorotypus caudelli
Diplatys sp
Apachyus charteceus
Gonolabis marginalis
Forficula auricularia
Proreus simulans
Hemimerus sp
Perla marginata
Cosmioperla kuna
Protonemura ausonia
Nemoura sp
Leuctra sp
Brachyptera vera cordubensis
Apteroperla tikumana
Diestrammena asynamora
Ceuthophilus sp
Nippancistroger testaceus
Phaeophilacris bredoides
Gryllotalpa sp
Xya variegata
Tetrix subulata
Prosarthria teretrirostris
Aularches miliaris
Tropidacris collaris
Stenobothrus lineatus
Grylloblatta bifratrilecta
Galloisiana yuasai
Tanzaniophasma sp
Mantophasma sp
Antipaluria urichi
Haploembia palaui
Aposthonia japonica
Rhagadochir virga
Ptiloceembia sp
Timema cristinae
Orthomeria sp
Peruphasma schultei
Creoxylus spinosus
Trachyaretaon brueckneri
Aretaon asperrimus
Sipyloidea biplagiata
Phyllium philippinicum
Pharnacia westwoodi
Diapherodes gigantea
Agamemnon cornutus
Metallyticus splendidus
Brunneria borealis
Acontista multicolor
Cheddikulama straminea
Empusa pennata
Phyllocrania paradoxa
Hymenopus coronatus
Creobroter pictipennis
Popa spurca
Mantis religiosa
Stagmatoptera biocellata
Orthodera novaezealandiae
Sundablatta sexpunctata
Symploce sp
Blattella germanica
Nauphoeta cinerea
Schultesia lampridiformis
Blaberus atropos
Ergaula capucina
Periplaneta americana
Catara rugosicollis
Lamproblatta albipalpus
Cryptocercus sp
Mastotermes darwiniensis
Zootermopsis nevadensis
Prorhinotermes simplex
Incisitermes marginipennis
Gynaikothrips ficorum
Pediculus humanus
Liposcelis bostrychophila
Trialeurodes vaporariorum
Xenophysella greensladeae
Ranatra linearis
Tenthredo koehleri
Nasonia vitripennis
Apis mellifera
Harpegnathos saltator
Acromyrmex echinatior
Tribolium castaneum
Dichochrysa prasina
Xanthostigma xanthostigma
Corydalus cornutus
Platycentropus radiatus
Bombyx mori
Ceratophyllus gallinae
Boreus hyemalis
Drosophila melanogaster
Anopheles gambiae
Aedes aegypti

**Figure S17.** Phylogeny of Polyneoptera inferred with the multi-matrix mixture model LG implemented IQ-TREE, on the 102-taxon dataset.

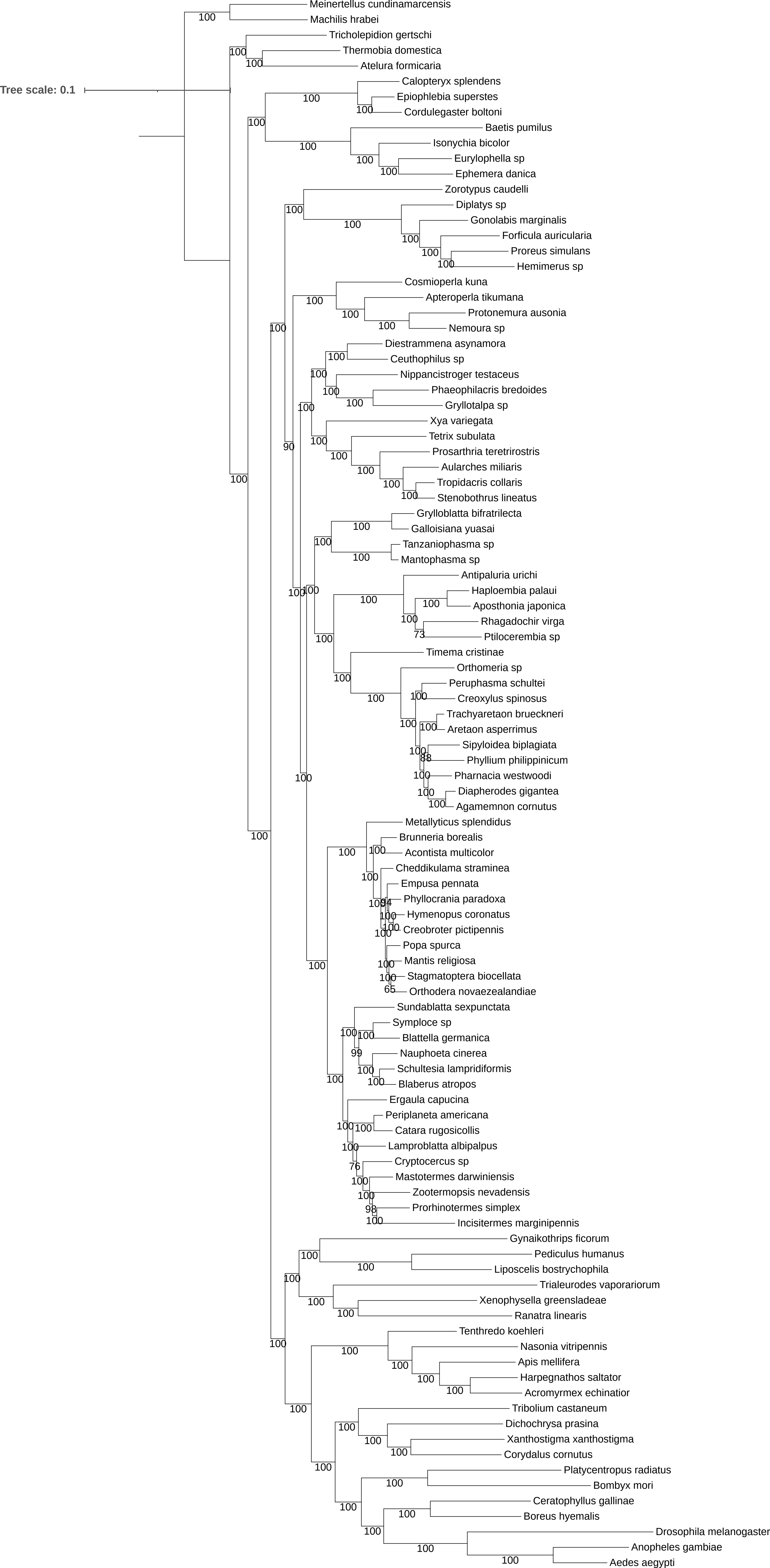

Tree scale: 0.1
Meinertellus cundinamarcensis
Machilis hrabei
Tricholepidion gertschi
Thermobia domestica
Atelura formicaria
Calopteryx splendens
Epiophlebia superstes
Cordulegaster boltoni
Baetis pumilus
Isonychia bicolor
Eurylophella sp
Ephemera danica
Zorotypus caudelli
Diplatys sp
Gonolabis marginalis
Forficula auricularia
Proreus simulans
Hemimerus sp
Cosmioperla kuna
Apteroperla tikumana
Protonemura ausonia
Nemoura sp
Diestrammena asynamora
Ceuthophilus sp
Nippancistroger testaceus
Phaeophilacris bredoides
Gryllotalpa sp
Xya variegata
Tetrix subulata
Prosarthria teretrirostris
Aularches miliaris
Tropidacris collaris
Stenobothrus lineatus
Grylloblatta bifratrilecta
Galloisiana yuasai
Tanzaniophasma sp
Mantophasma sp
Antipaluria urichi
Haploembia palaui
Aposthonia japonica
Rhagadochir virga
Ptilocerembia sp
Timema cristinae
Orthomeria sp
Peruphasma schultei
Creoxylus spinosus
Trachyaretaon brueckneri
Aretaon asperrimus
Sipyloidea biplagiata
Phyllium philippinicum
Pharnacia westwoodi
Diapherodes gigantea
Agamemnon cornutus
Metallyticus splendidus
Brunneria borealis
Acontista multicolor
Cheddikulama straminea
Empusa pennata
Phyllocrania paradoxa
Hymenopus coronatus
Creobroter pictipennis
Popa spurca
Mantis religiosa
Stagmatoptera biocellata
Orthodera novaezealandiae
Sundablatta sexpunctata
Symploce sp
Blattella germanica
Nauphoeta cinerea
Schultesia lampridiformis
Blaberus atropos
Ergaula capucina
Periplaneta americana
Catara rugosicollis
Lamproblatta albipalpus
Cryptocercus sp
Mastotermes darwiniensis
Zootermopsis nevadensis
Prorhinotermes simplex
Incisitermes marginipennis
Gynaikothrips ficorum
Pediculus humanus
Liposcelis bostrychophila
Trialeurodes vaporariorum
Xenophysella greensladeae
Ranatra linearis
Tenthredo koehleri
Nasonia vitripennis
Apis mellifera
Harpegnathos saltator
Acromyrmex echinatior
Tribolium castaneum
Dichochrysa prasina
Xanthostigma xanthostigma
Corydalus cornutus
Platycentropus radiatus
Bombyx mori
Ceratophyllus gallinae
Boreus hyemalis
Drosophila melanogaster
Anopheles gambiae
Aedes aegypti

**Figure S18.** Phylogeny of Polyneoptera inferred with the multi-matrix mixture model LG implemented IQ-TREE, on the 100-taxon dataset.

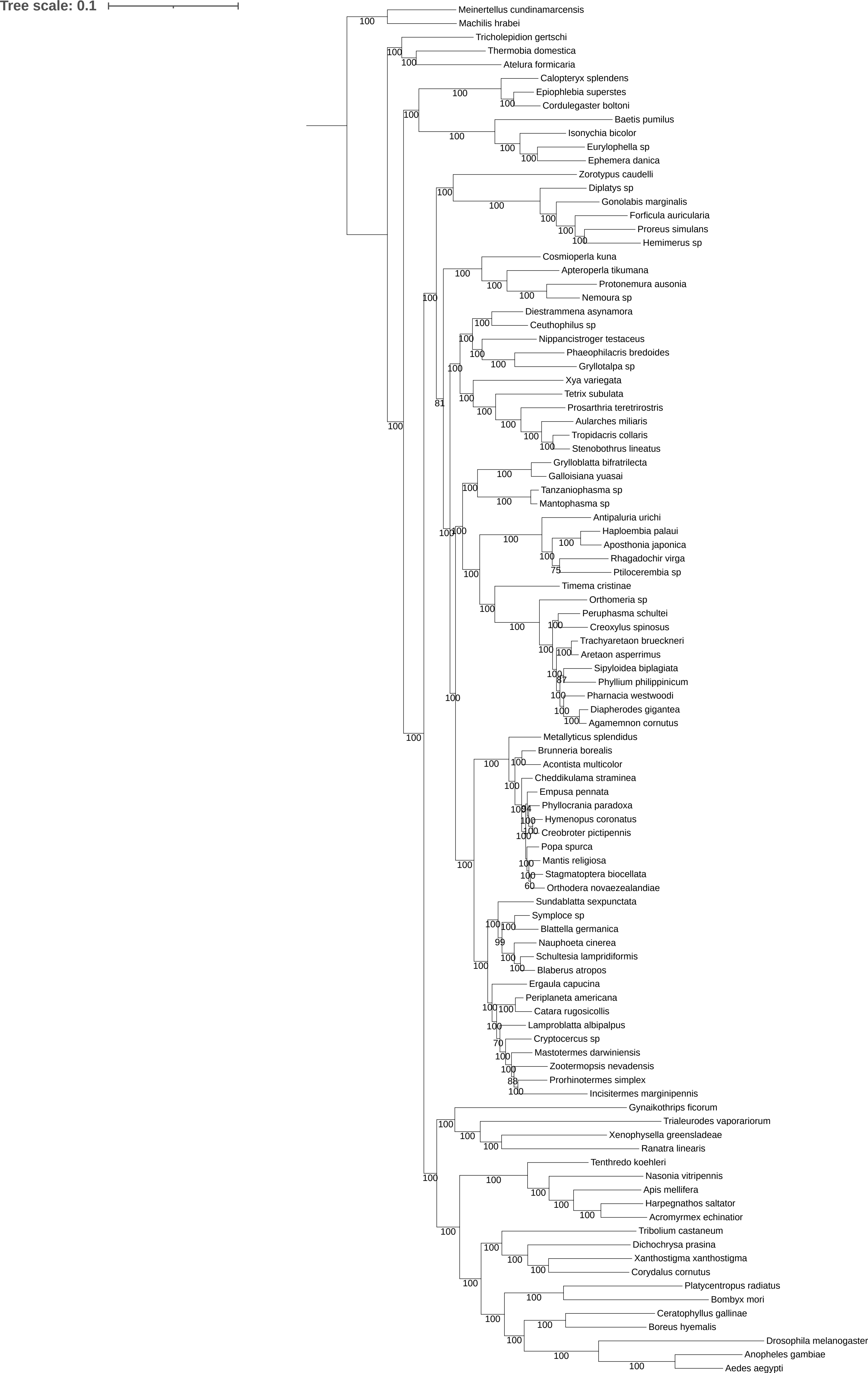
Tree scale: 0.1
Meinertellus cundinamarcensis
Machilis hrabei
Tricholepidion gertschi
Thermobia domestica
Atelura formicaria
Calopteryx splendens
Epiophlebia superstes
Cordulegaster boltoni
Baetis pumilus
Isonychia bicolor
Eurylophella sp
Ephemera danica
Zorotypus caudelli
Diplatys sp
Gonolabis marginalis
Forficula auricularia
Proreus simulans
Hemimerus sp
Cosmioperla kuna
Apteroperla tikumana
Protonemura ausonia
Nemoura sp
Diestrammena asynamora
Ceuthophilus sp
Nippancistroger testaceus
Phaeophilacris bredoides
Gryllotalpa sp
Xya variegata
Tetrix subulata
Prosarthria teretrirostris
Aularches miliaris
Tropidacris collaris
Stenobothrus lineatus
Grylloblatta bifratrilecta
Galloisiana yuasai
Tanzaniophasma sp
Mantophasma sp
Antipaluria urichi
Haploembia palaui
Aposthonia japonica
Rhagadochir virga
Ptilocerembia sp
Timema cristinae
Orthomeria sp
Peruphasma schultei
Creoxylus spinosus
Trachyaretaon brueckneri
Aretaon asperrimus
Sipyloidea biplagiata
Phyllium philippinicum
Pharnacia westwoodi
Diapherodes gigantea
Agamemnon cornutus
Metallyticus splendidus
Brunneria borealis
Acontista multicolor
Cheddikulama straminea
Empusa pennata
Phyllocrania paradoxa
Hymenopus coronatus
Creobroter pictipennis
Popa spurca
Mantis religiosa
Stagmatoptera biocellata
Orthodera novaezealandiae
Sundablatta sexpunctata
Symploce sp
Blattella germanica
Nauphoeta cinerea
Schultesia lampridiformis
Blaberus atropos
Ergaula capucina
Periplaneta americana
Catara rugosicollis
Lamproblatta albipalpus
Cryptocercus sp
Mastotermes darwiniensis
Zootermopsis nevadensis
Prorhinotermes simplex
Incisitermes marginipennis
Gynaikothrips ficorum
Trialeurodes vaporariorum
Xenophysella greensladeae
Ranatra linearis
Tenthredo koehleri
Nasonia vitripennis
Apis mellifera
Harpegnathos saltator
Acromyrmex echinatior
Tribolium castaneum
Dichochrysa prasina
Xanthostigma xanthostigma
Corydalus cornutus
Platycentropus radiatus
Bombyx mori
Ceratophyllus gallinae
Boreus hyemalis
Drosophila melanogaster
Anopheles gambiae
Aedes aegypti

**Figure S19.** Phylogeny of Polyneoptera inferred with the multi-matrix mixture model LG implemented IQ-TREE, on the 33-taxon dataset.

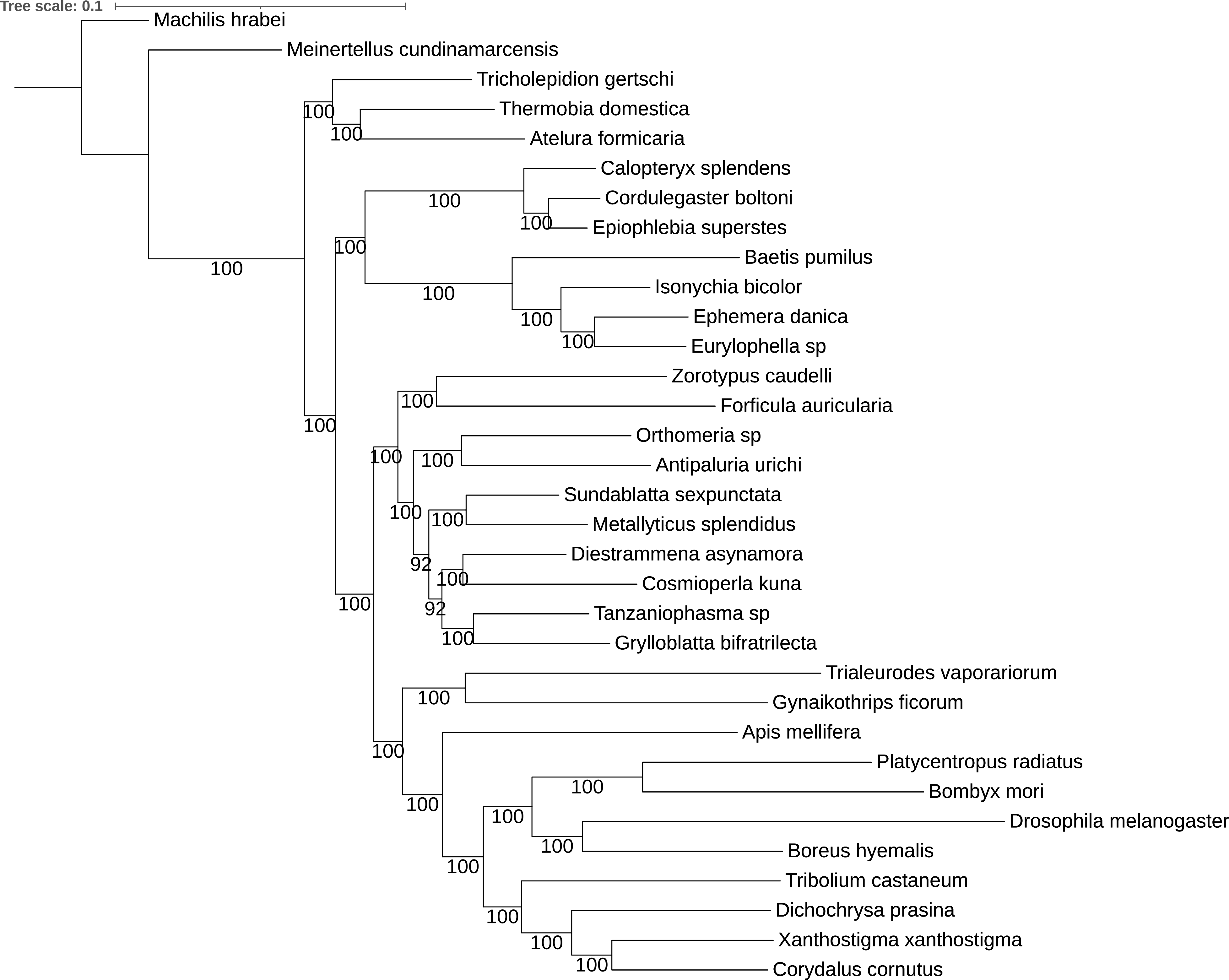

Tree scale: 0.1
Machilis hrabei
Meinertellus cundinamarcensis
Tricholepidion gertschi
Thermobia domestica
Atelura formicaria
Calopteryx splendens
Cordulegaster boltoni
Epiophlebia superstes
Baetis pumilus
Isonychia bicolor
Ephemera danica
Eurylophella sp
Zorotypus caudelli
Forficula auricularia
Orthomeria sp
Antipaluria urichi
Sundablatta sexpunctata
Metallyticus splendidus
Diestrammena asynamora
Cosmioperla kuna
Tanzaniophasma sp
Grylloblatta bifratrilecta
Trialeurodes vaporariorum
Gynaikothrips ficorum
Apis mellifera
Platycentropus radiatus
Bombyx mori
Drosophila melanogaster
Boreus hyemalis
Tribolium castaneum
Dichochrysa prasina
Xanthostigma xanthostigma
Corydalus cornutus
100
92

**Figure S20.** Phylogeny of Polyneoptera inferred with the site-homogeneous model GTR+G implemented IQ-TREE, on the 106-taxon dataset.

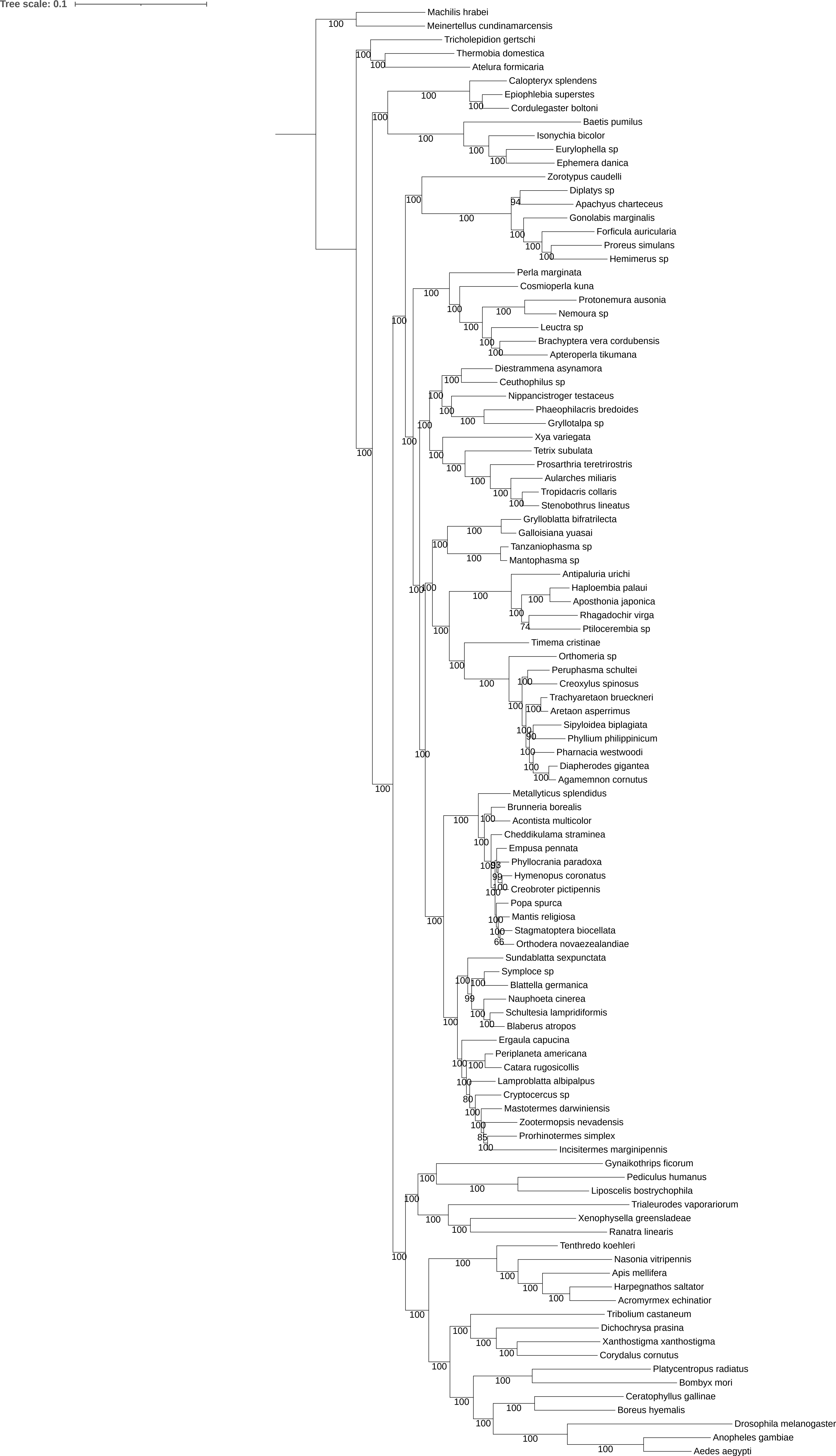

Tree scale: 0.1
Machilis hrabei
Meinertellus cundinamarcensis
Tricholepidion gertschi
Thermobia domestica
Atelura formicaria
Calopteryx splendens
Epiophlebia superstes
Cordulegaster boltoni
Baetis pumilus
Isonychia bicolor
Eurylophella sp
Ephemera danica
Zorotypus caudelli
Diplatys sp
Apachyus charteceus
Gonolabis marginalis
Forficula auricularia
Proreus simulans
Hemimerus sp
Perla marginata
Cosmioperla kuna
Protonemura ausonia
Nemoura sp
Leuctra sp
Brachyptera vera cordubensis
Apteroperla tikumana
Diestrammena asynamora
Ceuthophilus sp
Nippancistroger testaceus
Phaeophilacris bredoides
Gryllotalpa sp
Xya variegata
Tetrix subulata
Prosarthria teretrirostris
Aularches miliaris
Tropidacris collaris
Stenobothrus lineatus
Grylloblatta bifratrilecta
Galloisiana yuasai
Tanzaniophasma sp
Mantophasma sp
Antipaluria urichi
Haploembia palaui
Aposthonia japonica
Rhagadochir virga
Ptilocerembia sp
Timema cristinae
Orthomeria sp
Peruphasma schultei
Creoxylus spinosus
Trachyaretaon brueckneri
Aretaon asperrimus
Sipyloidea biplagiata
Phyllium philippinicum
Pharnacia westwoodi
Diapherodes gigantea
Agamemnon cornutus
Metallyticus splendidus
Brunneria borealis
Acontista multicolor
Cheddikulama straminea
Empusa pennata
Phyllocrania paradoxa
Hymenopus coronatus
Creobroter pictipennis
Popa spurca
Mantis religiosa
Stagmatoptera biocellata
Orthodera novaezealandiae
Sundablatta sexpunctata
Symploce sp
Blattella germanica
Nauphoeta cinerea
Schultesia lampridiformis
Blaberus atropos
Ergaula capucina
Periplaneta americana
Catara rugosicollis
Lamproblatta albipalpus
Cryptocercus sp
Mastotermes darwiniensis
Zootermopsis nevadensis
Prorhinotermes simplex
Incisitermes marginipennis
Gynaikothrips ficorum
Pediculus humanus
Liposcelis bostrychophila
Trialeurodes vaporariorum
Xenophysella greensladeae
Ranatra linearis
Tenthredo koehleri
Nasonia vitripennis
Apis mellifera
Harpegnathos saltator
Acromyrmex echinatior
Tribolium castaneum
Dichochrysa prasina
Xanthostigma xanthostigma
Corydalus cornutus
Platycentropus radiatus
Bombyx mori
Ceratophyllus gallinae
Boreus hyemalis
Drosophila melanogaster
Anopheles gambiae
Aedes aegypti

**Figure S21.** Phylogeny of Polyneoptera inferred with the multi-matrix mixture model GTR+G implemented IQ-TREE, on the 102-taxon dataset.

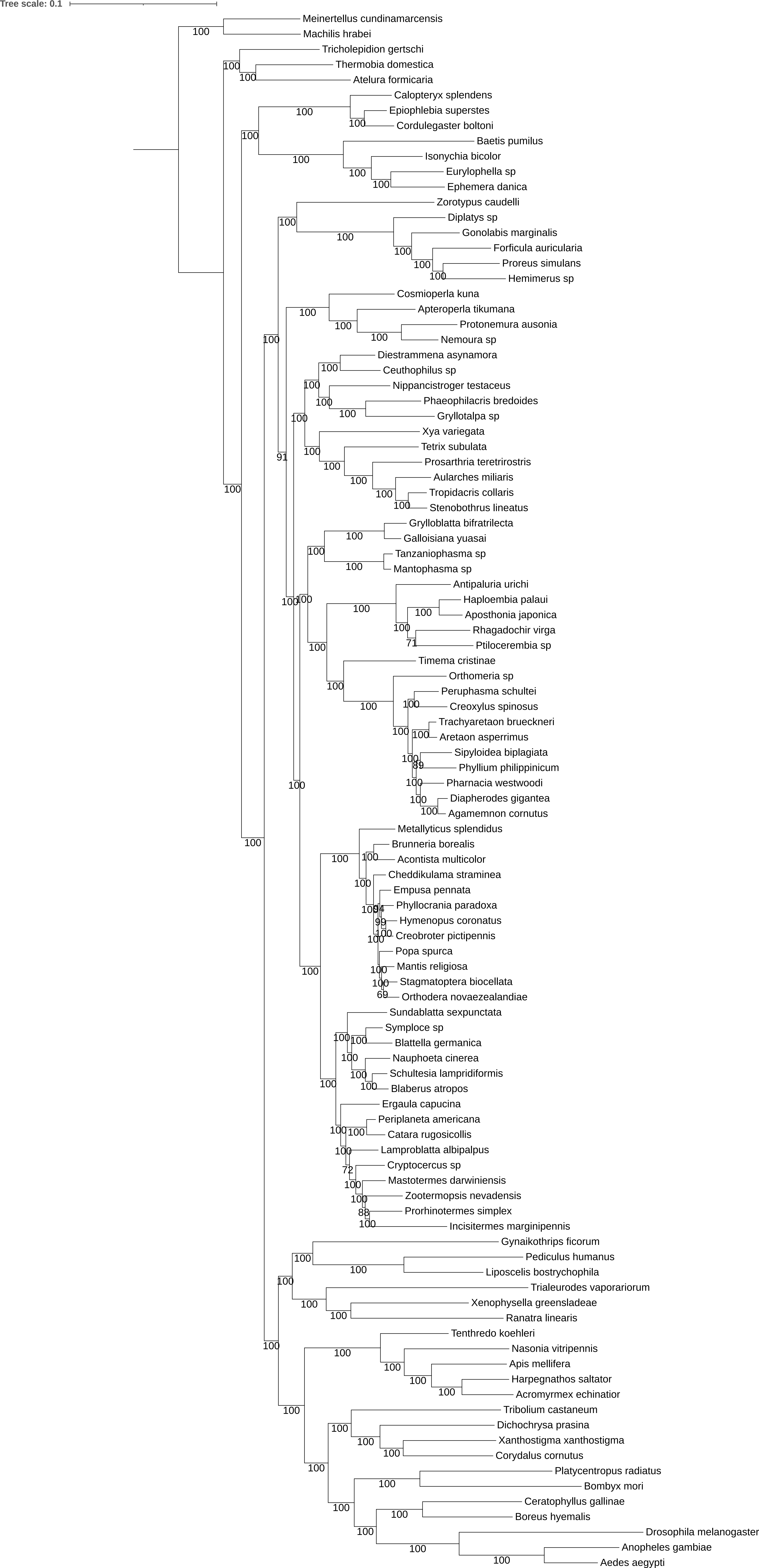
Tree scale: 0.1
Meinertellus cundinamarcensis
Machilis hrabei
Tricholepidion gertschi
Thermobia domestica
Atelura formicaria
Calopteryx splendens
Epiophlebia superstes
Cordulegaster boltoni
Baetis pumilus
Isonychia bicolor
Eurylophella sp
Ephemera danica
Zorotypus caudelli
Diplatys sp
Gonolabis marginalis
Forficula auricularia
Proreus simulans
Hemimerus sp
Cosmioperla kuna
Apteroperla tikumana
Protonemura ausonia
Nemoura sp
Diestrammena asynamora
Ceuthophilus sp
Nippancistroger testaceus
Phaeophilacris bredoides
Gryllotalpa sp
Xya variegata
Tetrix subulata
Prosarthria teretrirostris
Aularches miliaris
Tropidacris collaris
Stenobothrus lineatus
Grylloblatta bifratrilecta
Galloisiana yuasai
Tanzaniophasma sp
Mantophasma sp
Antipaluria urichi
Haploembia palaui
Aposthonia japonica
Rhagadochir virga
Ptilocerembia sp
Timema cristinae
Orthomeria sp
Peruphasma schultei
Creoxylus spinosus
Trachyaretaon brueckneri
Aretaon asperrimus
Sipyloidea biplagiata
Phyllium philippinicum
Pharnacia westwoodi
Diapherodes gigantea
Agamemnon cornutus
Metallyticus splendidus
Brunneria borealis
Acontista multicolor
Cheddikulama straminea
Empusa pennata
Phyllocrania paradoxa
Hymenopus coronatus
Creobroter pictipennis
Popa spurca
Mantis religiosa
Stagmatoptera biocellata
Orthodera novaezealandiae
Sundablatta sexpunctata
Symploce sp
Blattella germanica
Nauphoeta cinerea
Schultesia lampridiformis
Blaberus atropos
Ergaula capucina
Periplaneta americana
Catara rugosicollis
Lamproblatta albipalpus
Cryptocercus sp
Mastotermes darwiniensis
Zootermopsis nevadensis
Prorhinotermes simplex
Incisitermes marginipennis
Gynaikothrips ficorum
Pediculus humanus
Liposcelis bostrychophila
Trialeurodes vaporariorum
Xenophysella greensladeae
Ranatra linearis
Tenthredo koehleri
Nasonia vitripennis
Apis mellifera
Harpegnathos saltator
Acromyrmex echinatior
Tribolium castaneum
Dichochrysa prasina
Xanthostigma xanthostigma
Corydalus cornutus
Platycentropus radiatus
Bombyx mori
Ceratophyllus gallinae
Boreus hyemalis
Drosophila melanogaster
Anopheles gambiae
Aedes aegypti

**Figure S22.** Phylogeny of Polyneoptera inferred with the multi-matrix mixture model GTR+G implemented IQ-TREE, on the 100-taxon dataset.

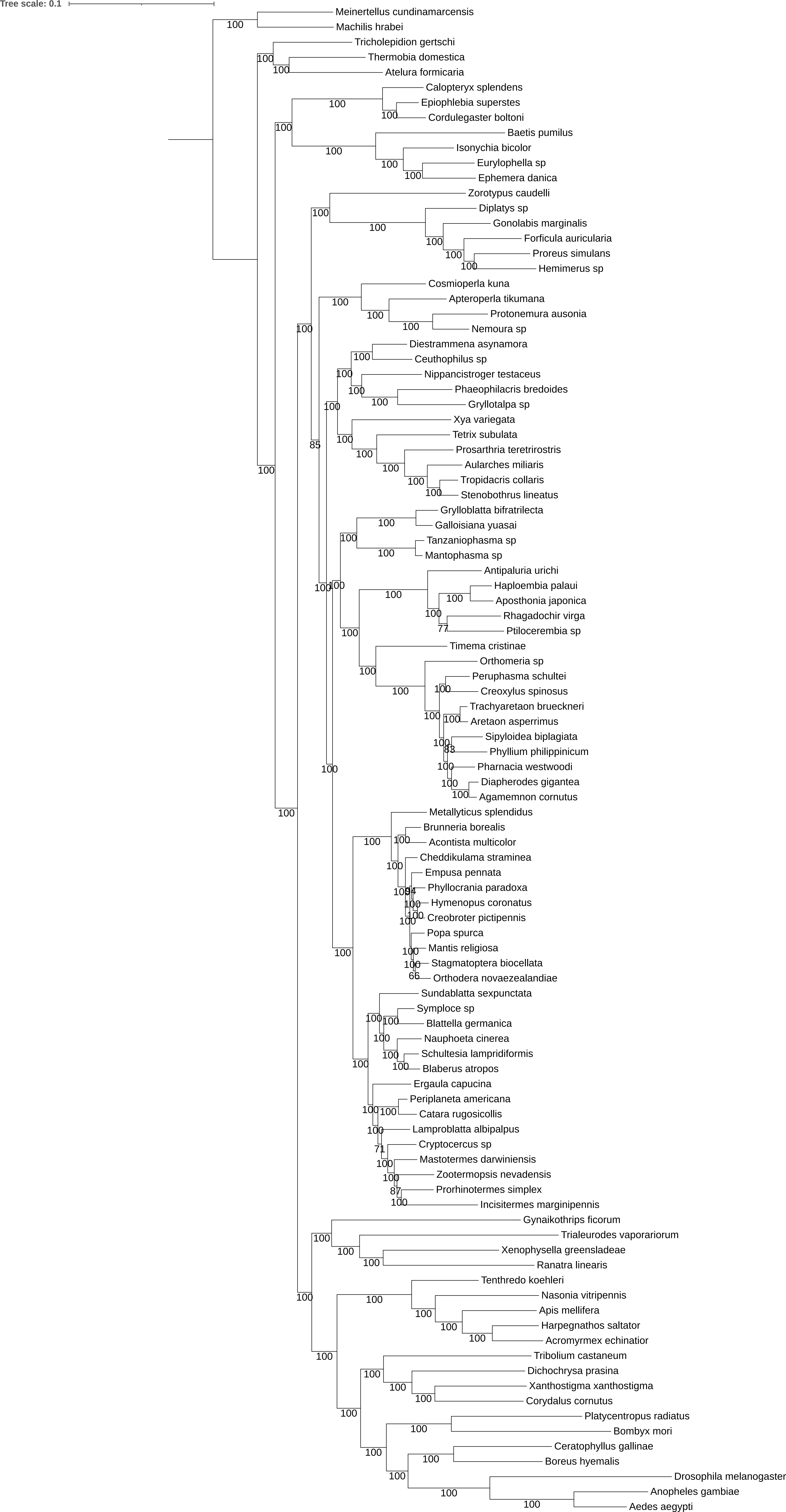
Tree scale: 0.1
Meinertellus cundinamarcensis
Machilis hrabei
Tricholepidion gertschi
Thermobia domestica
Atelura formicaria
Calopteryx splendens
Epiophlebia superstes
Cordulegaster boltoni
Baetis pumilus
Isonychia bicolor
Eurylophella sp
Ephemera danica
Zorotypus caudelli
Diplatys sp
Gonolabis marginalis
Forficula auricularia
Proreus simulans
Hemimerus sp
Cosmioperla kuna
Apteroperla tikumana
Protonemura ausonia
Nemoura sp
Diestrammena asynamora
Ceuthophilus sp
Nippancistroger testaceus
Phaeophilacris bredoides
Gryllotalpa sp
Xya variegata
Tetrix subulata
Prosarthria teretrirostris
Aularches miliaris
Tropidacris collaris
Stenobothrus lineatus
Grylloblatta bifratrilecta
Galloisiana yuasai
Tanzaniophasma sp
Mantophasma sp
Antipaluria urichi
Haploembia palaui
Aposthonia japonica
Rhagadochir virga
Ptilocerembia sp
Timema cristinae
Orthomeria sp
Peruphasma schultei
Creoxylus spinosus
Trachyaretaon brueckneri
Aretaon asperrimus
Sipyloidea biplagiata
Phyllium philippinicum
Pharnacia westwoodi
Diapherodes gigantea
Agamemnon cornutus
Metallyticus splendidus
Brunneria borealis
Acontista multicolor
Cheddikulama straminea
Empusa pennata
Phyllocrania paradoxa
Hymenopus coronatus
Creobroter pictipennis
Popa spurca
Mantis religiosa
Stagmatoptera biocellata
Orthodera novaezealandiae
Sundablatta sexpunctata
Symploce sp
Blattella germanica
Nauphoeta cinerea
Schultesia lampridiformis
Blaberus atropos
Ergaula capucina
Periplaneta americana
Catara rugosicollis
Lamproblatta albipalpus
Cryptocercus sp
Mastotermes darwiniensis
Zootermopsis nevadensis
Prorhinotermes simplex
Incisitermes marginipennis
Gynaikothrips ficorum
Trialeurodes vaporariorum
Xenophysella greensladeae
Ranatra linearis
Tenthredo koehleri
Nasonia vitripennis
Apis mellifera
Harpegnathos saltator
Acromyrmex echinatior
Tribolium castaneum
Dichochrysa prasina
Xanthostigma xanthostigma
Corydalus cornutus
Platycentropus radiatus
Bombyx mori
Ceratophyllus gallinae
Boreus hyemalis
Drosophila melanogaster
Anopheles gambiae
Aedes aegypti

**Figure S23.** Phylogeny of Polyneoptera inferred with the multi-matrix mixture model GTR+G implemented IQ-TREE, on the 33-taxon dataset.

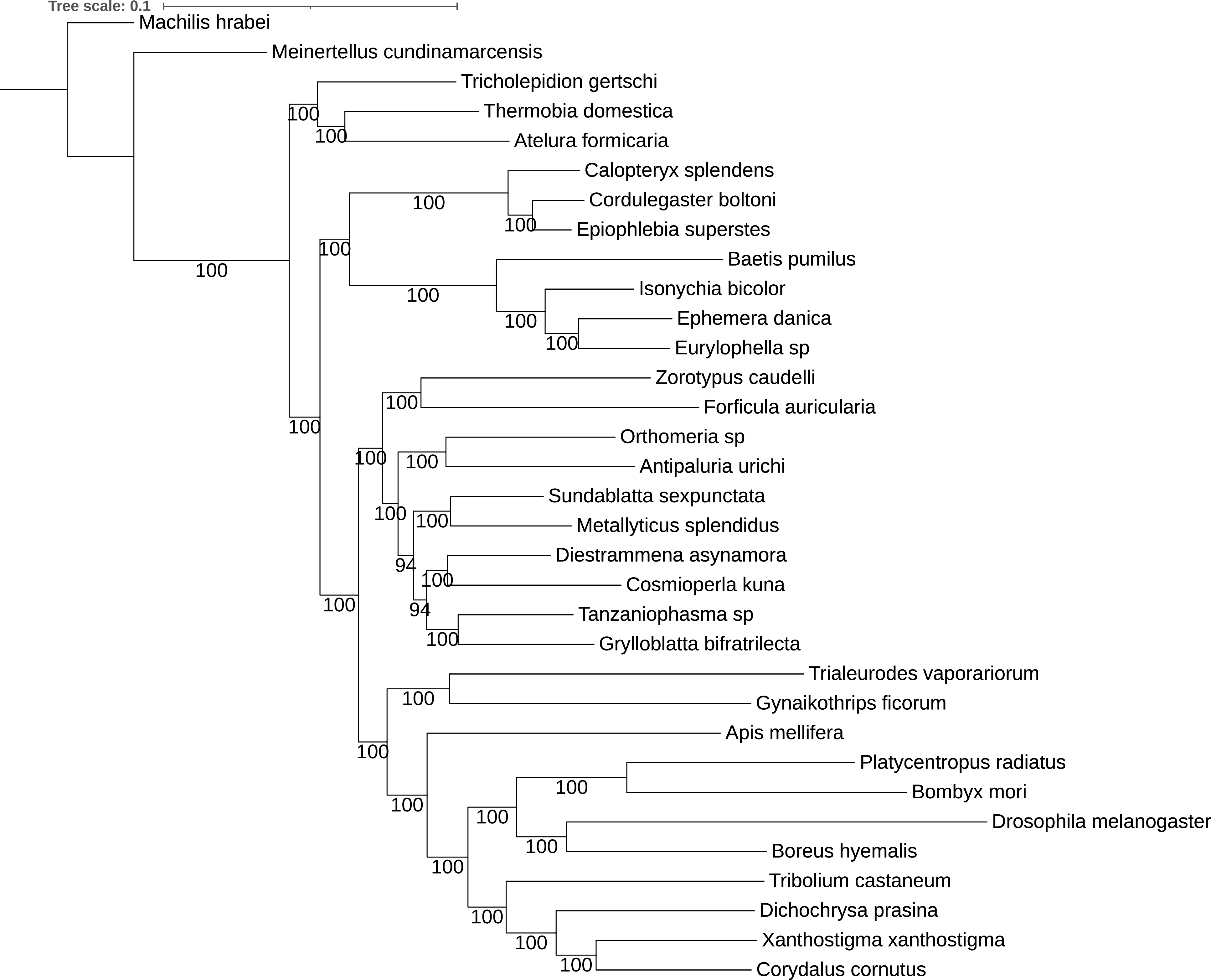
Tree scale: 0.1
Machilis hrabei
Meinertellus cundinamarcensis
Tricholepidion gertschi
Thermobia domestica
Atelura formicaria
Calopteryx splendens
Cordulegaster boltoni
Epiophlebia superstes
Baetis pumilus
Isonychia bicolor
Ephemera danica
Eurylophella sp
Zorotypus caudelli
Forficula auricularia
Orthomeria sp
Antipaluria urichi
Sundablatta sexpunctata
Metallyticus splendidus
Diestrammena asynamora
Cosmioperla kuna
Tanzaniophasma sp
Grylloblatta bifratrilecta
Trialeurodes vaporariorum
Gynaikothrips ficorum
Apis mellifera
Platycentropus radiatus
Bombyx mori
Drosophila melanogaster
Boreus hyemalis
Tribolium castaneum
Dichochrysa prasina
Xanthostigma xanthostigma
Corydalus cornutus
100
94

**Figure S24.** Dated phylogeny of Polyneoptera, inferred with the autocorrelated rates (AC) clock model.

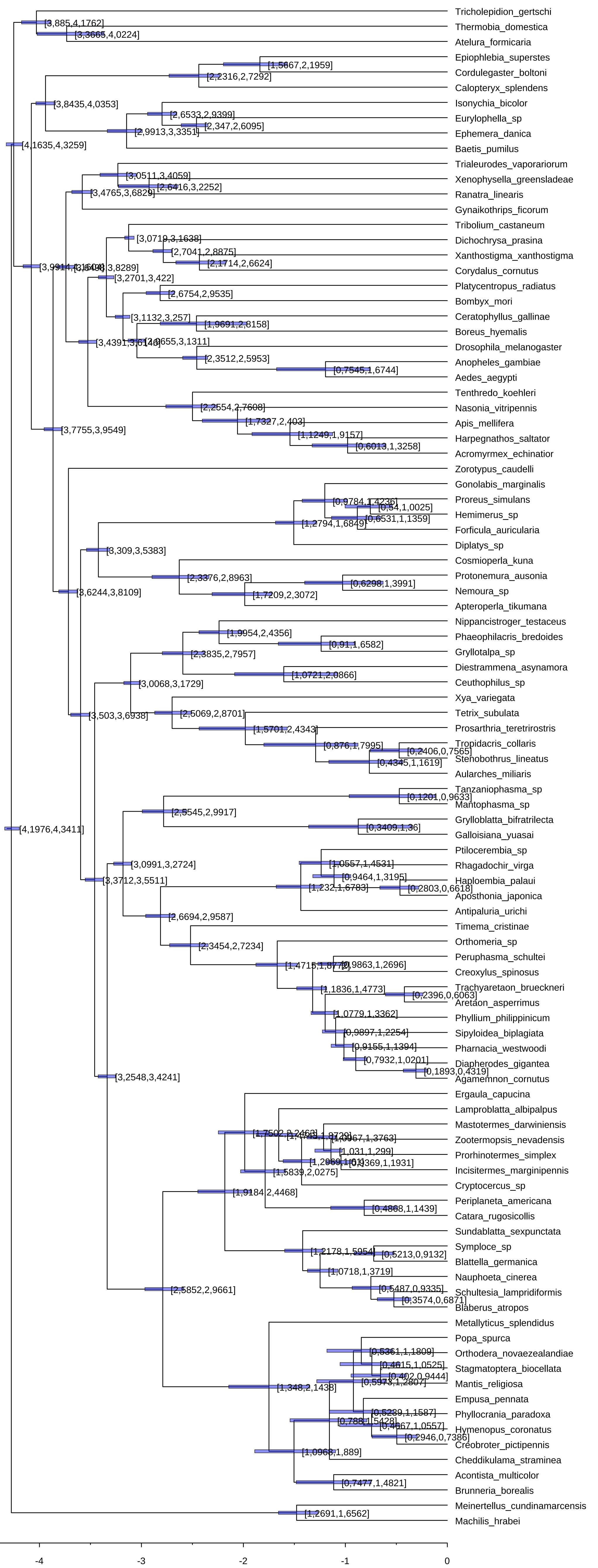

Tricholepidion_gertschi
Thermobia_domestica
Atelura_formicaria
Epiophlebia_superstes
Cordulegaster_boltoni
Calopteryx_splendens
Isonychia_bicolor
Eurylophella_sp
Ephemera_danica
Baetis_pumilus
Trialeurodes_vaporariorum
Xenophysella_greensladeae
Ranatra_linearis
Gynaikothrips_ficorum
Tribolium_castaneum
Dichochrysa_prasina
Xanthostigma_xanthostigma
Corydalus_cornutus
Platycentropus_radiatus
Bombyx_mori
Ceratophyllus_gallinae
Boreus_hyemalis
Drosophila_melanogaster
Anopheles_gambiae
Aedes_aegypti
Tenthredo_koehleri
Nasonia_vitripennis
Apis_mellifera
Harpegnathos_saltator
Acromyrmex_echinatior
Zorotypus_caudelli
Gonolabis_marginalis
Proreus_simulans
Hemimerus_sp
Forficula_auricularia
Diplatys_sp
Cosmioperla_kuna
Protonemura_ausonia
Nemoura_sp
Apteroperla_tikumana
Nippancistroger_testaceus
Phaeophilacris_bredoides
Gryllotalpa_sp
Diestrammena_asynamora
Ceuthophilus_sp
Xya_variegata
Tetrix_subulata
Prosarthria_teretrirostris
Tropidacris_collaris
Stenobothrus_lineatus
Aularches_miliaris
Tanzaniophasma_sp
Mantophasma_sp
Grylloblatta_bifratrilecta
Galloisiana_yuasai
Ptilocerembia_sp
Rhagadochir_virga
Haploembia_palaui
Aposthonia_japonica
Antipaluria_urichi
Timema_cristinae
Orthomeria_sp
Peruphasma_schultei
Creoxylus_spinosus
Trachyaretaon_brueckneri
Aretaon_asperrimus
Phyllium_philippinicum
Sipyloidea_biplagiata
Pharnacia_westwoodi
Diapherodes_gigantea
Agamemnon_cornutus
Ergaula_capucina
Lamproblatta_albipalpus
Mastotermes_darwiniensis
Zootermopsis_nevadensis
Prorhinotermes_simplex
Incisitermes_marginipennis
Cryptocercus_sp
Periplaneta_americana
Catara_rugosicollis
Sundablatta_sexpunctata
Symploce_sp
Blattella_germanica
Nauphoeta_cinerea
Schultesia_lampridiformis
Blaberus_atropos
Metallyticus_splendidus
Popa_spurca
Orthodera_novaezealandiae
Stagmatoptera_biocellata
Mantis_religiosa
Empusa_pennata
Phyllocrania_paradoxa
Hymenopus_coronatus
Creobroter_pictipennis
Cheddikulama_straminea
Acontista_multicolor
Brunneria_borealis
Meinertellus_cundinamarcensis
Machilis_hrabei
[3,885,4,1762]
[3,3665,4,0224]
[1,5667,2,1959]
[2,2316,2,7292]
[3,8435,4,0353]
[2,6533,2,9399]
[2,347,2,6095]
[2,9913,3,3351]
[4,1635,4,3259]
[3,0511,3,4059]
[2,6416,3,2252]
[3,4765,3,6829]
[3,0719,3,1638]
[2,7041,2,8875]
[2,1714,2,6624]
[3,2701,3,422]
[2,6754,2,9535]
[3,1132,3,257]
[1,9691,2,8158]
[2,3512,2,5953]
[0,7545,1,6744]
[2,2554,2,7608]
[1,7327,2,403]
[1,1249,1,9157]
[0,6013,1,3258]
[3,7755,3,9549]
[0,9784,1,4236]
[0,54,1,0025]
[0,6531,1,1359]
[1,2794,1,6849]
[3,309,3,5383]
[2,3376,2,8963]
[0,6298,1,3991]
[1,7209,2,3072]
[3,6244,3,8109]
[1,9954,2,4356]
[0,91,1,6582]
[2,3835,2,7957]
[1,0721,2,0866]
[3,0068,3,1729]
[3,503,3,6938]
[2,5069,2,8701]
[1,5701,2,4343]
[0,876,1,7995]
[0,2406,0,7565]
[0,4345,1,1619]
[0,1201,0,9633]
[2,5545,2,9917]
[0,3409,1,36]
[3,0991,3,2724]
[1,0557,1,4531]
[0,9464,1,3195]
[1,232,1,6783]
[0,2803,0,6618]
[3,3712,3,5511]
[2,6694,2,9587]
[2,3454,2,7234]
[0,9863,1,2696]
[1,1836,1,4773]
[0,2396,0,6063]
[1,0779,1,3362]
[0,9897,1,2254]
[0,9155,1,1394]
[0,7932,1,0201]
[0,1893,0,4319]
[3,2548,3,4241]
[1,031,1,299]
[0,1369,1,1931]
[1,5839,2,0275]
[1,9184,2,4468]
[0,4868,1,1439]
[1,2178,1,5954]
[0,5213,0,9132]
[1,0718,1,3719]
[0,5487,0,9335]
[0,3574,0,6871]
[2,5852,2,9661]
[0,5361,1,1809]
[0,4615,1,0525]
[0,402,0,9444]
[0,5973,1,2807]
[1,348,2,1438]
[0,5289,1,1587]
[0,4667,1,0557]
[0,2946,0,7386]
[1,0968,1,889]
[0,7477,1,4821]
[1,2691,1,6562]
[4,1976,4,3411]
-4
-3
-2
-1
0

**Figure S25.** Dated phylogeny of Polyneoptera, inferred with the independent rates (IR) clock model.

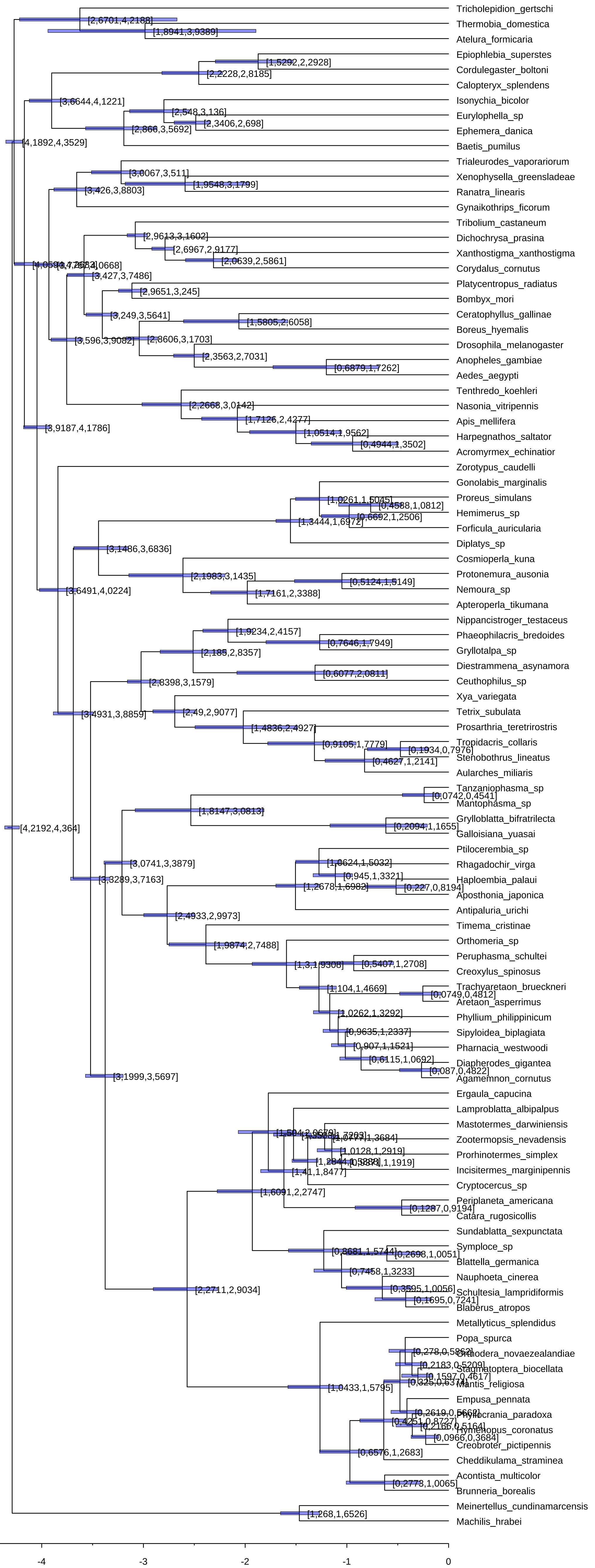

Tricholepidion_gertschi
Thermobia_domestica
Atelura_formicaria
Epiophlebia_superstes
Cordulegaster_boltoni
Calopteryx_splendens
Isonychia_bicolor
Eurylophella_sp
Ephemera_danica
Baetis_pumilus
Trialeurodes_vaporariorum
Xenophysella_greensladeae
Ranatra_linearis
Gynaikothrips_ficorum
Tribolium_castaneum
Dichochrysa_prasina
Xanthostigma_xanthostigma
Corydalus_cornutus
Platycentropus_radiatus
Bombyx_mori
Ceratophyllus_gallinae
Boreus_hyemalis
Drosophila_melanogaster
Anopheles_gambiae
Aedes_aegypti
Tenthredo_koehleri
Nasonia_vitripennis
Apis_mellifera
Harpegnathos_saltator
Acromyrmex_echinatior
Zorotypus_caudelli
Gonolabis_marginalis
Proreus_simulans
Hemimerus_sp
Forficula_auricularia
Diplatys_sp
Cosmioperla_kuna
Protonemura_ausonia
Nemoura_sp
Apteroperla_tikumana
Nippancistroger_testaceus
Phaeophilacris_bredoides
Gryllotalpa_sp
Diestrammena_asynamora
Ceuthophilus_sp
Xya_variegata
Tetrix_subulata
Prosarthria_teretrirostris
Tropidacris_collaris
Stenobothrus_lineatus
Aularches_miliaris
Tanzaniophasma_sp
Mantophasma_sp
Grylloblatta_bifratrilecta
Galloisiana_yuasai
Ptilocerembia_sp
Rhagadochir_virga
Haploembia_palaui
Aposthonia_japonica
Antipaluria_urichi
Timema_cristinae
Orthomeria_sp
Peruphasma_schultei
Creoxylus_spinosus
Trachyaretaon_brueckneri
Aretaon_asperrimus
Phyllium_philippinicum
Sipyloidea_biplagiata
Pharnacia_westwoodi
Diapherodes_gigantea
Agamemnon_cornutus
Ergaula_capucina
Lamproblatta_albipalpus
Mastotermes_darwiniensis
Zootermopsis_nevadensis
Prorhinotermes_simplex
Incisitermes_marginipennis
Cryptocercus_sp
Periplaneta_americana
Catara_rugosicollis
Sundablatta_sexpunctata
Symploce_sp
Blattella_germanica
Nauphoeta_cinerea
Schultesia_lampridiformis
Blaberus_atropos
Metallyticus_splendidus
Popa_spurca
Orthodera_novaezealandiae
Stagmatoptera_biocellata
Mantis_religiosa
Empusa_pennata
Phyllocrania_paradoxa
Hymenopus_coronatus
Creobroter_pictipennis
Cheddikulama_straminea
Acontista_multicolor
Brunneria_borealis
Meinertellus_cundinamarcensis
Machilis_hrabei
[2,6701,4,2188]
[1,8941,3,9389]
[1,5292,2,2928]
[2,2228,2,8185]
[3,6644,4,1221]
[2,548,3,136]
[2,3406,2,698]
[2,866,3,5692]
[4,1892,4,3529]
[3,0067,3,511]
[1,9548,3,1799]
[3,426,3,8803]
[2,9613,3,1602]
[2,6967,2,9177]
[2,0639,2,5861]
[3,427,3,7486]
[2,9651,3,245]
[3,249,3,5641]
[1,5805,2,6058]
[3,596,3,9082]
[2,8606,3,1703]
[2,3563,2,7031]
[0,6879,1,7262]
[2,2668,3,0142]
[1,7126,2,4277]
[1,0514,1,9562]
[0,4944,1,3502]
[3,9187,4,1786]
[1,0261,1,5045]
[0,4588,1,0812]
[0,6692,1,2506]
[1,3444,1,6972]
[3,1486,3,6836]
[2,1983,3,1435]
[0,5124,1,5149]
[1,7161,2,3388]
[3,6491,4,0224]
[1,9234,2,4157]
[0,7646,1,7949]
[2,185,2,8357]
[0,6077,2,0811]
[2,8398,3,1579]
[2,49,2,9077]
[3,4931,3,8859]
[1,4836,2,4927]
[0,9105,1,7779]
[0,1934,0,7976]
[0,4627,1,2141]
[0,0742,0,4541]
[1,8147,3,0813]
[0,2094,1,1655]
[4,2192,4,364]
[3,0741,3,3879]
[1,0624,1,5032]
[0,945,1,3321]
[3,3289,3,7163]
[1,2678,1,6982]
[0,227,0,8194]
[2,4933,2,9973]
[1,9874,2,7488]
[1,3,1,9308]
[0,5407,1,2708]
[1,104,1,4669]
[0,0749,0,4812]
[1,0262,1,3292]
[0,9635,1,2337]
[0,907,1,1521]
[0,6115,1,0692]
[0,087,0,4822]
[3,1999,3,5697]
[1,0777,1,3684]
[1,0128,1,2919]
[1,41,1,8477]
[1,6091,2,2747]
[0,1287,0,9194]
[0,8681,1,5744]
[0,2698,1,0051]
[0,7458,1,3233]
[0,3595,1,0056]
[0,1695,0,7241]
[2,2711,2,9034]
[0,278,0,5862]
[0,2183,0,5209]
[0,1597,0,4617]
[1,0433,1,5795]
[0,2619,0,5662]
[0,4251,0,8727]
[0,2166,0,5164]
[0,0966,0,3684]
[0,6576,1,2683]
[0,2773,1,0065]
[1,268,1,6526]
-4
-3
-2
-1
0

**Figure S26.** Phylogenetic tree of Polyneoptera displaying the effective priors.

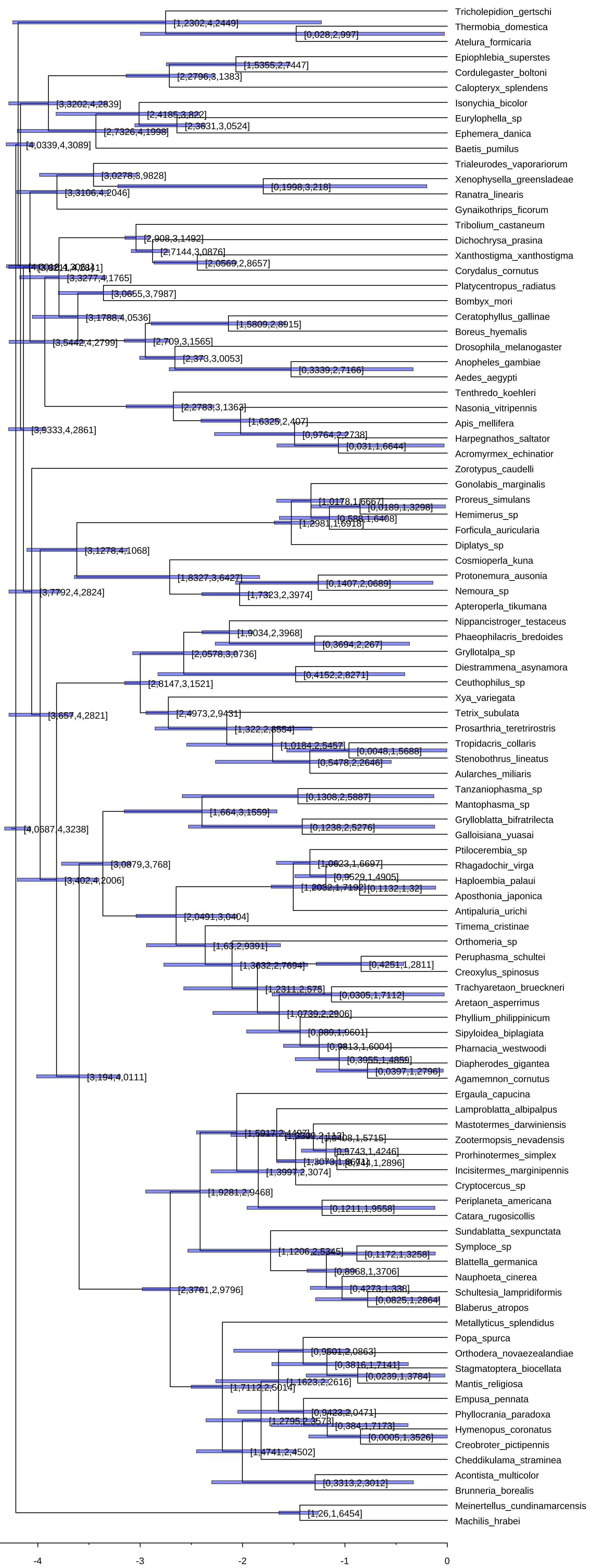
Tricholepidion_gertschi
Thermobia_domestica
Atelura_formicaria
Epiophlebia_superstes
Cordulegaster_boltoni
Calopteryx_splendens
Isonychia_bicolor
Eurylophella_sp
Ephemera_danica
Baetis_pumilus
Trialeurodes_vaporariorum
Xenophysella_greensladeae
Ranatra_linearis
Gynaikothrips_ficorum
Tribolium_castaneum
Dichochrysa_prasina
Xanthostigma_xanthostigma
Corydalus_cornutus
Platycentropus_radiatus
Bombyx_mori
Ceratophyllus_gallinae
Boreus_hyemalis
Drosophila_melanogaster
Anopheles_gambiae
Aedes_aegypti
Tenthredo_koehleri
Nasonia_vitripennis
Apis_mellifera
Harpegnathos_saltator
Acromyrmex_echinatior
Zorotypus_caudelli
Gonolabis_marginalis
Proreus_simulans
Hemimerus_sp
Forficula_auricularia
Diplatys_sp
Cosmioperla_kuna
Protonemura_ausonia
Nemoura_sp
Apteroperla_tikumana
Nippancistroger_testaceus
Phaeophilacris_bredoides
Gryllotalpa_sp
Diestrammena_asynamora
Ceuthophilus_sp
Xya_variegata
Tetrix_subulata
Prosarthria_teretrirostris
Tropidacris_collaris
Stenobothrus_lineatus
Aularches_miliaris
Tanzaniophasma_sp
Mantophasma_sp
Grylloblatta_bifratrilecta
Galloisiana_yuasai
Ptiloceembia_sp
Rhagadochir_virga
Haploembia_palaui
Aposthonia_japonica
Antipaluria_urichi
Timema_cristinae
Orthomeria_sp
Peruphasma_schultei
Creoxylus_spinosus
Trachyaretaon_brueckneri
Aretaon_asperrimus
Phyllium_philippinicum
Sipyloidea_biplagiata
Pharnacia_westwoodi
Diapherodes_gigantea
Agamemnon_cornutus
Ergaula_capucina
Lamproblatta_albipalpus
Mastotermes_darwiniensis
Zootermopsis_nevadensis
Prorhinotermes_simplex
Incisitermes_marginipennis
Cryptocercus_sp
Periplaneta_americana
Catara_rugosicollis
Sundablatta_sexpunctata
Symploce_sp
Blattella_germanica
Nauphoeta_cinerea
Schultesia_lampridiformis
Blaberus_atropos
Metallyticus_splendidus
Popa_spurca
Orthodera_novaezealandiae
Stagmatoptera_biocellata
Mantis_religiosa
Empusa_pennata
Phyllocrania_paradoxa
Hymenopus_coronatus
Creobroter_pictipennis
Cheddikulama_straminea
Acontista_multicolor
Brunneria_borealis
Meinertellus_cundinamarcensis
Machilis_hrabei
[1,2302,4,2449]
[0,028,2,997]
[1,5355,2,7447]
[2,2796,3,1383]
[3,3202,4,2839]
[2,3631,3,0524]
[2,7326,4,1998]
[4,0339,4,3089]
[3,0278,3,9828]
[0,1998,3,218]
[3,3106,4,2046]
[2,908,3,1492]
[2,7144,3,0876]
[2,0569,2,8657]
[3,3277,4,1765]
[3,0655,3,7987]
[3,1788,4,0536]
[1,5809,2,8915]
[3,5442,4,2799]
[2,709,3,1565]
[2,373,3,0053]
[0,3339,2,7166]
[2,2783,3,1363]
[1,6325,2,407]
[0,9764,2,2738]
[0,031,1,6644]
[3,9333,4,2861]
[1,0178,1,6667]
[0,0189,1,3298]
[0,588,1,6408]
[1,2981,1,6918]
[3,1278,4,1068]
[1,8327,3,6427]
[0,1407,2,0689]
[1,7323,2,3974]
[3,7792,4,2824]
[1,9034,2,3968]
[0,3694,2,267]
[2,0578,3,0736]
[0,4152,2,8271]
[2,8147,3,1521]
[2,4973,2,9431]
[1,322,2,8554]
[1,0184,2,5457]
[0,0048,1,5688]
[0,5478,2,2646]
[3,657,4,2821]
[0,1308,2,5887]
[1,664,3,1559]
[0,1238,2,5276]
[4,0687,4,3238]
[3,0879,3,768]
[1,0623,1,6697]
[0,9529,1,4905]
[0,1132,1,32]
[3,402,4,2006]
[2,0491,3,0404]
[1,63,2,9391]
[1,3632,2,7694]
[0,4251,1,2811]
[1,2311,2,575]
[0,0305,1,7112]
[1,0739,2,2906]
[0,989,1,9601]
[0,9813,1,6004]
[0,3955,1,4859]
[0,0397,1,2796]
[3,194,4,0111]
[1,0408,1,5715]
[0,9743,1,4246]
[1,3997,2,3074]
[1,9281,2,9468]
[0,1211,1,9558]
[1,1206,2,5345]
[0,1172,1,3258]
[0,8968,1,3706]
[0,4273,1,338]
[0,0825,1,2864]
[2,3761,2,9796]
[0,9501,2,0863]
[0,3816,1,7141]
[0,0239,1,3784]
[1,1623,2,2616]
[1,7112,2,5014]
[0,9423,2,0471]
[1,2795,2,3573]
[0,384,1,7173]
[0,0005,1,3526]
[1,4741,2,4502]
[0,3313,2,3012]
[1,26,1,6454]
-4
-3
-2
-1
0

**Figure S27.** Phylogeny of Zoraptera inferred with the site-heterogeneous model CAT-GTR+G implemented in PhyloBayes, on the 5-gene nucleotide dataset.

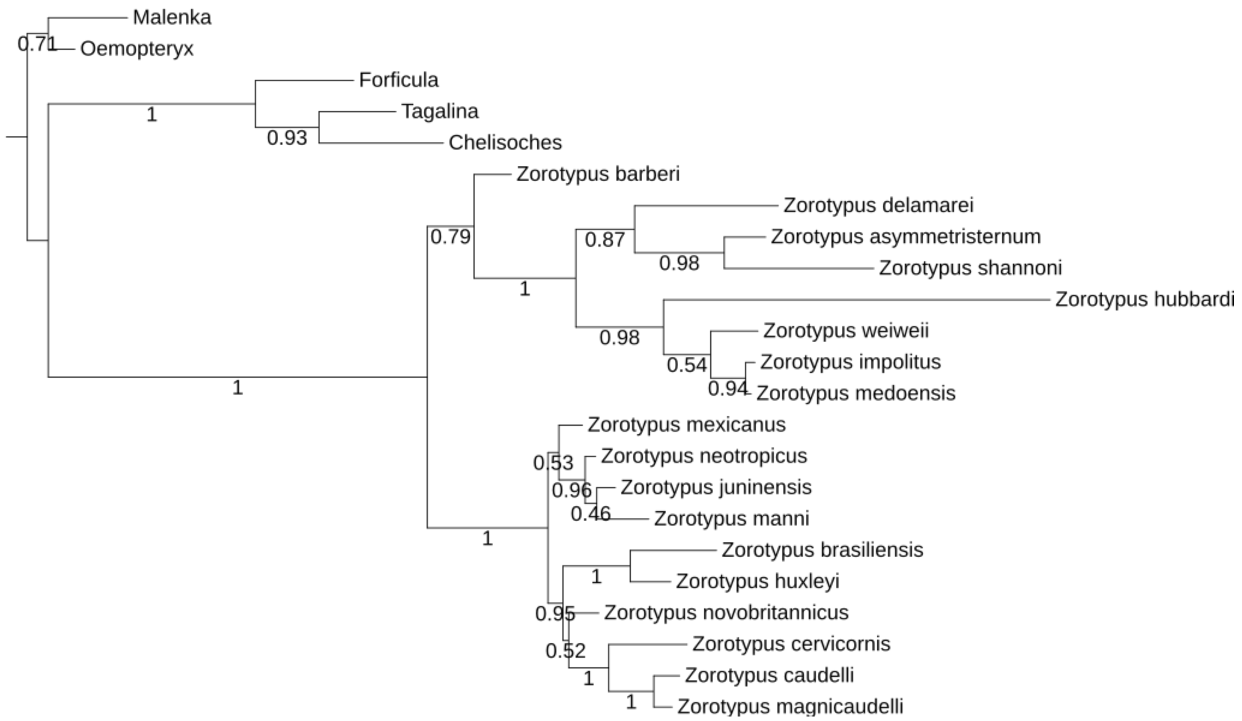

**Figure S28.** Phylogeny of Zoraptera inferred with the GTR+F+R3 model implemented in IQ-TREE, on the 5-gene nucleotide dataset.

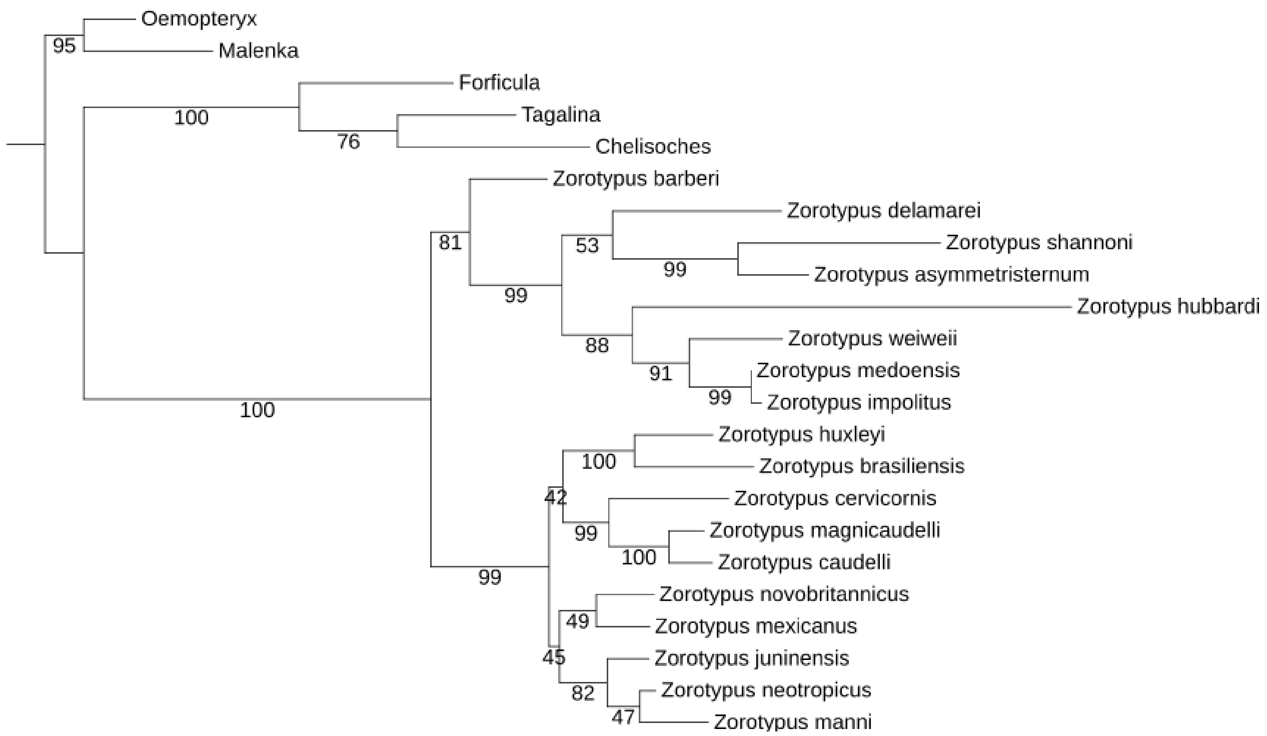

**Figure S29.** Phylogeny of Zoraptera inferred with the TVM+F+I+G4 model implemented in IQ-TREE, on the 5-gene nucleotide dataset.

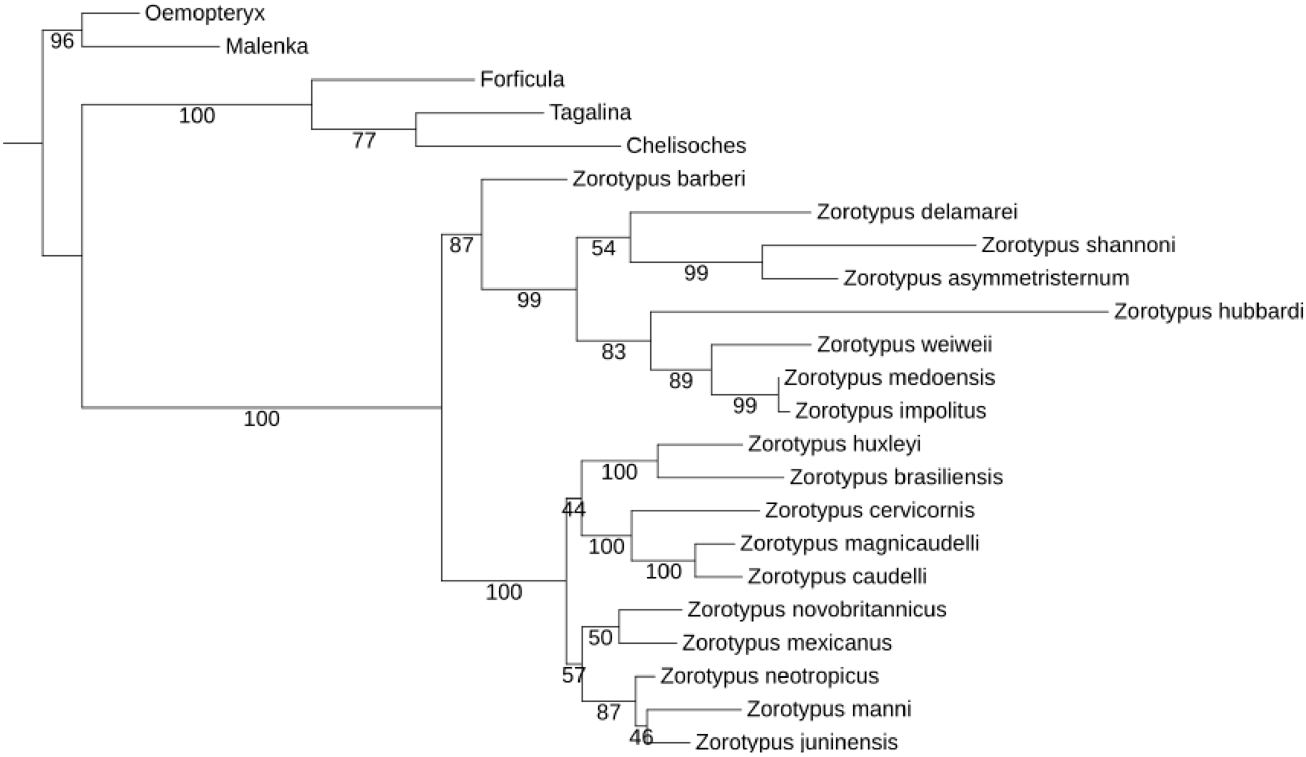

**Figure S30.** Phylogeny of Zoraptera inferred with the JC model implemented in IQ-TREE, on the 5-gene nucleotide dataset.

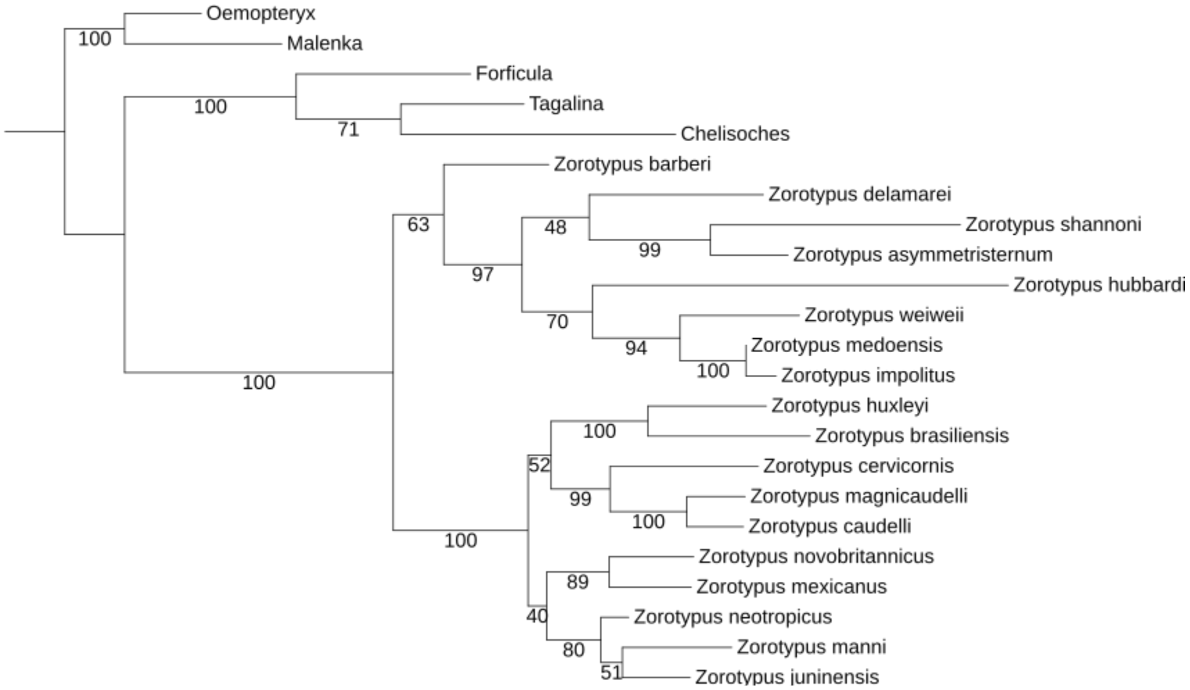

**Figure S31.** Phylogeny of Zoraptera inferred with maximum parsimony analysis of a 12-character matrix in TNT, with character states mapped.

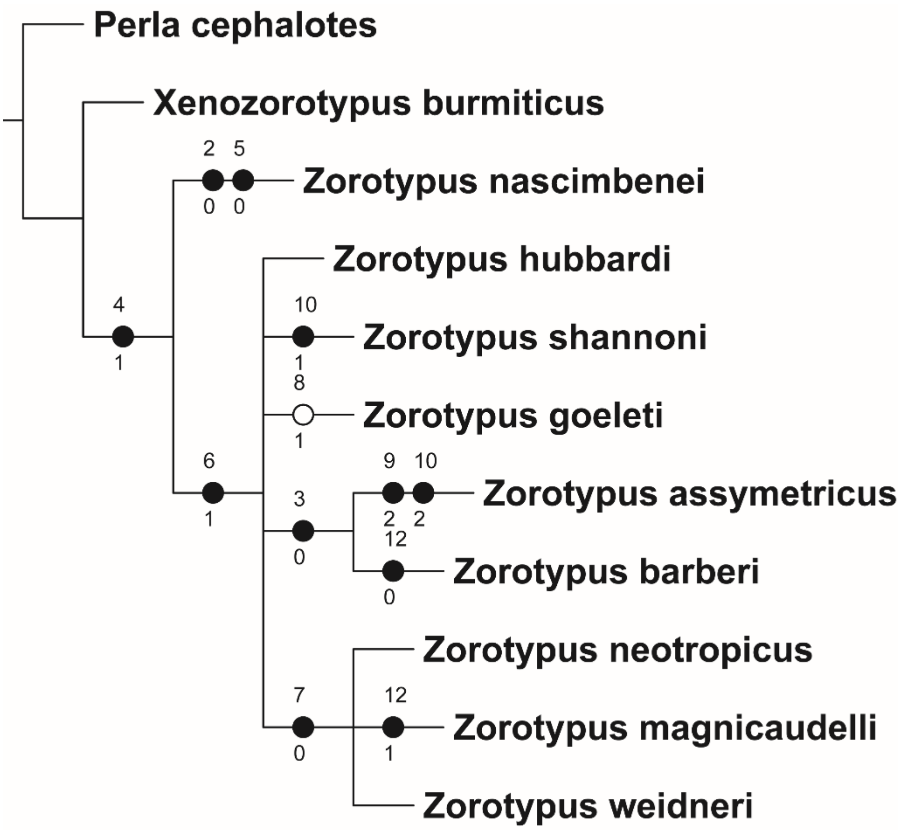

#### Supplementary Tables

**Table S1.** Comparison of historically proposed hypotheses of Zoraptera placement with approximately unbiased (AU) tests. *P*-value > 0.05: topology not rejected; *P*-value < 0.05: topology rejected significantly (*); *P*-value = 0: topology rejected with high significance (**).

| **Topology** | $\boldsymbol{P}_{\mathbf{AU}}$ |
|---|---|
| Zoraptera sister to remaining Polyneoptera, (Dermaptera + Plecoptera) forming the next branch | 0.995 |
| Zoraptera + Dermaptera sister to remaining Polyneoptera, Plecoptera forming the next branch | 0.006* |
| (Zoraptera (Dermaptera + Plecoptera) clade sister to remaining Polyneoptera | 0.000** |
| Zoraptera + Plecoptera sister to remaining Polyneoptera, Dermaptera forming the next branch | 0.000** |
| Dermaptera sister to remaining Polyneoptera, Zoraptera forming the next branch, then Plecoptera | 0.000** |
| Dermaptera sister to remaining Polyneoptera, Plecoptera forming the next branch, Zoraptera sister to Dictyoptera | 0.000** |
| Dermaptera sister to remaining Polyneoptera, Plecoptera forming the next branch, Zoraptera sister to Blattodea | 0.000** |
| Dermaptera sister to remaining Polyneoptera, Plecoptera forming the next branch, Zoraptera sister to Isoptera | 0.000** |
| Dermaptera sister to remaining Polyneoptera, Plecoptera forming the next branch, Zoraptera sister to Embioptera | 0.000** |
| Dermaptera sister to remaining Polyneoptera Plecoptera forming the next branch, Zoraptera sister to Eukinolabia | 0.000** |
| Dermaptera + Plecoptera sister to remaining Polyneoptera, Zoraptera sister to Dictyoptera | 0.000** |
| Dermaptera + Plecoptera sister to remaining Polyneoptera, Zoraptera sister to Blattodea | 0.000** |
| Dermaptera + Plecoptera sister to remaining Polyneoptera, Zoraptera sister to Isoptera | 0.000** |
| Dermaptera + Plecoptera sister to remaining Polyneoptera, Zoraptera sister to Embioptera | 0.000** |
| Dermaptera + Plecoptera sister to remaining Polyneoptera, Zoraptera sister to Eukinolabia | 0.000** |
| Zoraptera sister to Thysanoptera | 0.000** |
| Zoraptera sister to Eumetabola | 0.000** |
| Zoraptera sister to Holometabola | 0.000** |
| Plecoptera sister to remaining Neoptera | 0.000** |
| Dermaptera sister to Eukinolabia | 0.000** |

**Table S2.** Accession numbers used for the total evidence phylogenetic reconstruction of Zoraptera, based mainly on Matsumura et al. [279] and Kočárek et al. [1].

| Taxon | 18S | H3 | 12S | 16S | COI |
|---|---|---|---|---|---|
| ZORAPTERA | | | | | |
| *Z. asymmetristernum* | LC477109 | | | LC476756 | MN790617 |
| *Z. barberi* | LC477110 | | | LC476757 | |
| *Z. brasiliensis* | LC477104 | LC477128 | LC471603 | LC476748 | MN790624 |
| *Z. caudelli* | LC477097 | LC477120 | LC471596 | LC476741 | |
| *Z. cervicornis* | LC477099 | LC477122 | LC471598 | LC476743 | MN790626 |
| *Z. delamarei* | MN790597 | | | MN790583 | MN790618 |
| *Z. hubbardi* | LC477087 | AY521734 | LC471587 | LC476731 | |
| *Z. huxleyi* | LC477105 | LC477129 | LC471604 | LC476749 | |
| *Z. impolitus* | LC477088 | LC477112 | LC471588 | LC476732 | |
| *Z. juninensis* | LC477092 | LC477115 | LC471592 | LC476736 | |
| *Z. magnicaudelli* | LC477096 | LC477119 | LC471595 | LC476740 | MN790627 |
| *Z. manni* | LC477094 | LC477117 | | LC476738 | |
| *Z. medoensis* | KM246626 | | | JQ910991 | KJ467512 |
| *Z. mexicanus* | | LC477133 | | LC476753 | |
| *Z. neotropicus* | MN790606 | | | MN790584 | MN790619 |
| *Z. novobritannicus* | LC477100 | LC477123 | LC471599 | EF623273 | |
| *Z. shannoni* | LC477089 | | LC471589 | LC476733 | |
| *Z. weiweii* | MN790602 | | | MN790591 | MN790622 |
| DERMAPTERA | | | | | |
| *Chelisoches* sp. | AH012713 | AY707426 | EF623305 | EF623146 | |
| *Forficula* sp. | MN790613 | AY521703 | KF855790 | EU863268 | HM376337 |
| *Tagalina* sp. | AY521838 | AY521704 | | | KX069080 |
| PLECOPTERA | | | | | |
| *Malenka* sp. | AY338724 | | EU054931 | EF623182 | KM537296 |
| *Oemopteryx* sp. | AY521879 | AY521725 | EF623432 | EF623266 | HQ969542 |

**Tab. S3.** Fit of maximum likelihood (ML) models to the zorapteran molecular dataset analysed in ModelFinder. Abbreviations: AICc, corrected Akaike information criterion scores; BIC, Bayesian Information Criterion.

| **Model** | **AICc** | **BIC** |
|---|---|---|
| JC | 23033.144 | 23264.629 |
| JC+I | 21694.439 | 21931.253 |
| JC+G4 | 21561.185 | 21797.999 |
| JC+I+G4 | 21550.133 | 21792.273 |
| JC+R2 | 21584.099 | 21826.239 |
| JC+R3 | 21551.182 | 21803.967 |
| JC+R4 | 21552.288 | 21815.707 |
| F81+F | 22935.069 | 23182.533 |
| F81+F+I | 21572.142 | 21824.927 |
| F81+F+G4 | 21434.861 | 21687.645 |
| F81+F+I+G4 | 21415.983 | 21674.087 |
| F81+F+R2 | 21460.988 | 21719.091 |
| F81+F+R3 | 21419.083 | 21687.816 |
| F81+F+R4 | 21419.112 | 21698.463 |
| K2P | 22719.590 | 22956.404 |
| K2P+I | 21351.849 | 21593.989 |
| K2P+G4 | 21198.835 | 21440.975 |
| K2P+I+G4 | 21178.705 | 21426.169 |
| K2P+R2 | 21227.764 | 21475.227 |
| K2P+R3 | 21175.317 | 21433.420 |
| K2P+R4 | 21178.374 | 21447.107 |
| HKY+F | 22607.117 | 22859.902 |
| HKY+F+I | 21194.545 | 21452.649 |
| HKY+F+G4 | 21034.242 | 21292.345 |
| HKY+F+I+G4 | 21005.839 | 21269.258 |
| HKY+F+R2 | 21068.573 | 21331.993 |
| HKY+F+R3 | 20996.974 | 21271.017 |
| HKY+F+R4 | 21000.033 | 21284.690 |
| TNe | 22718.200 | 22960.340 |
| TNe+I | 21353.962 | 21601.426 |
| TNe+G4 | 21200.927 | 21448.390 |
| TNe+I+G4 | 21180.060 | 21432.845 |
| TNe+R2 | 21229.818 | 21482.603 |
| TNe+R3 | 21176.772 | 21440.191 |
| TNe+R4 | 21179.886 | 21453.930 |
| TN+F | 22597.044 | 22855.147 |
| TN+F+I | 21194.106 | 21457.526 |
| TN+F+G4 | 21033.824 | 21297.244 |
| TN+F+I+G4 | 21004.475 | 21273.207 |
| TN+F+R2 | 21068.879 | 21337.611 |
| TN+F+R3 | 20996.032 | 21275.384 |
| TN+F+R4 | 20998.866 | 21288.826 |
| K3P | 22688.293 | 22930.434 |
| K3P+I | 21315.139 | 21562.603 |
| K3P+G4 | 21165.176 | 21412.640 |

| | | |
|---|---|---|
| K3P+I+G4 | 21143.593 | 21396.377 |
| K3P+R2 | 21192.869 | 21445.653 |
| K3P+R3 | 21140.164 | 21403.583 |
| K3P+R4 | 21143.507 | 21417.550 |
| K3Pu+F | 22582.353 | 22840.457 |
| K3Pu+F+I | 21170.575 | 21433.995 |
| K3Pu+F+G4 | 21013.429 | 21276.848 |
| K3Pu+F+I+G4 | 20984.552 | 21253.285 |
| K3Pu+F+R2 | 21046.042 | 21314.774 |
| K3Pu+F+R3 | 20975.843 | 21255.194 |
| K3Pu+F+R4 | 20978.935 | 21268.895 |
| TPM2+F | 22556.938 | 22815.041 |
| TPM2+F+I | 21167.226 | 21430.645 |
| TPM2+F+G4 | 21006.682 | 21270.101 |
| TPM2+F+I+G4 | 20979.518 | 21248.251 |
| TPM2+F+R2 | 21040.595 | 21309.328 |
| TPM2+F+R3 | 20971.685 | 21251.037 |
| TPM2+F+R4 | 20974.303 | 21264.263 |
| TPM2u+F | 22556.938 | 22815.041 |
| TPM2u+F+I | 21167.225 | 21430.644 |
| TPM2u+F+G4 | 21006.682 | 21270.101 |
| TPM2u+F+I+G4 | 20979.447 | 21248.179 |
| TPM2u+F+R2 | 21040.596 | 21309.329 |
| TPM2u+F+R3 | 20971.687 | 21251.038 |
| TPM2u+F+R4 | 20974.249 | 21264.209 |
| TPM3+F | 22478.334 | 22736.437 |
| TPM3+F+I | 21082.729 | 21346.148 |
| TPM3+F+G4 | 20919.969 | 21183.388 |
| TPM3+F+I+G4 | 20897.473 | 21166.206 |
| TPM3+F+R2 | 20950.089 | 21218.822 |
| TPM3+F+R3 | 20895.545 | 21174.896 |
| TPM3+F+R4 | 20898.709 | 21188.668 |
| TPM3u+F | 22478.333 | 22736.437 |
| TPM3u+F+I | 21082.729 | 21346.148 |
| TPM3u+F+G4 | 20919.968 | 21183.387 |
| TPM3u+F+I+G4 | 20897.440 | 21166.172 |
| TPM3u+F+R2 | 20950.085 | 21218.818 |
| TPM3u+F+R3 | 20895.523 | 21174.874 |
| TPM3u+F+R4 | 20898.659 | 21188.618 |
| TIMe | 22686.898 | 22934.362 |
| TIMe+I | 21317.254 | 21570.039 |
| TIMe+G4 | 21167.261 | 21420.046 |
| TIMe+I+G4 | 21145.183 | 21403.287 |
| TIMe+R2 | 21194.925 | 21453.028 |
| TIMe+R3 | 21142.277 | 21411.010 |
| TIMe+R4 | 21145.475 | 21424.827 |
| TIM+F | 22572.269 | 22835.688 |
| TIM+F+I | 21170.129 | 21438.862 |
| TIM+F+G4 | 21013.197 | 21281.929 |
| TIM+F+I+G4 | 20983.293 | 21257.336 |
| TIM+F+R2 | 21046.440 | 21320.483 |

| | | |
|---|---|---|
| TIM+F+R3 | 20975.121 | 21259.778 |
| TIM+F+R4 | 20977.990 | 21273.251 |
| TIM2e | 22646.998 | 22894.461 |
| TIM2e+I | 21308.506 | 21561.291 |
| TIM2e+G4 | 21151.966 | 21404.751 |
| TIM2e+I+G4 | 21131.897 | 21390.001 |
| TIM2e+R2 | 21181.401 | 21439.505 |
| TIM2e+R3 | 21129.201 | 21397.934 |
| TIM2e+R4 | 21131.683 | 21411.034 |
| TIM2+F | 22547.472 | 22810.892 |
| TIM2+F+I | 21166.740 | 21435.473 |
| TIM2+F+G4 | 21006.313 | 21275.046 |
| TIM2+F+I+G4 | 20978.380 | 21252.423 |
| TIM2+F+R2 | 21040.965 | 21315.008 |
| TIM2+F+R3 | 20970.928 | 21255.585 |
| TIM2+F+R4 | 20973.064 | 21268.324 |
| TIM3e | 22512.225 | 22759.689 |
| TIM3e+I | 21171.910 | 21424.695 |
| TIM3e+G4 | 21002.450 | 21255.235 |
| TIM3e+I+G4 | 20985.051 | 21243.154 |
| TIM3e+R2 | 21033.251 | 21291.355 |
| TIM3e+R3 | 20987.801 | 21256.533 |
| TIM3e+R4 | 20990.976 | 21270.327 |
| TIM3+F | 22466.438 | 22729.857 |
| TIM3+F+I | 21081.659 | 21350.392 |
| TIM3+F+G4 | 20920.167 | 21188.899 |
| TIM3+F+I+G4 | 20897.510 | 21171.553 |
| TIM3+F+R2 | 20950.200 | 21224.244 |
| TIM3+F+R3 | 20895.624 | 21180.281 |
| TIM3+F+R4 | 20898.454 | 21193.714 |
| TVMe | 22444.109 | 22696.894 |
| TVMe+I | 21120.817 | 21378.920 |
| TVMe+G4 | 20950.174 | 21208.277 |
| TVMe+I+G4 | 20933.210 | 21196.629 |
| TVMe+R2 | 20981.142 | 21244.561 |
| TVMe+R3 | 20935.465 | 21209.508 |
| TVMe+R4 | 20938.861 | 21223.518 |
| TVM+F | 22430.368 | 22699.100 |
| TVM+F+I | 21056.389 | 21330.432 |
| TVM+F+G4 | 20894.016 | 21168.059 |
| TVM+F+I+G4 | 20872.673 | 21152.024 |
| TVM+F+R2 | 20923.806 | 21203.158 |
| TVM+F+R3 | 20871.157 | 21161.117 |
| TVM+F+R4 | 20874.035 | 21174.593 |
| SYM | 22441.885 | 22699.988 |
| SYM+I | 21122.896 | 21386.315 |
| SYM+G4 | 20952.255 | 21215.674 |
| SYM+I+G4 | 20935.319 | 21204.051 |
| SYM+R2 | 20983.257 | 21251.989 |
| SYM+R3 | 20937.511 | 21216.863 |
| SYM+R4 | 20940.955 | 21230.915 |

| | | |
|---|---|---|
| GTR+F | 22419.105 | 22693.148 |
| GTR+F+I | 21055.265 | 21334.616 |
| GTR+F+G4 | 20894.254 | 21173.606 |
| GTR+F+I+G4 | 20872.803 | 21157.460 |
| GTR+F+R2 | 20923.995 | 21208.652 |
| GTR+F+R3 | 20871.145 | 21166.405 |
| GTR+F+R4 | 20874.041 | 21179.894 |

**Tab. S4.** List of characters used to evaluate the intraordinal relationships within Zoraptera.

| # | **Character** | **States** |
|---|---|---|
| 1 | Orientation of head | (0) prognathous; (1) orthognathous |
| 2 | Number of antennomeres | (0) eight; (1) nine; (2) more than nine |
| 3 | Males' vertex. | (0) with a hairy patch; (1) without a hairy patch |
| 4 | Wing vein $M_{3+4}$ | (0) present; (1) absent |
| 5 | Jugal setae along the middle the third of the posterior forewing margin | (0) present; (1) absent |
| 6 | Empodium of the meta-pretarsus | (0) pronounced and slightly expanded; (1) reduced to a single, slender seta or absent altogether |
| 7 | Metafemoral spines | (0) occurring at regular intervals; (1) basal spine or spines separated by a distinctive gap (0) |
| 8 | Cerci segmentation | (0) unsegmented; (1) two-segmented; (2) multisegmented |
| 9 | Male abdominal tergite | (0) with a small mating hook; (1) with a long mating hook; (2) mating hook with two long prolongations; (3) mating hook absent |
| 10 | Subgenital plate | (0) unmodified; (1) bifurcated; (2) depressed; (3) projections |
| 11 | Male genitalia | (0) symmetrical; (1) asymmetrical |
| 12 | Intromittent organ | (0) elongated, straight; (1) elongated, spiral; (2) not elongated |

**Tab. S5.** Character matrix for evaluating the intraordinal relationships within Zoraptera.

| **Taxon** | **Matrix** |
|---|---|
| *Xenozorotypus burmiticus* | 11101010???? |
| *Zorotypus nascimbenei* | 10110010???? |
| *Zorotypus hubbardi* | 111111100012 |
| *Zorotypus shannoni* | 111111101112 |
| *Zorotypus assymetricus* | 110111102212 |
| *Zorotypus neotropicus* | 11?11100???? |
| *Zorotypus magnicaudelli* | 11?111000001 |
| *Zorotypus weidneri* | 111?11001002 |
| *Zorotypus barberi* | 110111100000 |
| *Zorotypus goeleti* | 11?11111???? |
| *Perla cephalotes* | 021010-2-002 |
